# Identification of the intracellular protein targets of a bio-active clickable half-sandwich iridium complex by chemical proteomics

**DOI:** 10.1101/2023.05.24.542041

**Authors:** Robin Ramos, Anthi Karaiskou, Candice Botuha, Michaël Trichet, Florent Dingli, Jérémy Forté, France Lam, Alexis Canette, Chloé Chaumeton, Murielle Salome, Thomas Chenuel, Céline Bergonzi, Philippe Meyer, Sylvain Bohic, Damarys Loew, Michèle Salmain, Joëlle Sobczak-Thépot

**Affiliations:** Sorbonne Université, INSERM, Centre de Recherche Saint Antoine, 184 rue du Faubourg Saint Antoine, F-75012 Paris, France.; Sorbonne Université, CNRS, Institut Parisien de Chimie Moléculaire, 4 place Jussieu, F75005 Paris, France.; Sorbonne Université, CNRS, Institut de Biologie Paris-Seine, Service d’imagerie cellulaire, F-75005, Paris; Institut Curie, PSL Research University, CurieCoreTech Mass Spectrometry Proteomics, F-75248 Paris, France; ESRF, The European Synchrotron, Grenoble, France; Sorbonne Université, PSL, CNRS, UMR8226, Institut de Biologie Physico-Chimique, Laboratoire de Biologie Moléculaire et Cellulaire des Eucaryotes, 75005 Paris, France.; Université Grenoble Alpes, INSERM, UA7 STROBE, Synchrotron Radiation for Biomedicine, Grenoble, France

## Abstract

Identification of intracellular targets of anticancer drug candidates provides key information on their mechanism of action. Exploiting the ability of the anticancer (C^N)-chelated half-sandwich iridium(III) complexes to covalently bind proteins, click chemistry with a bioorthogonal azido probe was used to localize a phenyloxazoline-chelated iridium complex within cells and profile its interactome at the proteome-wide scale. Proteins involved in protein folding and actin cytoskeleton regulation were identified as high affinity targets. Upon iridium complex treatment, HSP90 folding activity was inhibited *in vitro* and major cytoskeleton disorganization was observed. We used a wide array of imaging and biochemical methods to validate selected targets and obtain a multiscale overview of the effects of this complex on live human cells. We demonstrate that it behaves as a dual agent, inducing both electrophilic and oxidative stresses in cells that account for its cytotoxicity.

## Introduction

Uncovering metal-based alternatives to platinum anticancer drugs is an active research area as oncologists are still in need of molecules with an original mode of action and less side effects. Iridium(III) complexes are actively investigated to this purpose^1^ with two main families of molecules under current study, namely the kinetically inert photoluminescent bis-cyclometallated complexes^2–4^ and the so-called half-sandwich complexes comprising a penta-substituted cyclopentadienyl ligand. The coordination sphere of this latter class of compounds is generally completed by a bidentate chelating ligand and an exchangeable halogeno ligand^5^.

The first cytotoxic half-sandwich iridium(III) complexes have been introduced ten years ago^6, 7^ and have since continuously attracted interest, mostly owing to a mode of action different from platinum complexes. These antiproliferative properties were demonstrated on cancer cell cultures including platinum-resistant cell lines, as well as on solid tumors^8–11^. Most complexes reported in the literature are known to increase reactive oxygen species (ROS) levels in cells^12–15^ and induce mitochondrial uncoupling^16^, triggering apoptotic cell death. Consistently, upregulation of the antioxidant response was demonstrated in a pharmaco-genomic study realized in ovarian cells treated with a highly active azopyridine iridium complex^17^.

Importantly, the lability of the halogeno ligand of some half-sandwich iridium complexes provides them with a reactivity towards various cellular substrates. The first recognized cellular substrate was NADH, whose catalytic oxidation is responsible for the production of H_2_O_2_ from O_2_ by hydride transfer^18^ demonstrated both *in vitro* and *in vivo*^12, 14^. While other potential cellular targets were investigated, such as the tripeptide GSH (via its central cysteine residue)^19^, calmodulin (via some of its methionines)^20^ and the redox enzyme thioredoxin reductase^13^, no general approach has been undertaken to allow the unbiased identification of the targets of half-sandwich iridium complexes *in vivo*.

We previously introduced a series of ten half-sandwich complexes including diversely substituted phenyloxazoline (phox) chelating ligands. From the subsequent structure-activity relationship (SAR) study, **Ir2** carrying a dimethyl substituent on the oxazoline ring was identified as the best hit amongst the series in terms of overall cytotoxicity, inhibition of proliferation, apoptosis induction and H_2_O_2_ production, both *in vitro* and *in vivo*^1^^4^.

We also designed a half-sandwich iridium complex comprising a modified phenyloxazoline ligand to which was appended a BODIPY entity, so as to address the drug to membrane-rich organelles^21^. Thanks to its fluorescent properties, the complex was localized in the endoplasmic reticulum and mitochondrial membranes. Most importantly, we could demonstrate that this complex forms covalent adducts with a subset of intracellular proteins via a ligand exchange mechanism. The ability of phenyloxazoline-chelated half-sandwich iridium complexes to both bind covalently to proteins and to generate oxidative stress prompted us to develop new strategies to elucidate their mechanism of action, to identify their intracellular protein targets and investigate their relevance in terms of protein function and cellular behavior.

In recent years, metalloproteomics has emerged as a powerful tool to identify the cellular protein targets of various anticancer metal-based drugs^22–24^. Different workflows have been designed, the most advanced ones relying on bioorthogonal probes, combined with protein pull-down, shotgun LC-MS/MS analysis and database interrogation for protein identification. The first step of this workflow can be performed either with cell lysates^25–29^ or with live cells^30^. This latter strategy has been applied to uncover the main protein targets of inert gold(III)^31, 32^ and bis-cyclometallated iridium(III)^33^ complexes using photoaffinity probes, but also of arsenic trioxide^34, 35^ in cancer cells and cisplatin in yeast^36^ using azido derivatives.

Herein we report the identification of **Ir2** main protein targets using an insightful and unbiased chemoproteomics approach. Two half-sandwich iridium probes **IrN_3_** and **Ir2N_3_** were synthesized (**Figure 1a**), comprising a bioorthogonal and minimally invasive azido function, ready to undergo chemoselective (“click”) ligations with relevant partners. For this, the well-known copper-catalyzed [3+2] azide-alkyne cycloaddition (CuAAC) on the iridium probes was performed for fluorescence imaging and for partner protein enrichment followed by identification using quantitative mass spectrometry (**Figure 1b**). In combination with an array of approaches including multiscale cell imaging and *in vitro* assays, this study uncovers the main cellular processes that are altered by this class of complexes, providing molecular bases for their cytotoxicity.

**Figure 1.**
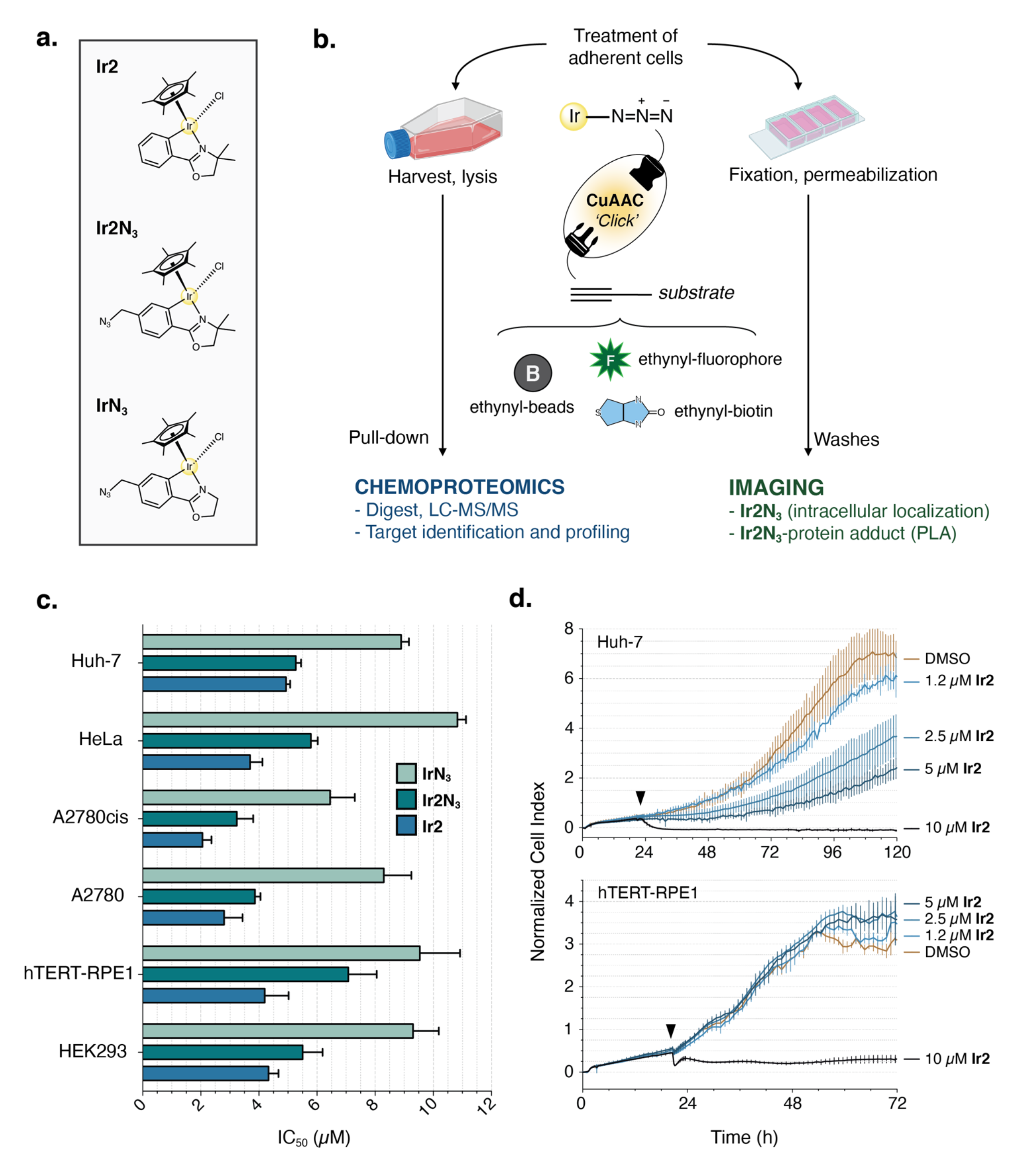
Experiment design and inhibition of proliferation on a panel of cell lines **a.** Half-sandwich complexes used in this study (**Ir2**, **IrN_3_**, **Ir2N_3_**). **b.** Experimental strategy using CuAAC (CuSO_4_, THPTA, sodium ascorbate) with ethynyl-substituted substrates to perform chemoproteomics and cell imaging (created with BioRender.com). **c.** Histograms representing the IC_50_ of the three complexes in various cell lines given in *µ*M as mean ± SEM of at least three replicates (seeding 4,000 cells/well, endpoint 72 h, Alamar Blue assay). **d.** Real time cell analysis (RTCA) curves of Huh-7 and hTERT-RPE1 cells treated with the indicated concentrations of **Ir2** or the vehicle (DMSO).

## Results and Discussion

### 1. Synthesis, characterization, and reactivity of Ir2, Ir2N_3_, IrN_3_

The half-sandwich iridium(III) complexes **Ir2**, **Ir2N_3_** and **IrN_3_** were synthesized according to previously described procedures^14^. Their chemical synthesis and characterization by NMR spectroscopy and high-resolution mass spectrometry (HRMS) are detailed in the Supporting information. Consistently with our previous results, dissolution of the complexes in DMSO vehicle rapidly afforded the cationic species resulting from chlorido to DMSO ligand exchange as observed by ^1^H NMR spectroscopy, with half-lives of 5, 10 and 11 min for **Ir2**, **Ir2N_3_** and **IrN_3_**, respectively (**Figure S2a**). The cationic *solvento* adducts **IrN_3_-DMSO** and **Ir2-DMSO** were also quantitatively isolated by AgPF_6_ treatment. **Ir2-DMSO** crystallized in P2_1_ (monoclinic) space group whereas complexes **IrN_3_** and **Ir2N_3_** crystallized in P2_1_/n (monoclinic) space group. Their X-ray diffraction crystal structure shows the classical piano-stool geometry around the iridium atom (**Figure S1**). Selected geometric parameters, bond lengths and bond angles can be found in **Tables S1-2**.

Formation of covalent adducts by reaction of **Ir2** with various amino acids and side chain models was probed *in vitro* by ESI-HRMS in positive ionization mode. Expectedly, **Ir2** did not form adducts with the aliphatic amino-acids alanine and proline, however they were systematically observed for amino-acids comprising a sulfur or nitrogen donor atom on their side chain (N-acetyl histidine, **Figure S2b** and N-acetyl cysteine-methyl ester, **Table S3**). In addition, formation of an adduct between **Ir2** and L-methionine, by coordination of its sulfur atom, was evidenced by ^1^H NMR spectroscopy in deuterated methanol (**Figure S2c**). Formation of monovalent adducts is a direct consequence of the lability of the chlorido ligand in **Ir2**, in sharp contrast with bis-cyclometallated complexes that are kinetically inert.

Of great interest, our previous SAR study showed that **Ir2** was able to spontaneously produce H_2_O_2_ in the cell cytoplasm via a catalytic mechanism presumably combined to depletion of intracellular NAD(P)H. Herein, we checked whether its congeners **IrN_3_** and **Ir2N_3_** displayed the same feature *in vitro* (**Figure S3**). Confirming our previous observations, we found that **Ir2** and **Ir2N_3_** were more potent than **IrN_3_** to catalytically produce H_2_O_2_ from NADH and O_2_ in model conditions, highlighting again the importance of the dimethyl substituent on the oxazoline ring for pro-oxidant activity.

### 2. Toxicity and pro-oxidant properties of the complexes

The toxicity of **Ir2**, **Ir2N_3_** and **IrN_3_** was investigated on a selected panel of human cell lines. As in our previous studies, the HeLa (late-stage cervix adenocarcinoma) and near-diploid hTERT- RPE1 (telomerase-immortalized retinal pigment epithelial cells) cell lines were used as models of tumoral and non-tumoral cells, respectively. The panel of cancer cell lines was completed with the hepatocarcinoma Huh-7 and the ovarian carcinoma A2780 and its cisplatin resistant derivative A2780cis, representative of a highly sensitive cell line^17^. The HEK293 cell line (adenovirus-immortalized human embryonic kidney cells) that displays a broad spectrum of expressed proteins was chosen for chemoproteomics. The median inhibitory concentration (IC_50_) of cell proliferation was evaluated for each complex (**Figure 1c**), showing ranges of IC_50_ comprised between 2 and 7 µM for **Ir2** and **Ir2N_3_** and between 6 and 11 µM for the less cytotoxic **IrN_3_**. Interestingly, the iridium-based drugs were potent in both cisplatin-resistant and sensitive A2780 cells. Furthermore, in all cell lines, **Ir2** and **Ir2N_3_** displayed similar IC_50_ values, confirming that **Ir2N_3_** is an appropriate surrogate to probe the intracellular effects of **Ir2**.

The cell response to **Ir2** was also investigated by a label-free viability assay based on impedance readout, using the Real Time Cell Analysis (RTCA) system for dynamic monitoring of live cell adhesion and proliferation (**Figure 1d**). According to the proliferation curves, hTERT-RPE cells displayed a distinct response to treatment, as cells survived and proliferated when exposed to 5 µM **Ir2**. In contrast, proliferation of Huh-7 tumor cells was inhibited in a dose-dependent manner. Furthermore, the proliferation curves of treated cells at toxic concentrations indicate a very rapid effect of **Ir2** on cell adhesion and viability. Consistently, accumulation of **Ir2** in cells was observed after 30 min by assaying the elemental Iridium concentration by ICP-OES (Inductively Coupled Plasma - Optical Emission Spectrometry). In HeLa cells, the intracellular content of **Ir2** was found to be dose-dependent with 405 ± 3 and 855 ± 14 ng of Ir per million cells after exposure to 5 and 10 µM, respectively, indicating an efficient intracellular accumulation of the compound.

Taking a mean HeLa cell volume of 3700 µm^3^, the intracellular concentration of elemental iridium was as high as 570 or 1200 µM, respectively. This 100-fold concentration ratio indicates that HeLa cells display a typical “sponge-like” behavior^37^ towards **Ir2**, i.e. displacement of the in/out equilibrium of the free molecule that might be driven by the formation of intracellular covalent adducts (see below).

By contrast, the intracellular concentration of elemental iridium was only 130 ± 2 ng per million cells after 30 min exposure of hTERT-RPE1 cells to 5 µM **Ir2**, that is 4 times lower than in HeLa cells. This striking discrepancy between the two cell lines can provide an explanation for the observed threshold effect on hTERT-RPE1 cell proliferation (**Figure 1d**). Indeed, correlation between drug intracellular concentration and toxicity is widely documented and was initially suggested by Sadler *et al.* while studying isoelectronic (C^N) and (N^N)-chelated half-sandwich iridium complexes^6^ and more recently by Pizarro *et al.*^13^

**Ir2** cytotoxicity was also assayed in HeLa cells using a classical clonogenicity assay (**Figure S4**). **Ir2** dramatically reduced the ability of cells to form colonies even at a concentration of 1.25 µM. In addition, only 20 % of the cells were still able to survive and form colonies after a transient 24 h exposure to 4 µM **Ir2**, consistent with the results of the above survival assays.

The fluorescent H_2_O_2_-selective ratiometric probe “HyPer” developed by Belousov et al.^38^ was used to confirm the pro-oxidant properties of **Ir2** and **Ir2N_3_** *in cellulo* (**Figure 2a**). The fluorescence properties of HyPer change upon its reversible oxidation by H_2_O_2_ and reduction by glutathione-dependent reductases, thus monitoring intracellular hydrogen peroxide levels expressed as HyPer index^14^. Cells exposed to 100 µM H_2_O_2_ as positive control exhibited a fast recovery as the HyPer index returned to its basal value within one hour. In contrast, upon treatment with **Ir2** and **Ir2N_3_** in the same time lapse, the HyPer index rose progressively, indicating a continuous catalytic production of H_2_O_2_ and/or defective detoxification processes by the cells challenged with the drugs. **IrN_3_** was less active in this assay, consistent with its higher IC_50_. These results are in full agreement with the respective ability of the complexes to oxidize NADH *in vitro* (**Figure S3**).

**Figure 2.**
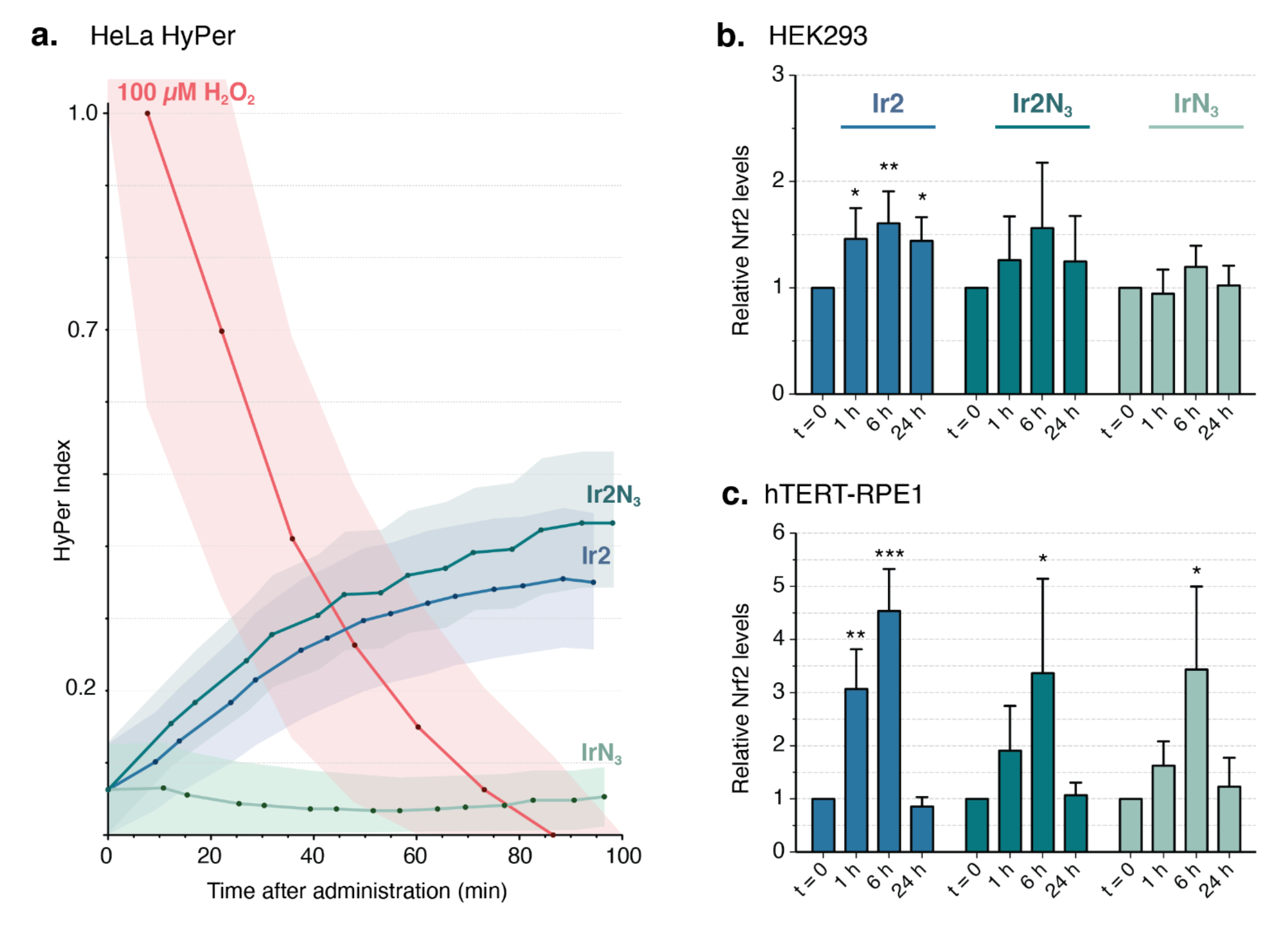
Production of H_2_O_2_ and antioxidant response **a.** Real-time monitoring of the HyPer index in live cells treated with H_2_O_2_, **Ir2**, **Ir2N_3_,** or **IrN_3_**. A stable HeLa cell line that constitutively expresses the H_2_O_2_ probe HyPer was exposed to the three complexes, then reduced and oxidized forms of HyPer were quantified by flow cytometry as the ratio of fluorescence emission at 520 nm after excitation at 405 and 488 nm respectively. The mean ratios of the HyPer-expressing cells (N = 5,000 – 10,000) were used to calculate a HyPer index relative to the negative (DMSO) and positive (H_2_O_2_) controls. Western Blot analysis of the antioxidant response in **b.** HEK293 and **c.** hTERT-RPE1 cells. Semi-quantification of Nrf2 levels following exposure to the three complexes during 1, 6 or 24 h, relative to untreated cells (t = 0) and calculated from at least 3 independent experiments. Statistical significance was calculated using one-way analysis of variance with a Dunnett’s post-test compared to t=0 (*p < 0.05; **p < 0.01; ***p < 0.001).

ROS production in cells is known to elicit the antioxidant response, marked by the accumulation and activation of the transcription factor Nrf2^39^. Western blot analysis (**Figure 2b**) showed that in the HEK293 cell line, only **Ir2** and **Ir2N_3_** were able to induce a mild upregulation of the transcription factor that was sustained throughout 24 h of treatment. In hTERT-RPE1 cells, a higher increase of Nrf2 was observed in response to **Ir2** with respect to the azido-substituted complexes, consistent with the respective IC_50_ of the three complexes. Moreover, hTERT-RPE1 cells were markedly able to cope with electrophilic or oxidative stress induced by the complexes, with Nrf2 returning to basal levels after 24 h. Once generated, ROS can diffuse throughout the cell and indirectly induce DNA double strand breaks^40^. Therefore, we monitored the presence of phosphorylated histone H2AX (gamma-H2AX) foci marking regions of DNA double strand breaks in HeLa and hTERT-RPE1 cells treated with **Ir2**. Conversely to Etoposide, used as positive control, **Ir2** did not significantly increase the number of DNA damage foci within 48 h (**Figure S5**), suggesting that the DNA damage response pathway is not directly implicated in cytotoxicity.

To sum up, both iridium complexes **Ir2** and **Ir2N_3_** reported herein rapidly increase H_2_O_2_ intracellular levels, prompting the activation of the transcription factor Nrf2 as a response to oxidative stress. This pro-oxidant property participates to the antiproliferative activity of these molecules on cancer cells and more generally of metal-based drug candidates.^41^

### 3. Target profiling by click-based chemoproteomics

Prior to biological experiments, the CuAAC reaction between **IrN_3_** and phenylacetylene as model substrate was carried out in a chemical set-up. The 1,4-triazole produced by this reaction was only formed using a DMSO/H_2_O (5:1) mixture as solvent, where DMSO acted as competitor ligand hindering the direct insertion^42, 43^ of the ethynyl function of phenylacetylene at the labile position site. The clicked half-sandwich complex produced *in vitro* was characterized by NMR and HRMS (**Supporting information**). This result gave us confidence that this reaction can be transposed *in cellulo*, where the labile position site is expected to be occupied by amino acids’ lateral chain of the protein targets.

Next, a gel-based method was undertaken as a proof of concept to determine optimal conditions of the click reaction. For this, protein extracts from cells treated with the clickable iridium compounds were labeled with ethynyl-biotin (**Figure S6**). Under these conditions, a whole range of protein adducts were revealed by streptavidin peroxidase in addition to the endogenous biotin-associated proteins. Performing the click reaction on cell extracts rather than intact cells and adding a step of diafiltration after cell lysis enhanced the subsequent detection of protein targets, as previously shown^44^.

A click-on-resin strategy was employed to pull down proteins covalently bound to **Ir2N_3_** using ethynyl-substituted agarose beads. HEK293 cells were cultivated for 20 min in the presence of **Ir2N_3_** at 0.2 and 5.0 µM, to assess for dose-dependent effects, and compared to untreated cells. After on-bead trypsin digestion, mass spectrometry analysis of the samples, each in quintuplicate, led to the absolute quantification of 3686 proteins at the concentration of 5 µM **Ir2N_3_**, 2140 proteins at 200 nM **Ir2N_3_**, and 1594 proteins in the control sample, expressed in molar percentage (Absolute quantification, **Figure S7a**). In the differential analysis, proteins enriched in the iridium-treated samples were selected according to the criteria of at least 2 distinct peptides in 3 replicates with a fold change ≥ 1.5 compared to the control and a p-value ≤ 0.05. 211 proteins were selected from the 200 nM treated sample and termed high affinity targets (List A, **Table S4**) while proteins selected from the 5 µM treated sample constituted List B. As expected, 96% of the proteins from List A were also present in List B. Proteins belonging to List B but not List A were compared according to their fold change between 0.2 and 5.0 µM conditions. In this comparison, 388 proteins having a fold change comprised between 1 and 16 were qualified as intermediate affinity targets, and 786 proteins only captured at 5 µM were defined as low affinity targets (**Figure S7b**).

High affinity targets ranked according to their fold change between 0.2 and 5.0 µM conditions and p-value are shown in the volcano plot, **Figure 3a**. GO term analysis was used to highlight the predominant molecular functions amongst these 211 targets. Cytoskeletal protein binding (GO:0008092), RNA binding (GO:0003723) and Unfolded protein binding (GO:0051082) were the most significant categories (p-value ≤ 6.10^-6^). The 50 most abundant high affinity targets were selected according to their absolute quantification in the 5.0 µM condition for further GO analyses. Proteins with GO terms “Unfolded protein binding” and “Cytoskeleton” in the respective categories molecular function and cellular component, displayed the highest enrichment compared to their background frequency in the human proteome (**Figure 3b**). According to the Cellular Component GO analysis, these 50 targets were found in the cytoplasm and in a variety of organelles (nucleus, mitochondria, ER).

**Figure 3.**
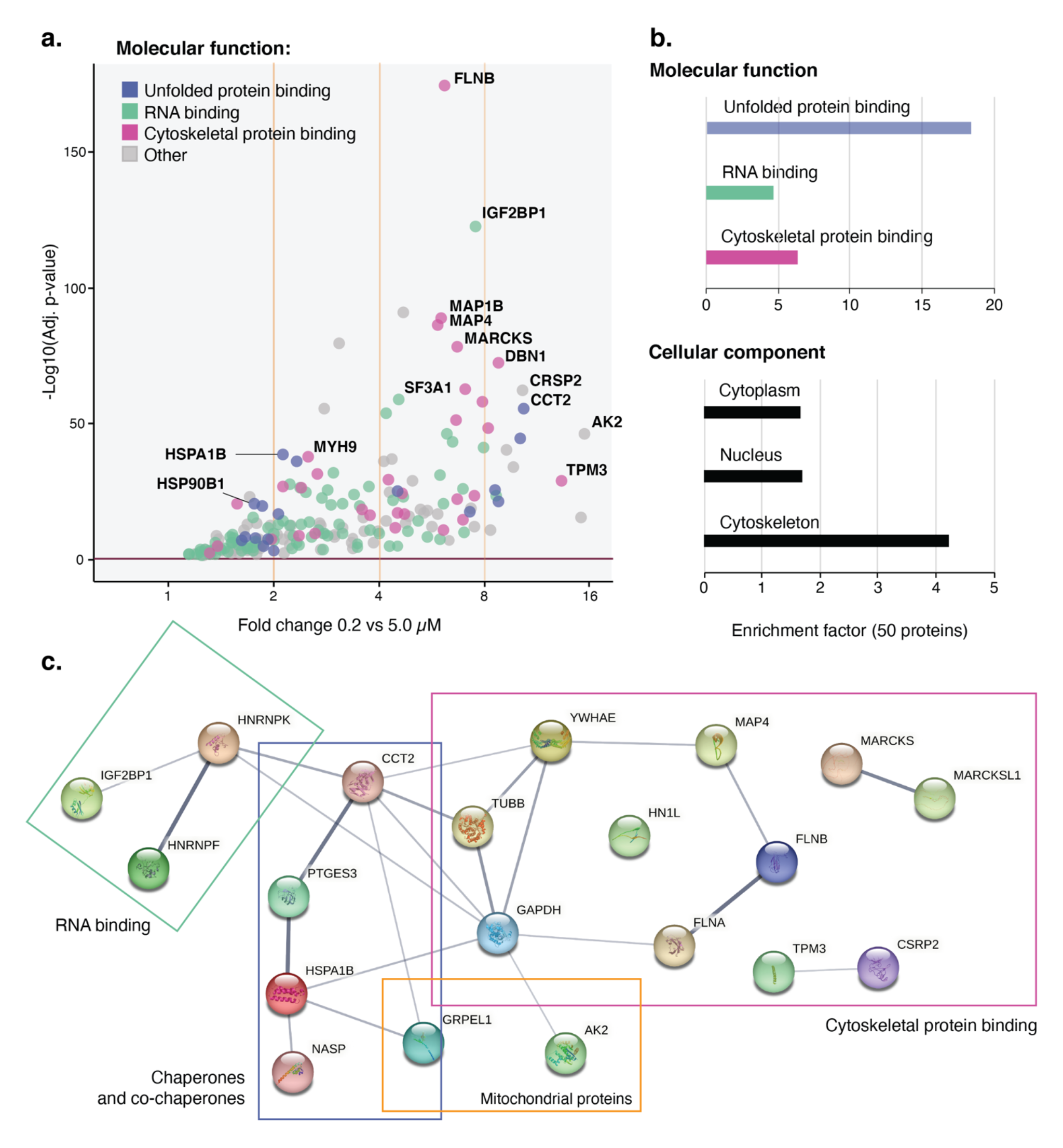
Identification of the intracellular targets of **Ir2N_3_** using “click” chemoproteomics **a.** Volcano plot showing statistical significance (p-value) versus magnitude of change (fold change) for the 211 “high affinity” targets captured at two concentrations of **Ir2N_3_** in HEK293 cells. Color-annotation of the most represented molecular functions. **b.** Ratio between the set and background frequency of representative GO annotations in each category, performed over the 50 most abundant proteins amongst the high affinity targets, according to their absolute quantification in the 5 *µ*M condition. **c.** STRING map of the interaction network between the 20 most abundant proteins amongst the high affinity targets, according to their absolute quantification in the 5 *µ*M condition.

The 20 most abundant high affinity targets are shown in the STRING interaction map (**Figure 3c**). Interestingly, the same GO terms are represented within this reduced list, containing 5 chaperones and co-chaperones, 3 RNA-binding proteins and 11 proteins related to cytoskeleton, including mostly actin binding proteins. In addition, two mitochondrial proteins are present, AK2 and GRPEL1.

### 4. Intracellular localization and target validation

To obtain an unbiased view of the sites of accumulation of **Ir2** in hTERT-RPE1 living cells, we used an advanced method of cryo-fixation followed by synchrotron high resolution X-ray fluorescence (XRF) mapping of elements at the ID16a beamline of ESRF (**Figure 4a**). This method preserves all intracellular elements including the highly diffusible ions like K (uniform subcellular distribution^45^) and highlights several compartments such as the nucleus with P and Zn and the Golgi apparatus with Mn (stored at physiological level in the Golgi apparatus in human cells^46, 47^). The mapped XRF signal of iridium was readily detected in the cytoplasm and the nucleus and appeared enriched on structures identified as mitochondria, actin bundles and nuclear envelope when compared to cryo-optical fluorescence microscopy using vital markers^48^. This distribution fits well with the Cellular Component GO analysis from the chemoproteomics. Quantitation of iridium in the whole cell shown in **Figure 4a** led to an estimate of 70 fg consistent with the ICP-OES bulk analysis performed in parallel, giving an iridium content of 130 ± 2 fg per cell.

**Figure 4.**
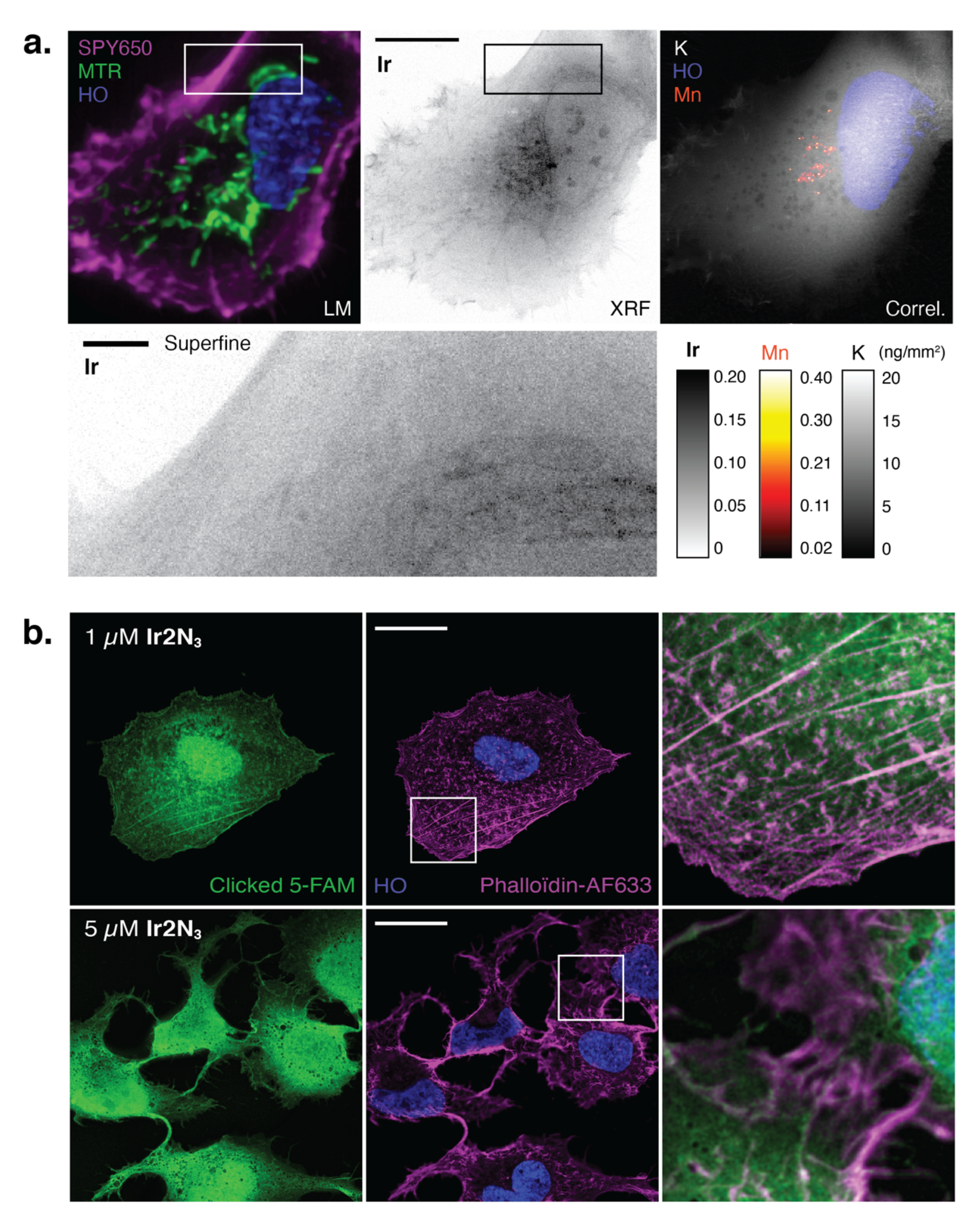
Intracellular detection of **IrN_3_**, **Ir2N_3_** and **Ir2** highlighting nucleus, cytoskeleton and mitochondria localization **a.** Correlative analysis of a frozen-hydrated hTERT-RPE1 cell treated with 5 *µ*M **Ir2** for 15 min and imaged at 88 K. Light cryomicroscopy (LM) of the actin cytoskeleton (SPY650-Fast Act, magenta), mitochondria (Mitotracker Green, MTR) and the nucleus (Hoechst 33342, blue, HO). Cryo-X-Ray Fluorescence imaging (XRF) of elemental Ir, Mn and K expressed in ng/mm_2_ as color-scales of Amber, Fire and Gray respectively, acquired at 50 nm/px/50 ms or “superfine” 30 nm/px/100 ms resolution. Scale bars: 10 and 1 *µ*m. **b.** Super-resolution confocal images of Huh-7 cells exposed to **Ir2N_3_** for 1 h (top 1 *µ*M, bottom 5 *µ*M), fixed and clicked *in situ* with ethynyl-FAM (green). Staining of actin cytoskeleton (phalloidin, magenta) and DNA (Hoechst 33342, blue). Zoom on a selected region of 200 *µ*m^2^ showing colocalization of **Ir2N_3_** and microfilaments (white) on the merged images. Scale bar: 20 *µ*m.

Taking advantage of the bioorthogonal azido function carried by **Ir2N_3_**, its intracellular distribution was also analyzed using in-cell labeling by click chemistry^49^ with a fluorophore introduced in fixed samples, once the complex has reached its intracellular biological targets (*in situ*). Using this approach, only the strongly bound and chemically accessible iridium complex pool and not the free and therefore diffusible one should be observed by fluorescence microscopy. Both complexes **IrN_3_** and **Ir2N_3_** clicked to the ethynyl-substituted derivative of fluorescein (5-FAM) displayed a constant staining pattern at various concentrations and were detectable from 15 min post treatment. The acquired images are consistent with the XRF data described above, supporting that **Ir2N_3_** and **IrN_3_** (**Figure 4b** and **Figure S8**) are detectable using in-cell click chemistry in both the nucleus and cytoplasm. Notably, clicked iridium complexes co-localized with the actin cytoskeleton and perturbed these fibers in a dose-dependent manner. Stress fibers (large contractile filaments of actin and myosin) identified with fluorescent phalloidin, were stained at low concentration of **Ir2N_3_** with 5-FAM as shown in **Figure 4b**. At concentrations of complexes around the IC_50_, they were not detectable and the cortical actin network appeared diffuse and disorganized, indicating that actin cytoskeleton with its associated proteins is a key target of the tested iridium compounds, in full agreement with the presence of relevant actin-binding proteins in the chemoproteomic analysis.

**Ir2N_3_** was also readily detected in the nucleus, again in agreement with the XRF results for **Ir2** and consistent with our detection of nuclear proteins in the iridium interactome. A possible interaction of **Ir2N3** with other nuclear components, in particular DNA cannot be formally excluded. Indeed, other half-sandwich Ir complexes were shown to form adducts with the model nucleobase 9- ethylguanine *in vitro*^50, 51^. However, as mentioned above, the absence of DNA damage in **Ir2** treated cells further argues against a possible interaction of **Ir2** with nuclear DNA *in vivo*.

In order to probe the interaction of **Ir2N_3_** with several protein targets representative of predominant molecular functions identified in our GO term analysis, we introduced a proximity ligation assay (PLA) in which **Ir2N_3_** was clicked to biotin and the protein target of interest was labelled with a specific primary antibody. Then PLA was performed using selective oligonucleotide-labelled antibodies to biotin and the primary antibody (**Figure S9**).

Strong and dose-dependent PLA signals were obtained with the chaperone HSP90 and the actin-binding protein filamin B, belonging to the group of **Ir2N_3_** “high affinity” targets. Two proteins with similar functions, the chaperone GRP78 (intermediate affinity) and the actin-binding protein Arp2/3 (absent from the iridium interactome), gave little to no PLA signal after **Ir2N_3_** treatment (**Figure 5a**). We thus conclude that HSP90 and Filamin B are preferred **Ir2N_3_** protein targets in hTERT-RPE1 cells as well.

**Figure 5.**
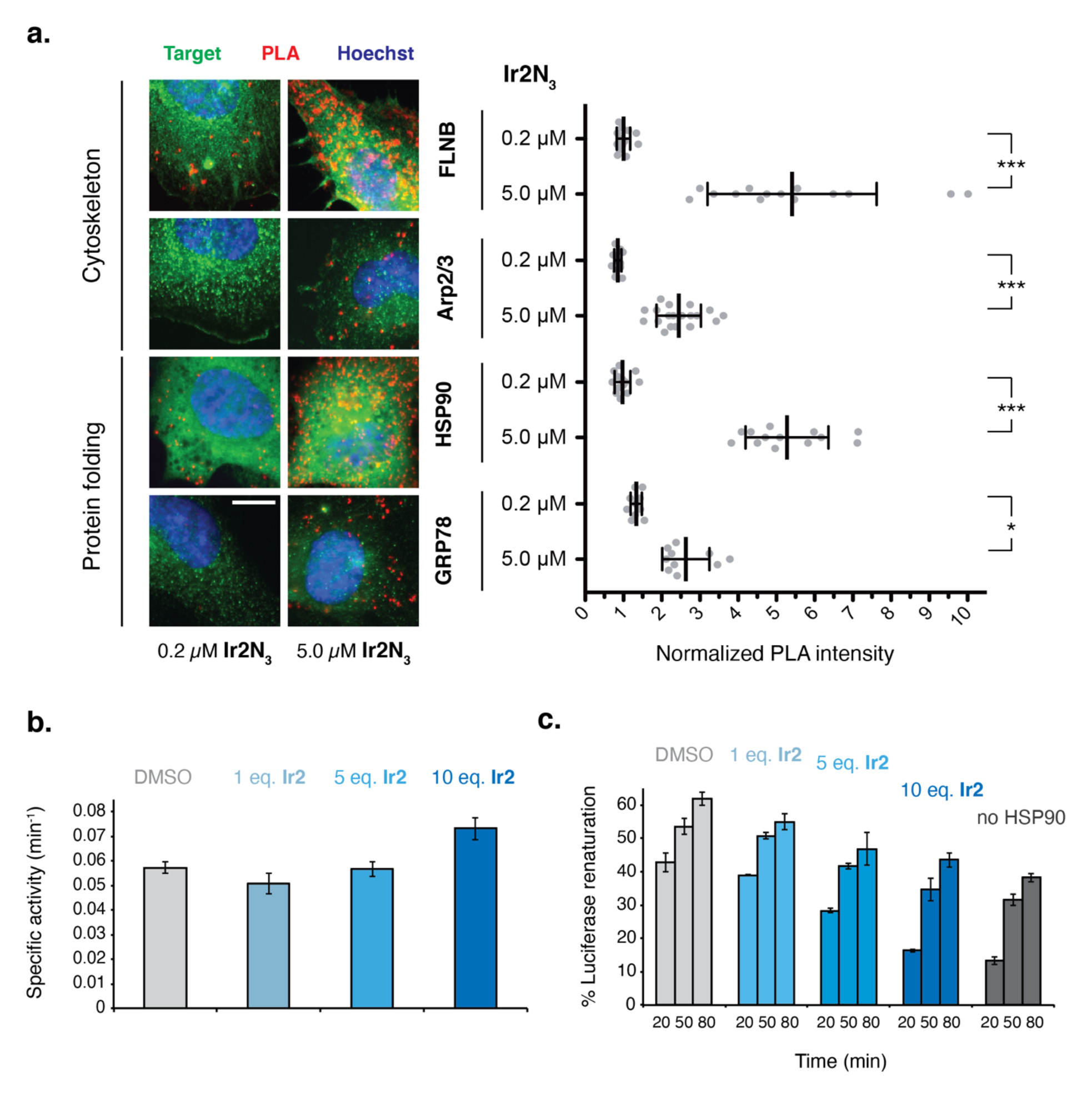
Validation of selected targets by Proximity Ligation Assay (PLA) and functional *in vitro* assays **a.** Quantification of the PLA signal in hTERT-RPE1 cells treated with **Ir2N_3_** for 15 min at the two indicated concentrations. Representative images of the PLA signal (red) and co-immunofluorescence of the protein target (green), DNA stained with Hoechst 33342 (blue). Scale bar: 5 *µ*m. PLA signal intensity per cell (N = 10 – 20) was normalized using a control condition with the target-antibody alone. Statistical significance was calculated using one-way analysis of variance with a Bonferroni post-test (*p < 0.05; ***p < 0.001). **b.** Specific ATPase activity of HSP90, either treated with DMSO vehicle (light grey) or with 1 (light blue), 5 (medium blue) or 10 (dark blue) molar equivalents of **Ir2**, given as mean of three replicates ± SD. **c.** HSP90 foldase activity reported as the percentage of luciferase renaturation after 20, 50 or 80 minutes of incubation in a reconstituted chaperone system with HSP90, either treated with DMSO vehicle (light grey) or with 1 (light blue), 5 (medium blue) or 10 (dark blue) molar equivalents of **Ir2**, or in the absence of HSP90 (dark grey). Results are expressed as mean of three replicates ± SD.

The two chaperones HSP90 and HSP70 (encoded by HSP90B1 and HSPA1B respectively) identified as high affinity targets were used as a paradigm to investigate the effects of iridium adducts on their biochemical activities. The dose-dependent effect of **Ir2** was probed by two functional *in vitro* assays monitoring their ATPase and ATP-dependent foldase activities. The latter was quantified on a model luciferase substrate using a reconstituted chaperone system composed of HSP90, HSP70 and their respective cochaperones STIP1 and DNAJB1. Although we observed a moderate increase of HSP90 ATPase activity after **Ir2** treatment (**Figure 5b**), we found that **Ir2** strongly inhibited HSP90 capacity to refold luciferase (**Figure 5c**).

We used a similar approach to probe the activities of HSP70 upon **Ir2** treatment and first observed a moderate inhibition of its ATPase activity by **Ir2**. However, when we added to the assay DNAJB1, an HSP40 cochaperone needed for HSP70 chaperone function and accelerating its ATPase activity, we observed a stronger response to the cochaperone stimulation of HSP70 when treated with **Ir2** compared to DMSO (**Figure S10a**). We also looked at the refolding capacity of HSP70 on denatured luciferase and observed no difference in the foldase activity upon treatment with **Ir2** or DMSO vehicle (**Figure S10b**).

The contrasting effects of **Ir2** on the two HSPs revealed distinct modes of action and indicate that **Ir2** does not necessarily inhibit target protein function. We conclude that **Ir2** most probably alters HSP90 ability to recognize its client protein substrates without interfering with its ATPase activity and hence its overall structure.

### 5. Ir2N_3_ effects on the cytoskeleton

We previously observed that cellular motility is rapidly abolished upon treatment with half-sandwich iridium complexes including **Ir2**^14^. Dose-dependent effects of both **IrN_3_** and **Ir2N_3_** observed on cell motility (**Video S1**) and the actin cytoskeleton (**Figure 4b**), together with our finding of numerous microfilament-associated proteins amongst **Ir2N_3_** high affinity targets, prompted us to further investigate the dynamics of actin filaments. Near-diploid hTERT-RPE1 cells were chosen for this analysis as they exhibit a non-transformed phenotype with an unperturbed actin meshwork^52^.

Time-lapse videomicroscopy of the actin cytoskeleton was performed using a fluorescent probe (SPY650-FastAct). Complete inhibition of the exploratory phenotype was observed in hTERT-RPE1 cells upon **Ir2** treatment, in contrast to control cells displaying highly dynamic lamellipodia and filopodia (**Video S2**). The assembly of new actin microfilaments was inhibited immediately after addition of 5 µM **Ir2**, followed by the complete disappearance of large structures (stress fibers) after *ca.* 60 min. Condensation of the fluorescent signal in patches suggested aggregation of shorter microfilaments that accumulated close to the plasma membrane.

Alterations of the cytoskeleton at the nanometer scale were further examined by correlative light and scanning electron microscopy (CLSEM) on cells seeded on fibronectin-coated micropatterns to constrain cellular architecture. This method allows a highly reproducible analysis of cell morphology and facilitates correlative microscopy^53, 54^.

To expose cytoskeletal structures and more generally reveal the topography of non-membranous internal structures before CLSEM acquisitions, hTERT-RPE1 cells exposed or not to 10 µM **Ir2N_3_** for 15 min were extracted with a detergent prior to fixation by dissolving the plasma membrane and removing soluble particles. During this step, a fluorescent phalloidin was employed both to stabilize the actin microfilaments and stain the network, then **Ir2N_3_** was labeled with ethynyl-FAM. Under these conditions, the intracellular distribution of **Ir2N_3_** was unaffected, showing a reproducible colocalization with microfilaments. According to the SEM images, the cytoskeleton of treated cells appears less dense and less organized compared to control cells (**Figure 6a**). Disorganization of cytoskeleton in treated cells was quantified using a directionality analysis (**Figure 6b**) of cells constrained on disc micropatterns, demonstrating that **Ir2N_3_** disrupts cytoskeletal organization after only 15 min of incubation.

**Figure 6.**
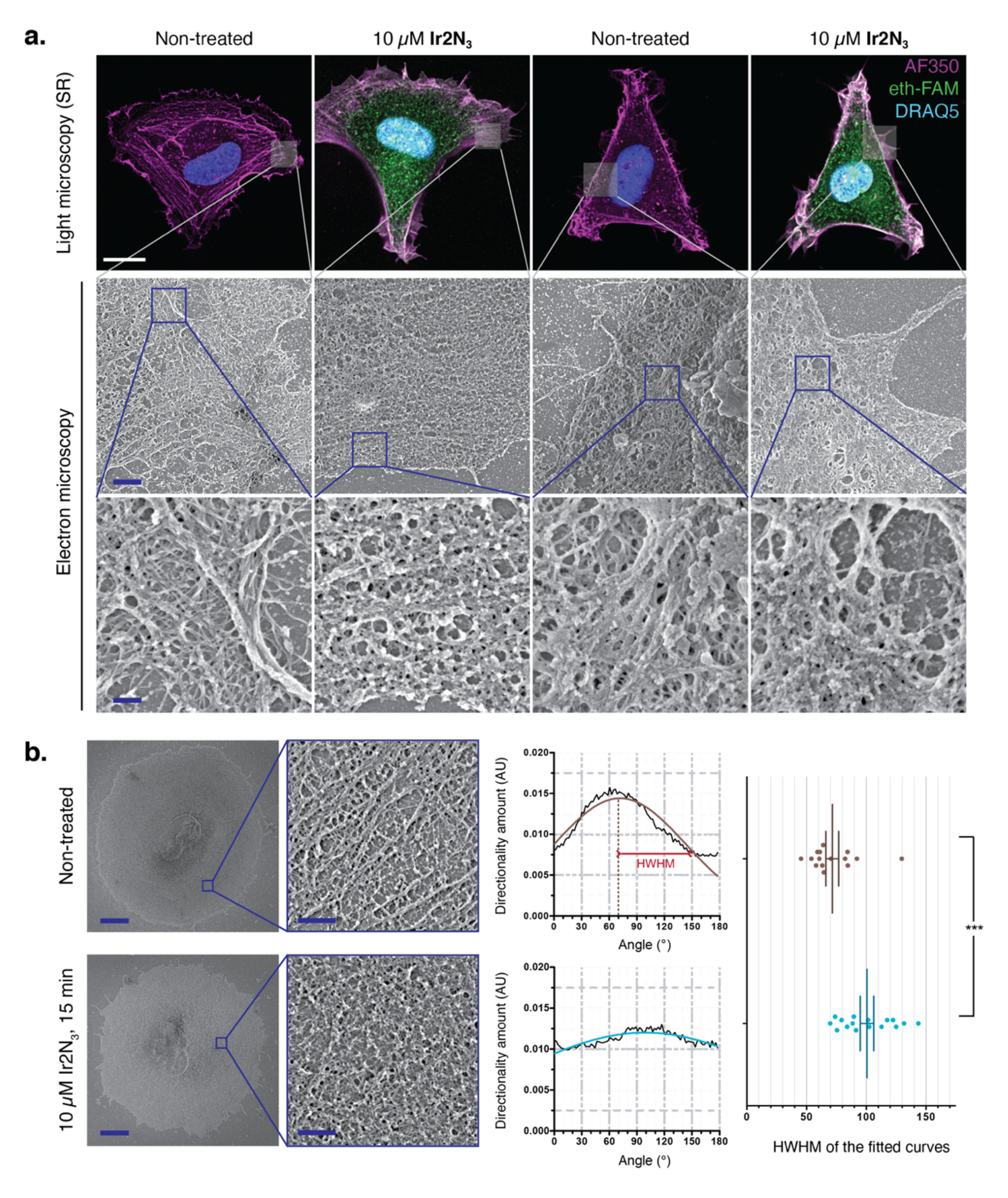
Quantification of cytoskeletal disorganization upon **Ir2N_3_** treatment **a.** Multiscale analysis by Correlative Light and Scanning Electron Microscopy. Cells were seeded on crossbow or arrowhead-shaped fibronectin-coated micropatterns, treated for 15 min with **Ir2N_3_** or vehicle alone, then extracted with detergent prior to fixation. Labelling of **Ir2N_3_** (ethynyl-FAM, green), microfilaments (AlexaFluor-350 phalloidin, magenta) and DNA (DRAQ5, blue), detected by super-resolution confocal microscopy. Correlative scanning electron microscopy. Scale bars: 10 *µ*m, 1 *µ*m, 200 nm respectively. **b.** Cytoskeletal organization of extracted cells seeded on disc-shaped micropatterns of 42 *µ*m diameter. Scale bars: 5 *µ*m, 500 nm. Directionality analysis was performed over 15 – 16 cropped squares in the cortical region of 10 cells per condition. For each image, directionality of cytoskeletal filamentous structures was analyzed using local gradient orientation for each considered angle. The half-width at half-maximum (HWHM) of each fitted gaussian are compared for the treated and control conditions. Statistical significance was calculated using a two-tailed t-test (p = 0.0009, ***).

Alterations of the actin cytoskeleton can result from perturbation of the actin monomer *per se* and its intrinsic ability to polymerize/depolymerize but also of the dynamic interaction of actin monomers and microfilaments with actin-binding proteins. Our chemoproteomic results point to the latter hypothesis as only actin regulators were identified in the iridium interactome, but not actin itself. To rule out any possible direct interference of **Ir2N_3_** with actin polymerization/depolymerization properties, *in vitro* assays were performed. Actin monomer polymerization into microfilaments was not prevented in the presence of 1 or 10 equivalents of **Ir2N_3_**. Likewise, the polymers remained stable in the presence of **Ir2N_3_** (**Figure S11**). We thus conclude that actin cytoskeleton alterations in the presence of **Ir2**/**Ir2N_3_** exclusively arise from their interaction with actin-binding proteins and cross-linkers responsible for the integrity of the cytoskeletal meshwork.

## Conclusion

Our results demonstrate that the cytotoxic effect of the phenyloxazoline-chelated half-sandwich iridium(III) **Ir2** is linked to its ability to rapidly accumulate in cells and to induce both stable accumulation of hydrogen peroxide leading to oxidative stress and formation of covalent adducts with a subset of proteins. This dual chemical reactivity of **Ir2** is shared by its clickable congener **Ir2N_3_**.

Herein, we report the first comprehensive list of protein targets of Ir2 that gives clues to understand the mechanisms underlying its cytotoxicity from the molecular through to cellular levels. Besides, by combining proteomics with imaging, cell biology and biochemistry approaches, we established explicit links between the accumulation of the complex, and its subcellular distribution, protein targets and cellular response.

Proteins belonging to RNA binding, protein folding and cytoskeleton GO terms were the most represented amongst the 211 high affinity and 20 most enriched targets of **Ir2N_3_**. This result is in good agreement with the sites of intracellular **Ir2** accumulation detected by XRF imaging, in particular the nucleus, microfilaments, and mitochondria. We further demonstrated the relevance of two selected targets, the chaperone HSP90 and the actin-binding protein Filamin *in vivo* by a click chemistry-coupled proximity ligation assay.

The ability of **Ir2** to inhibit protein function was demonstrated on HSP90 folding activity *in vitro*, which may be deleterious for cell survival as HSP90 takes part in the maturation of essential signaling proteins. We also propose that **Ir2** interferes with the activity of actin-binding proteins identified in our screen as high affinity targets, including Filamin A/B (exhibiting the lowest p-value) a protein essential to actin filaments crosslinking and cytoskeletal remodeling during cell adhesion and migration. The phenotype of **Ir2** treated cells verifies this hypothesis, as cells exhibit a rapid and dose-dependent disorganization of the cytoskeleton, confirmed by high-resolution imaging approaches, and quantified at the nanometer scale by scanning electron microscopy. This cytoskeleton disorganization is coupled to loss of microfilaments and hence impaired cell motility and adhesion, a phenotype sufficient to trigger anoikis-induced apoptosis^55^.

One of the main hallmarks of cancer is cytoskeletal rearrangement initiated during epithelial-mesenchymal transition that drives metastasis onset. Another feature of some cancer cell types is their higher sensitivity to ROS accumulation^56^ or their higher dependency upon molecular chaperones^57^. In this context, **Ir2** provides a unique combination by targeting cytoskeletal and chaperones proteins in direct relation with cancer progression and by triggering ROS production to selectively elicit cancer cell death. Future approaches will be needed to challenge these various aspects on animal models of tumorigenesis and metastasis.

## Methods

### Antibodies

Rabbit anti-Nrf2 (Abcam #137550); Mouse anti-β-Actin (Cell Signaling #4967); Mouse anti-phospho-Histone H2A.X (Ser139) (Upstate #05-636 clone JBW301); Mouse anti-biotin (Sigma B7653); Rabbit anti-FLNB [N1] (GeneTex #GTX101206); Rabbit anti-P34-Arc/ARPC2 (Merck 07-227-I, lot 3247280); Rabbit anti-HSP90 (Proteintech 13171-1-AP); Rabbit anti-GRP78/BIP (Proteintech 11587-1-AP); Secondary antibodies: Streptavidin-HRP conjugate (R&D systems, lot P237707); HRP-conjugated anti-mouse (Cell Signaling 7076) and anti-rabbit IgG (Cell Signaling #7074); Alexa Fluor® 488 Goat-anti Mouse (Invitrogen, A11001); Alexa Fluor® 488 Goat-anti Rabbit (Invitrogen, A11008).

### Cell Culture

HeLa (human cervix carcinoma), HEK293 (human embryonic kidney) and A2780 cells were obtained from the American Type Culture Collection (ATCC) and cultured in DMEM High Glucose. hTERT-RPE1 cell line was obtained from the ATCC and cultured in DMEM/F12 supplemented with 15 mM HEPES. Huh-7 human hepatocellular carcinoma cell line was obtained from the ATCC, cultured in Minimum Essential Medium (MEM) supplemented with 1 % sodium pyruvate and 1 % non-essential amino acids (Gibco, Invitrogen®). All media were supplemented with GlutaMAX® (Gibco, Invitrogen®), antibiotics (penicillin, streptomycin) and 10 % fetal bovine serum, and were later referred to as complete medium. All short treatments with **Ir2**, **IrN_3_**, **Ir2N_3_** were performed in serum-free media and treatments longer than 3 h were performed in medium containing 5% FBS. Stock solutions of **Ir2** (10 mM), **Ir2N_3_** and **IrN_3_** (20 mM) were prepared in DMSO and incubated 24 h at room temperature before storage, to allow chlorido to DMSO exchange (Figure S2).

### AlamarBlue® assay

4,000 cells were seeded in 100 µL of complete growth media per well in 96-well clear-bottom plates. After allowing cells to grow/adhere overnight, cells were treated with 100 µl per well of 2X drug solutions or vehicle in serum-free growth medium (8 doses ranging from either 40-1.25 µM or 20- 0.625 µM in quadruplicate) and incubated for 72 h. Cell viability was assessed with AlamarBlue® reagent (Life Technologies) according to the manufacturer’s instructions. Normalized fluorescence intensities were plotted with Prism (GraphPad) and IC_50_ values were calculated from 4-parameter nonlinear regression curves. The reported IC_50_ values represent the average of at least three independent determinations (±SEM).

### RTCA

xCELLigence E-plates were calibrated for a baseline definition and cells were seeded at 2,000 cells per well in 100 µL complete medium. After about 24 h, drugs were added at the indicated concentration in triplicate and the cell index was measured over a period of 48-96 h (Real Time Cell Analyzer, Agilent).

### Home-made “click-cocktail”

A stock solution (25X) containing 100 mM CuSO_4_ and 500 mM THPTA was prepared in water and stored at 4°C. 200 mg/mL sodium ascorbate (10X) was prepared extemporaneously in buffer. The click-cocktail was prepared by diluting the alkynyl substrate at a concentration of 4.5 to 10 µM in the buffer, then adding the catalyst premixed solution and finally sodium ascorbate. The cocktail solution turns from blue to yellow, indicating the formation of the Cu(I) species.

### HyPer-based assay

We used the HeLa-HyPer stable cell line expressing the 2^nd^ generation HyPer probe as previously described^14^. Briefly, cells were treated with the drugs (20 µM to a suspension at 500,000 cells/mL in PBS containing 2 % FBS at 37°C) and directly analyzed by flow cytometry (Gallios, Beckman-Coulter, 405/488 nm laser) over 2 h. The mean ratio (M) of Ex488/Em525 and Ex405/Em525 signals was determined. Intracellular hydrogen peroxide level was assessed using a HyPer-index defined as H = (M_sample_- M_DMSO_)/(M_H2O2_-M_DMSO_) where M_DMSO_ and M_H2O2_ correspond to the mean ratio values obtained for the negative (DMSO) and positive (100 µM H_2_O_2_) controls respectively.

### Identification of the protein targets of Ir2N_3_ by mass spectrometry

About 30 million HEK293 cells were used per condition. Cells were cultured in 75-cm^2^ flasks to reach 80 % confluency, washed with 10 mL of warm PBS prior to a 20 min treatment with **Ir2N_3_** at 0.2 or 5 µM (in quintuplicate) in serum-free medium. Untreated cells were used as negative control. Cells were harvested in 10 mL of cold PBS and pelleted by centrifugation. Lysis buffer (200 µL; 200 mM Tris-HCl, pH 8.0, 8 M Urea, 4% CHAPS, 1 M NaCl, supplemented with protease and phosphatase inhibitors, Thermo Scientific) was added to the pellets and the extracts were sonicated 10 x 3s with a probe sonicator. Samples were then diafiltered at 4°C using 3 kDa cutoff VivaSpin®500 centrifuge filters (Sartorius) for 3 cycles of 20 min at 14,000 rpm, with addition of 125 µL of lysis buffer before each cycle. The click reaction was performed according to the supplier’s protocol: 50 µL of ethynyl-beads suspension (Jena biosciences) was mixed with 200 µL of extract and 250 µL of click cocktail. The CuAAC was allowed to perform overnight at 4°C on a wheel. Resin-bound proteins were then reduced with DTT and alkylated with iodoacetamide according to the supplier’s instructions. The resin was thoroughly washed 5 times with agarose-SDS buffer, 5 times with 8 M Urea in 100 mM Tris-HCl pH 8.5 and 5 times with 20 % MeCN in water. Beads were resuspended in 500 µL of 100 mM Tris-HCl pH 8.0 containing 2 mM CaCl_2_ and digested at 37 °C for 1 h with 0.2 μg of trypsin/LysC (#V5073 Promega). The samples were then loaded onto homemade SepPak C18 Tips packed by stacking three AttractSPE disk (#SPE-Disks-Bio-C18–100.47.20. Affinisep) into a 200 μL micropipette tip for desalting. The peptides were eluted with MeCN/H_2_O 40:60 with 0.1 % formic acid and vacuum concentrated to dryness. The samples were resuspended in 10 μL of 0.3% TFA before LC-MS/MS analysis.

### LC-MS/MS analysis and data processing

Samples (5 µL) were chromatographically separated using an RSLCnano system (Ultimate 3000, Thermo Scientific) coupled online to an Orbitrap Eclipse mass spectrometer (Thermo Scientific). Peptides were first loaded onto a C18 trapped column (i.d. 75 μm x 2 cm, nanoViper Acclaim PepMap^TM^ 100, Thermo Scientific) with eluent A (MeCN/H_2_O 2:98 with 0.1 % formic acid) at a flow rate of 3.0 µL/min over 4 min and then switched for separation to a C18 column (i.d. 75 μm x 50 cm, nanoViper C18, 2 μm, 100 Å, Acclaim PepMap^TM^ RSLC, Thermo Scientific) regulated to a temperature of 50°C with a linear gradient from 2 % to 25 % eluent B (MeCN with 0.1 % formic acid) at a flow rate of 300 nL/min over 91 min. MS1 data were collected in the Orbitrap (120,000 resolution; maximum injection time 60 ms; AGC 4e5). Charges states between 2 and 5 were required for MS2 analysis, and a 45 s dynamic exclusion window was used. MS2 scans were performed in the ion trap in rapid mode with HCD fragmentation (isolation window 1.2 Da; NCE 30 %; maximum injection time 60 ms; AGC 1e4). For identification, the data were searched against the *Homo sapiens* (UP000005640_9606) UniProt database using Sequest HT through proteome discoverer (version 2.4). Enzyme specificity was set to trypsin and a maximum of two miss cleavages sites were allowed. Oxidized methionine, Met-loss, Met-loss-Acetyl and N-terminal acetylation were set as variable modifications. Carbamidomethylation of cysteines was set as fixed modification. Maximum allowed mass deviation was set to 10 ppm for monoisotopic precursor ions and 0.6 Da for MS/MS peaks. The resulting files were further processed using myProMS v3.9.3 (https://github.com/bioinfo-pf-curie/myproms)^58^. FDR calculation used Percolator^59^ and was set to 1% at the peptide level for the whole study. The label free quantification was performed by peptide Extracted Ion Chromatograms (XICs), reextracted across all conditions and computed with MassChroQ version 2.2.21^60^. For protein quantification, XICs from proteotypic peptides shared between compared conditions (TopN matching) and missed cleavages were allowed. Median and scale normalization was applied on the total signal to correct the XICs for each biological replicate (N=5 in each condition) for total signal and global variance biases. To estimate the significance of the change in protein abundance, a linear model (adjusted on peptides and biological replicates) was performed, and p-values were adjusted using the Benjamini–Hochberg FDR procedure. Also, for each treatment condition (NT, 0.2 µM and 5 µM), we computed the expression of proteins as a molar proportion and mass percentage estimated by using top 3^61^ (including proteins with less than 3 peptides) as the Protein Quantification Index and the direct proportionality model^62^.

### CuAAC of ethynyl-PEG_4_-biotin on adherent cells. DuoLink® PLA and immunofluorescence

100,000 hTERT-RPE1 cells were seeded on 12-mm diameter coverslips and cultured overnight in complete medium. Cells were washed with PBS and incubated for 15 min in serum-free medium containing **Ir2N_3_** at 0.2 or 5 µM or left untreated. Cells were washed and fixed for 15 min at room temperature in 4 % PFA in PBS containing 1 µg/mL of Hoechst 33342. After four PBS washes, the click reaction was performed using the cocktail containing 5 µM ethynyl-PEG-biotin (Lumiprobe) for 30 min at room temperature. Cells were washed twice with PBS and incubated for 90 min with 3 % BSA in PBS with 0.3 % Triton X-100. Coverslips were incubated overnight at 4°C with Mouse anti-Biotin antibody (1:1000) and/or Rabbit anti-[FLNB, HSP90, GRP78, P34-Arc/ARPC2] (1:100). The DuoLink® proximity ligation assay was carried out according to the supplier’s instructions and all slides were incubated for 1 h with an Alexa488-anti-Rabbit secondary antibody (or Alexa488-anti-mouse for a subset of controls). Slides were imaged using a Leica DM4 B upright microscope with a 63x objective and signal intensities were quantified using Fiji.

### Effect of Ir2 on the activities of HSP molecular chaperones *in vitro*

<u>Protein labeling by Ir2:</u> Purified recombinant Heat Shock Proteins (see also **Supplementary Information**) were diluted in labeling buffer (25 mM Tris-HCl pH=7.5; 150 mM NaCl; 5% glycerol) to a final concentration of 50 µM. 1, 5 or 10 molar eq. of **Ir2**, or DMSO vehicle, was added (corresponding respectively to final concentrations of 5, 50 or 500 µM **Ir2** and 5 % DMSO). Samples were incubated at room temperature during one hour and labeled proteins were separated from **Ir2** and DMSO vehicle by 5 cycles of diafiltration (12,000 g, 4°C, molecular cutoff 10kDa, Sartorius) adding 5 volumes of labeling buffer before each cycle. **Ir2** monitored by absorption in the diafiltration flow-through was undetectable in 1 and 5 eq. **Ir2** samples and present at 8 µM in the 10 eq. **Ir2**. In order to eliminate remaining free **Ir2**, proteins were further purified by size-exclusion chromatography with PD-10 column (Cytiva) and elution with labeling buffer. Samples were concentrated and the protein concentration measured by BCA assay kit (Sigma). <u>ATPase assay:</u> Steady-state ATPase activity of Heat Shock Proteins was evaluated by a coupled enzymatic assay (PK/LDH) monitoring the oxidation of NADH as described previously^63^ with a modified reaction buffer (50 mM Hepes pH=7.5, 100 mM KCl, 5 mM ATP; 5 mM MgCl_2_) in 96 well plates on a CLARIOstar reader (BMG Labtech). Heat Shock Proteins concentrations were either 10 µM for HSP90 or 5 µM for HSP70 (**Supplementary Information**). The HSP90 ATPase activity was corrected by subtraction of the background ATPase activity after addition of 50 μM radicicol. <u>Luciferase refolding assay:</u> QuantiLum Recombinant Luciferase (Promega) was diluted to 55 µM in denaturation buffer containing 6M Guanidium/HCl and 1 mM DTT during 30 minutes at room temperature. Denatured luciferase was diluted 125 times (0.4 µM) in renaturation buffer (20 mM Tris-HCl pH=7.5; 50 mM KCl; 5 mM MgCl_2_; 1 mM DTT) supplemented with HSP samples treated with **Ir2** or DMSO vehicle. Final concentrations of recombinant chaperones and cochaperones were: 10 µM HSP70, 2 µM DNAJB1 and, for the evaluation of HSP90 activity, 0.5 µM STIP1 and 2 µM HSP90. Refolded luciferase activity was measured by mixing of 5 µL aliquots of labeled samples with 120 µL of luciferase buffer (20 mM Tris-HCl pH=7.5; 200 µM luciferin; 0.5 mM ATP; 10 mM MgCl_2_). The chemiluminescence was measured on a CLARIOStar plate reader (BMG Labtech). The percentage of refolded luciferase was evaluated relatively to the activity of native luciferase.

### CuAAC of ethynyl-5-FAM on fixed cells

5.10^4^ HuH-7 cells were seeded on 14 mm coverslips (#1.5) in a 12-well plate and cultured in complete medium. The next day, cells were washed and treated for 1 h with **Ir2N_3_** at 1 or 5 μM in serum-free media. Cells were fixed with 4 % PFA in PBS and washed three times with PBS. Specimens were blocked and permeabilized using PBS containing 3 % BSA and 0.5 % Triton X-100 during 30 min at room temperature. Coverslips were washed with PBS and incubated with the click cocktail containing 4.5 µM ethynyl-5-FAM (Lumiprobe) during 30 min. Finally, samples were washed 3 times with PBS and stained with 1 μg/mL Hoechst 33342 and Alexa Fluor® 633-Phalloidin in PBS during 30 min. Coverslips were washed with PBS and mounted in Prolong Diamond mounting medium. Acquisitions were performed on an inverted AXIO OBSERVER 7 LSM 980 6 CH AIRYSCAN 2 from Zeiss company, with a 63x oil immersion NA 1.4 objective, the following excitation laser lines: 405 nm, 488 nm, 639 nm and the corresponding bandpass filters: BP 422-477, BP 495-550 and BP 655-720. Images were taken and processed with the Airyscan 2 technology in SR (Super-Resolution) mode.

### Micropatterning

35-mm dishes with a #1.5H glass coverslip bottom imprinted with 50 µm cell location grid (IBIDI®) were used as patterning substrates. *<u>Passivation:</u>* Glass bottoms were first glowed with a plasma cleaner to ensure hydrophilicity, then coated for 30 min at room temperature with 0.1 mg/mL PLL (Poly-L-Lysine). After 3 washes with PBS, the dishes were incubated with a freshly made solution of 100 mg/mL PEG-SVa (PolyEthylene Glycol-Succinimidyl Valerate) in HEPES buffer pH 8-8.5 for 45 min at room temperature. *<u>Photopatterning:</u>* Substrates were thoroughly washed with deionized water and patterns were imprinted using a gel photo-initiator (PLPP, Alvéole) and a photomicropatterning device (PRIMO, Alvéole) equipped with a UV laser, delivering doses of 15 mJ/mm^2^. The shapes of the patterns (crossbows 40×40 µm, discs of 35 or 42 µm diameter, arrowheads 40×30 µm) were imprinted using an inverted Axio Observer 7 from Zeiss company with a 20x objective, their position and repetition on the substrate surface were managed using the associated plug-in (Leonardo, Alvéole).

### Correlative light and scanning electron microscopy (CLSEM)

<u>Sample preparation:</u> Buffers used were PEM (PIPES 100mM, MgCl_2_ 1 mM, EGTA 1 mM) and Extraction Buffer^64^ (PEM, 1% Triton X-100, 1% Phalloidin-Alexa350). Micropatterned IBIDI meshed dishes were coated with 50 µg/mL fibronectin (from human plasma, Invitrogen) in PBS for 30 min at 37°C and washed with PBS. Dishes were seeded with 100,000 hTERT-RPE1 cells in 3 mL of complete medium, left to adhere for 2 h. Cells were treated for 15 min in serum-free medium with either 5 µM, 10 µM **Ir2N_3_** or vehicle. Dishes were washed with 3 mL of warm HBSS (with Ca^2+^ and Mg^2+^) and extracted with 200 µL of extraction buffer during 5 min, then cells were fixed with 2 % glutaraldehyde (GA) in PBS for 20 min at room temperature. Cells were washed 5 times with PBS and samples were incubated for 30 min at room temperature with the click cocktail containing 4.5 µM ethynyl-5-FAM (Lumiprobe) to label **Ir2N_3_**. Cells were washed 3 times with PBS, and GA was quenched with two 4 min baths of 1% NaBH_4_ in PBS followed by 3 PBS washes. Samples were kept at 4°C overnight in PBS. Actin cytoskeleton was re-stained for 30 min with Phalloidin-Alexa350 followed by 3 PBS washes. Cell nuclei were stained using DRAQ5 for 20 min followed by 3 PBS washes. The same day, light microscopy acquisitions were performed on an inverted AXIO OBSERVER 7 LSM 980 6 CH AIRYSCAN 2 from Zeiss company, with a 63x oil immersion NA 1.4 objective, the following excitation laser lines: 405 nm, 488 nm, 639 nm and the corresponding bandpass filters: BP 422-477, BP 495-550 and BP 655-720. Images were taken with the Airyscan 2 technology in SR (Super-Resolution) mode and processed with the Airyscan option in Huygens software.

Afterwards, samples were sequentially post-fixed for 10 min each with 1 % aqueous tannic acid then 1 % aqueous uranyl acetate before extensive washing in distilled water^65^, before gradual dehydration with ethanol and hexamethyldisilazane, then drying under vacuum. Finally, micropatterned slides were removed from the dishes, mounted on stubs, and coated by sputtering of 2-3 nm of Chromium (ACE600, Leica microsystems) prior to observation with a high-resolution Field-Emission SEM operating at 1.5 kV (GeminiSEM500, Zeiss) with a 15 μm objective aperture diameter (19 pA). Secondary electrons were collected with an in-lens detector and high-sensitivity Everhart-Thornley (SE2) detectors, scan speed and line integration were adjusted during observation and images were averaged from both detectors. <u>Directionality analysis:</u> A set of 6.25×6.25 µm^2^ square images were acquired in the cortical region of 7-8 cells per condition at high magnification. The images were filtered with a bandpass FFT filter, and the local gradient orientation was analyzed (Directionality Plugin, Fiji) over 36 directions from 0° (east direction) to 180° (counterclockwise), giving a gaussian distribution around the main direction of filaments. The resulting HWHM (half width at half maximum) of fitted curves was calculated using Prism (GraphPad).

### Correlative Light/XRF analysis

Silicon nitride membranes (Silson Ltd) consisting in square silicon frames of 5 x 5 mm^2^ and 200 µm thickness with a central Si_3_N_4_ membrane of 1.5 x 1.5 mm^2^ and 500 nm thickness were used for cell culture and allow contaminant-free XRF low background signal. During manufacturing, a second, smaller (0.1 x 0.1 mm^2^) membrane is added in one of the corners of the silicon frame to serve as orientation object. Membranes were coated for 15 min with 50 µg/mL fibronectin in PBS at room temperature, thoroughly washed with PBS and seeded with 4’000 – 8’000 hTERT-RPE1 cells in 10 µL of complete medium. Cells were incubated for 3-4 h at 37°C and 5% CO_2_ to adhere and spread on the substrates. They were stained for 90 min with SPY650-FastAct (Spirochrome®), including 30 min of co-staining with 100 nM Mitotracker Green FM (Invitrogen®) and 5 µg/mL Hoechst 33342. Membranes were washed and further incubated in medium containing 5% FBS ± 5 µM **Ir2** for 15 min, then rinsed with PBS and serum-free medium. Samples were quickly rinsed with ammonium acetate solution (150 mM) followed by manual blotting and plunge-freezing into liquid ethane chilled with liquid nitrogen^66^. Prior to be transferred for synchrotron X-ray fluorescence cryo-nanoimaging, plunge-frozen cells were imaged using cryo-fluorescence light microscope (cryo-FLM) Leica cryo-CLEM Thunder system equipped with a ceramic-tipped, 0.9 NA, 50X lens. The brightfield and band pass filter cubes of GFP (λ_em_ = 525 nm), DAPI (λ_em_ = 477 nm), and Y5 (λ_em_ = 660 nm) were used. A complete mosaic of the 1.5 x 1.5 mm Si_3_N_4_ active area containing vitrified cellular region of interest was registered with the collection for each field of view of a Z-stack projection over ∼ 20 µm. Images were acquired at 88 K Large Volume Computational Clearance (LVCC) from Leica LAS X Thunder package was applied for fluorescent image deconvolution and blurring reduction on the cryo-FLM image stacks. The ID16A end station is under high-vacuum and is equipped with a cryostage to allow measurements of frozen-hydrated sample kept at 120K and the combined used of LEICA EM-VCM, EM-VCT systems allows the cryotransfer. Nano-positioning is performed by a piezo-driven short-range hexapod stage regulated with the metrology of twelve capacitive sensors. All scanning uses “on-the-fly” acquisition with the sample translated at constant speed in the horizontal direction^67^. The beam was focused to 26 × 42 nm^2^ (vertical × horizontal) using a pair of Kirkpatrick-Baez mirrors^68^. The fluorescence signal emitted from each sample pixel was recorded by 2 custom multi-element Silicon Drift Detectors (SDD) placed on both sides of the sample and facing each other at 90 degrees from the incident x-ray beam. A multi-element SDD (Hitachi Ltd.) and an ARDESIA-16 spectrometer based on monolithic SDD array^69^ were used. The resulting XRF spectra were fitted pixel by pixel using a dedicated Python script to correct for detector deadtime and perform normalization for variation of the incident X-ray beam. The script exploits the PyMca librairies^70^. The elemental areal mass concentration was calculated using the Fundamental Parameters (FP) approach implemented in PyMca software package and a reference standard material containing element of certified concentration (RF7-200-S2371 from AXO, Dresden, Germany) with uniform mass depositions in the range of ng/mm² (1-3 atomic layers). The resulting elemental areal mass density maps were visualized with ImageJ software.

## Data availability

The data that support the findings of this study are available within the paper and its Supplementary Information files or from the corresponding authors upon request. Crystallographic data from the complexes **Ir2-DMSO**, **IrN_3_** and **Ir2N_3_** reported in this Article have been deposited at the Cambridge Crystallographic Data Centre (under deposition numbers CCDC2205966, CCDC22055967 and CCDC2205968, respectively). The mass spectrometry proteomics raw data have been deposited to the ProteomeXchange Consortium via the PRIDE partner repository with the dataset identifier PXD038711 (Username, Password: 7qPwQnXC).

## Acknowledgements

We thank Dr. Patricia Forgez for the gift of ovarian cell lines, Chelly Accipe and Lucile Diot for technical support. We are indebted to Annie Munier, Romain Morichon (CISA platform), Benoit Caron (ALIPP6 platform) and Cédric Przybylski (MSU^3^ platform). We are grateful for the help of Louise Bonnemay, Matthieu Opitz and Hélène Delobel from Alvéole company and Fabrice Schmitt from Zeiss company to imprint micropatterned substrates. PM thanks L. Qin, J. Henri and F. Georgescauld for their help in expression vector design and initial set-up of functional assays. This work was supported by a PhD fellowship from Sorbonne Université and a post-doctoral fellowship from the PCSI program of ITMO Cancer (20CP175-00; R. Ramos), SiRIC CURAMUS financially supported by the French National Cancer Institute, the French Ministry of Solidarity and Health and Inserm (R20113DD; J.S-T), INCa-DGOS-Inserm_12560 (J.S-T), PCSI program of ITMO Cancer supported by Inserm and Aviesan (grant no. 20CP175- 00; M.S and J.S-T). We acknowledge the European Synchrotron Facility (ESRF) for granting beamtime through experiment LS-3136 at beamline ID16a. PM acknowledges the financial support of the French National Research Agency as part of the “Investissements d’Avenir” Program (LabEx Dynamo ANR-11-LABX-0011-01) and ANR16- CE11-0032-03, the French Plan Cancer and the Fondation ARC (PDF20151203636). MS analysis was performed with financial support from “la Région Ile-de-France” (N°EX061034) and ITMO Cancer of Aviesan and INCa on funds administered by Inserm (N°21CQ016-00).

The funders had no role in study design, data collection and analysis, decision to publish or preparation of the manuscript.

## Author contributions

RR Investigation, formal analysis, visualization, writing of original draft. AK, CB, Conceptualization, writing review and editing. MT FL JF SB PM TC CBe DL FD Data acquisition, formal analysis. AC CC MS Data acquisition. MS*, JST* Funding acquisition, data curation, project administration, manuscript editing, joint supervision.

All authors analyzed data, discussed results. The manuscript was written and edited through contributions from all the authors.

