## Supplementary information for "Identification of the intracellular protein targets of a bio-active clickable half-sandwich iridium complex by chemical proteomics"

### Table of Contents

**I. Experimental procedures ........................................................................................3**

1. **Chemical synthesis and characterization**
2. **Supplementary figures-associated protocols and instrumentation**
3. **Heat Shock Proteins cloning and purification**

**II. Supplementary figures, videos and tables ....................................................................12**

**Figure S1.** Crystal structure representations of the three half-sandwich complexes
**Figure S2. Ir2** forms adducts with DMSO, Histidine and Methionine
**Figure S3. Ir2** and **Ir2N$_3$** catalyze the oxidation of NADH and the production of $H_2O_2$ *in vitro*
**Figure S4.** Clonogenicity assay on HeLa cells
**Figure S5. Ir2** does not induce severe DNA damage in HeLa and hTERT-RPE1 cells
**Figure S6.** On-blot detection of **Ir2N$_3$** protein adducts using CuAAC with an ethynyl-substituted biotin and streptavidin peroxidase
**Figure S7.** Absolute quantification and scheme of the chemoproteomic analysis of **Ir2N$_3$** targets
**Figure S8.** Bio-orthogonal labelling of **IrN$_3$** by ethynyl-FAM and correlative LM-XRF imaging of **Ir2** in hTERT-RPE1 cells (in support to **Figure 4**)
**Figure S9.** Principle of PLA and negative controls (in support to **Figure 5a**)
**Figure S10. Ir2** labeled HSP70 functional assays
**Figure S11.** *In vitro* polymerization of actin monomer in presence of **Ir2**

**Video S1.** Cell motility over 9 h.
**Video S2.** Actin cytoskeleton dynamics over 3 h.

**Table S1.** Crystal data and structure refinement for [**Ir2**-DMSO], **IrN$_3$** and **Ir2N$_3$**
**Table S2.** Selected distances (Å ±98% CI) and angles (° ±98% C) for [**Ir2**-DMSO], **IrN$_3$** and **Ir2N$_3$**
**Table S3.** Observed adducts in MeOH with the lateral chain of model substrates (ESI-HRMS); relative intensity of peaks
**Table S4.** Protein lists and GO analyses (link)

1. Cryo-XRF (relevant elements) and photonic raw images acquired in a treated cell
2. Cryo-XRF (relevant elements) and photonic raw images acquired in untreated cells

3. 1H and 13C NMR (DEPT-135) Spectra of **Ir2-DMSO**
4. 1H and 13C NMR Spectra of **L06**
5. 1H and 13C NMR Spectra of **L06N$_3$**
6. 1H and 13C NMR Spectra of **IrN$_3$**
7. 1H and 13C NMR Spectra of **IrN$_3$-DMSO**
8. 1H and 13C NMR Spectra of **phenyl-©-IrN$_3$**
9. 1H and 13C NMR Spectra of **L08**
10. 1H and 13C NMR Spectra of **L08N$_3$**
11. 1H and 13C NMR (DEPT-135) Spectra of **Ir2N$_3$**

### I. Experimental procedures

#### 1. Chemical synthesis and characterization

**Instrumentation.** Reagents were purchased as reagent-grade and used without further purification. All reactions were monitored by analytical TLC on silica gel 60 F254 plates 0.25 mm, and visualized under UV light (λ = 254 and 365 nm). Silica gel (SDS 60 ACC 35–70 mm) or alumina (90 basic, Macherey-Nagel) was used for column chromatography. NMR spectra were recorded on Bruker Avance III 300 MHz or 400 MHz spectrometers at room temperature. Chemical shifts (δ) are expressed in part per million (ppm), reported as s = singlet, d = doublet, t = triplet, m = multiplet; and referenced to the solvent peak of respectively $CDCl_3$, $CD_2Cl_2$, $(CD_3)_2SO$ ($^{13}C$ NMR: δ = 77.23; 53.84; 39.52 ppm; $^1H$ NMR: δ = 7.26; 5.32; 2.50 ppm). ESI-HRMS analysis were carried out using a LTQ-Orbitrap XL from Thermo Scientific (Thermo Fisher Scientific, Courtaboeuf, France) and operated in positive ionization mode. IR spectra were recorded on a FT-IR spectrometer (Tensor27, Bruker) equipped with an ATR MIRacle (Pike Technologies) accessory.

**Ir2-DMSO: [Dimethylsulfoxide-(η5-pentamethylcyclopentadienyl)(2-phenyl-κC2-(4-dimethyl)oxazoline-κN)Iridium (III)] Hexafluorophosphate**

**Ir2**[1] (200 mg, 372 µmol) was dissolved in 5 mL of dichloromethane and 5 mL of MeOH:$H_2O$ (1:1) were added. Another 10 mL of methanol were added to obtain one homogeneous layer. Silver nitrate (69.5 mg, 409 µmol, 1.1 eq) was poured into the solution and a white precipitate immediately appeared. The solution was left stirring for 1 h. DMSO (26.4 µL, 372 µmol, 1.0 eq) was added and the solution turned from golden to pale yellow. Stirring continued for 5 min and $NH_4PF_6$ (339 mg, 2.08 mmol, 5.6 eq) was added. After 15 min, the crude mixture was filtered through a sintered glass frit (Porosity 3 equipped with paper filter) and the filtrate was evaporated. The obtained suspension was redissolved in dichloromethane and the solution was washed with $H_2O$ to remove the $NH_4PF_6$ excess. The product was isolated from the organic phases, after drying with $MgSO_4$ and evaporating volatiles, as a pale-yellow solid (271 mg, 99%). A single monocrystal was obtained in from a concentrated MeOH solution for X-Ray diffraction analysis.

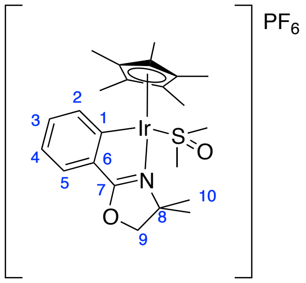

**$^1H$ NMR** (400 MHz, MeOD) δ (ppm): 7.71 (d, $J$ = 7.6 Hz, 1H), 7.59 (dd, $J$ = 7.6, 1.5 Hz, 1H), 7.44 (td, $J$ = 7.5, 1.5 Hz, 1H), 7.29 (td, $J$ = 7.5, 1.0 Hz, 1H), 4.81 (s, 1H), 4.67 (d, $J$ = 8.9 Hz, 1H), 3.22 (s, 3H), 2.53 (s, 3H), 1.90 (s, 15H), 1.59 (s, 3H), 1.51 (s, 3H). **$^{13}C$ {$^1H$} NMR** (101 MHz, MeOD) δ (ppm): 192.4 ($C_1$), 174.7 ($C_6$), 136.9 ($C^{Ar}$), 135.2 ($C^{Ar}$), 134.6 ($C_7$), 129.2 ($C^{Ar}$), 125.8 ($C^{Ar}$), 116.4 ($C_8$) 98.0 ($C_9$), 84.5 ($C^{Cp*}$), 47.0-41.9 ($CH_3^{DMSO}$), 29.6-26.2 ($C_{10}$), 10.0 ($CH_3^{Cp*}$).

**L06: 2-(4-(chloromethyl)phenyl)-4,5-dihydrooxazole**

Obtained following previously described procedure[1] as a yellow solid, 99%. This compound is commercially available but was previously synthesized for this project.

**$^1H$ NMR** (400 MHz, $CDCl_3$) δ (ppm): 7.94 (d, 2H, J = 8.2 Hz, $H_3$), 7.43 (d, 2H, J = 8.2 Hz, $H_2$), 4.61 (s, 2H, $H_8$), 4.44 (t, 2H, J = 9.5 Hz, $H_7$), 4.07 (t, 2H, J = 9.5 Hz, $H_6$). **$^{13}C$ {$^1H$} NMR** (75 MHz, $CDCl_3$) δ (ppm): 164.2 ($C_5$), 140.6 ($C_4$), 128.7 ($C_3$), 128.6 ($C_2$), 127.9 ($C_1$), 67.8 ($C_7$), 55.1 ($C_6$), 45.7 ($C_8$).

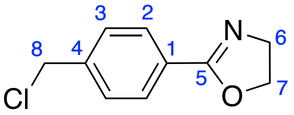

**L06N3: 2-(4-(azidomethyl)phenyl)-4,5-dihydrooxazole**

**L06** (500 mg, 2.55 mmol) was dissolved in 20 mL of MeCN and 1 mL of DMF was added. $NaN_3$ (250 mg, 1.5 eq.) was added to the stirred solution that was refluxed for 16 h. The reaction mixture was treated with 20 mL of water after it cooled down to room temperature, and the aqueous layer was extracted three times with dichloromethane. The organic layers were washed three times with $H_2O$, dried over solid magnesium sulfate, filtered and evaporated to give a yellow oil. $^1H$ NMR showed that it was the expected product with 100% conversion and no further purification needed. Only 1eq. of DMF was left to remove. The product was thus dissolved in $Et_2O$ and washed three times with brine. After drying with solid magnesium sulfate, filtering and removing the solvent, a yellowish oil was obtained again. It solidified after complete volatiles evaporation. Its final weight was 466 mg (90%).

**$^1H$ NMR** (300 MHz, $CDCl_3$) δ (ppm): 7.96 (d, 2H, $J$ = 8.2 Hz, $H_3$), 7.36 (d, 2H, $J$ = 8.1 Hz, $H_2$), 4.44 (t, 2H, $J$ = 9.5 Hz, $H_7$), 4.39 (s, 2H, $H_8$), 4.07 (t, 2H, $J$ = 9.5 Hz, $H_6$). **$^{13}C$ {$^1H$} NMR** (75 MHz, $CDCl_3$) δ (ppm): 164.3 ($C_5$), 138.7 ($C_4$), 128.8 ($C_3$), 128.1 ($C_2$), 127.9 ($C_1$), 67.8 ($C_7$), 55.1 ($C_6$), 54.5 ($C_8$). **IR (ATR):** ν(N=N=N) = 2081$cm^{-1}$

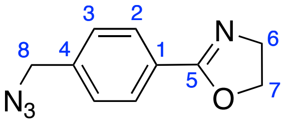

**IrN3: Chlorido(η$^5$-pentamethylcyclopentadienyl)(2-((4-azidomethyl)phenyl-κC$^2$)-4,5-dihydrooxazole-κN)Iridium(III)**

10 mL of anhydrous dichloromethane was degassed for 15 min over 4Å MS and $[Cp^*IrCl_2]_2$ (204 mg, 256 µmol, 1.0 eq.) was dissolved into the mixture. Activated NaOAc (126 mg, 1.537 mmol, 6.0 eq.) was added to the stirred solution and **L06N3** (109 mg, 539 µmol, 2.2 eq.) was added after 90 min stirring under argon atmosphere. The reaction was stirred 1 h at room temperature and became orange, it was filtered over a pad of dry Celite® 545 and evaporated to a semi-solid foam weighting 293 mg. This crude product was washed twice with

pentane and dissolved in 3 mL of dichloromethane, excess pentane was added and 150 mg of red precipitate containing mostly impurities were formed. The filtrate was a yellow solution from which part of the pure product could be crystallized by slow evaporation from a concentrated solution in dichloromethane (1 week) : 62 mg of pure orange crystals were obtained (8% yield).

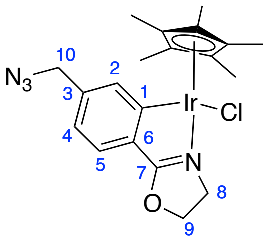

**$^1$H NMR** (300 MHz, $CD_2Cl_2$) δ (ppm): 7.72 (s, 1H, $H_2$), 7.41 (d, 1H, *J* = 7.7 Hz, $H_5$), 6.97 (dd, 1H, *J* = 7.7, 1.3 Hz, $H_4$), 4.93 – 4.76 (m, 2H, $H_9$), 4.46 (d, 1H, *J* = 13.6 Hz, $H_{10a}$), 4.36 (d, 1H, *J* = 13.6 Hz, $H_{10b}$), 4.10 (ddd, 1H, *J* = 12.5, 9.7, 7.8 Hz, $H_{8a}$), 3.91 (ddd, 1H, *J* = 12.5, 10.6, 9.2 Hz, $H_{8b}$), 1.76 (s, 15H, $H^{Cp^*}$). **$^{13}$C {$^1$H} NMR** (75 MHz, $CD_2Cl_2$) δ (ppm): 180.0 ($C_1$), 165.5 ($C_7$), 139.2 ($C_3$), 135.8 ($C_2$), 131.6 ($C_6$), 126.7 ($C_4$), 122.1 ($C_5$), 88.3 ($C^{Cp^*}$), 72.2 ($C_9$), 55.7 ($C_8$), 50.8 ($C_{10}$), 9.6 ($CH_3^{Cp^*}$). **HRMS** (ESI+): *m/z* calculated for $C_{20}H_{24}ClIrN_4O$: 529.1574; found: 529.1574 $[M-Cl]^+$. **IR (ATR):** ν(N=N=N) = 2091$cm^{-1}$

**IrN$_3$-DMSO: [Dimethylsulfoxide-(η5-pentamethylcyclopentadienyl)(2-((4-azidomethyl)phenyl-κC2)-4,5-dihydrooxazole-κN)Iridium(III)] Nitrate**

**IrN$_3$** (14 mg, 25 µmol) was dissolved in 3 mL of dichloromethane and $AgNO_3$ (4 mg, 1.0 eq.) was added to the solution. After 10 min, DMSO (2 µL, 1.1 eq.) was added and the mixture was stirred at room temperature for 16 h. The mixture went turbid over an hour, and AgCl (brown precipitate) appeared overnight. The suspension was filtered over a sintered glass frit (porosity 4) and the obtained clear yellow solution was evaporated to give 15 mg of a yellow solid. Residual cyclohexane was removed under vacuum to quantitatively give the product as a bright yellow solid.

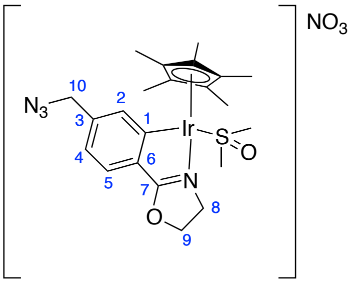

**$^1$H NMR** (400 MHz, $CD_2Cl_2$) δ (ppm): 7.63 (s, 1H, $H_2$), 7.58 (d, 1H, *J* = 7.8 Hz, $H_5$), 7.23 (dd, 1H, *J* = 7.8, 1.3 Hz, $H_4$), 5.26 – 5.14 (m, 1H, $H_{9a}$), 5.07 – 4.95 (m, 1H, $H_{9b}$), 4.46 (d, 2H, *J* = 1.08 Hz, $H_{10}$), 4.20 – 4.01 (m, 2H, $H_8$), 3.07 (s, 3H, $H_{DMSO}$), 2.63 (s, 3H, $H_{DMSO}$), 1.85 (s, 15H, $H^{Cp^*}$). **$^{13}$C {$^1$H} NMR** (101 MHz, $CD_2Cl_2$) δ (ppm): 182.1 ($C_1$), 156.0 ($C_7$), 142.3 ($C_3$), 135.9 ($C_2$), 131.0 ($C_6$), 129.0 ($C_4$), 125.2 ($C_5$), 96.8 ($C^{Cp^*}$), 73.6 ($C_9$), 55.0 ($C_8$), 52.0 ($C_{10}$), 46.7 – 44.5 ($CH_3^{DMSO}$), 9.4 ($CH_3^{Cp^*}$).

**phenyl-©-IrN$_3$: Chlorido(η$^5$-pentamethylcyclopentadienyl)(2-((4-(5-benzyl-1*H*-1,2,3-triazolyl-4-yl)methyl)phenyl-κC$^2$)-4,5-dihydrooxazole-κN)Iridium(III)**

**IrN$_3$** (278 mg, 492 µmol) was dissolved in 5 mL of DMSO for 10 min at 50°C using a water bath. The color of the solution became lighter leading to the formation a cationic DMSO adduct. Then, phenylacetylene (55 µL, 500 µmol) was added to the solution followed by a copper sulfate pentahydrate (250 mg, 1 mmol, 2.0 eq.) solution in water (1 mL). The reaction mixture turned green and translucid. Sodium ascorbate (600 mg, 3 mmol, 6.0 eq) was added and the reaction mixture turned yellow indicating the formation of Cu(I). The mixture was stirred 2 h at 50°C and concentrated under vacuum for one more hour to evaporate DMSO. The dark red crude product was dissolved in 5 mL of dichloromethane and 5 mL of water were added to the solution. The mixture was decanted and the aqueous layer was extracted twice with dichloromethane. The combined organic layers were washed twice with an aqueous 0.5 M EDTA solution before being evaporated to dryness, filtered through a pad of silica with dichloromethane, then eluted and washed with methanol. During this operation, the methoxo neutral product could have been formed. The most polar fraction was eluted in reverse phase preparative HPLC column using MeOH to give ca. 5 mg of pure product as pale-yellow crystals.

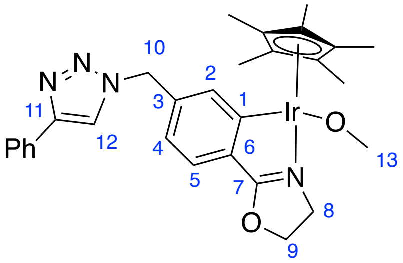

**$^1$H NMR** (400 MHz, $CD_2Cl_2$) δ (ppm): 7.84 – 7.73 (m, 3H, 2$H^{Ph}$ + $H_2$), 7.49 (s, 1H, $H_{12}$), 7.42 – 7.32 (m, 5H, $H_5$ + $H^{Ph}$), 7.32 (m, *J* 1H, $H_5$), 7.05 (dd, *J* = 7.5, 1.5 Hz, 1H, $H_4$), 5.72 (d, *J* = 15.2 Hz, 1H, $H_{10a}$), 5.62 (d, *J* = 15.2 Hz, 1H, $H_{10b}$), 4.95 (q, *J* = 8.6 Hz, 1H, $H_{9a}$), 4.79 (q, *J* = 9.5 Hz, 1H, $H_{9b}$), 4.05 – 3.91 (m, 2H, $H_8$), 2.21 (s, 3H, $H_{13}$), 1.73 (s, 15H, $H^{Cp}$). **$^{13}$C {$^1$H} NMR** (101 MHz, $CD_2Cl_2$) δ (ppm): 181.1 ($C_1$), 161.0 ($C_7$), 138.8 ($C_3$), 135.1 ($C_{12}$), 132.8 ($C_2$), 131.5 ($C_6$), 131.3 ($C_{11}$), 129.3 (2$C^{Ph}$), 128.6 ($C^{Ph}$), 127.4 ($C_5$), 126.1 (2$C^{Ph}$), 122.1 ($C_4$), 120.5 ($C^{Ph}$), 93.7 ($C^{Cp^*}$), 72.6 ($C_9$), 54.4 ($C_8$), 51.9 ($C_{10}$), 50.9 ($C_{13}$), 9.4 ($CH_3^{Cp^*}$). **HRMS** (ESI+): *m/z* calculated for $C_{28}H_{30}IrN_4O$: 631.2043; found: 631.2039 $[M-Cl]^+$(-0.7 ppm).

**L08: 4,4-dimethyl-2-(4-(chloromethyl)phenyl)-4,5-dihydrooxazole**

To a stirred solution of p-chloromethyl-benzoyl chloride (1.046 g, 5.53 mmol) in 30 mL of dichloromethane, 2-amino-2-methyl-1-propanol (1.233 g, 2.5 eq, 13.83 mmol) was added portion wise and the reaction was left stirring for 1 h. 10 mL of $H_2O$ were added to the mixture and the aqueous layer was extracted three times with ethyl acetate. Organic layers were dried over magnesium sulfate and evaporated to give 1.410 g of a white powder. ($^1$H NMR analysis) This intermediate product (M=241.72 g/mol) was dissolved in 20 mL of dichloromethane and a large excess of thionyl chloride (1.7 mL, 4.0 eq) was added dropwise. The solution was left stirring at room temperature overnight. The suspension was poured into ice and the aqueous layer was neutralized with 10 g of solid sodium bicarbonate. The neutral aqueous layer was then extracted 3 times with ethyl acetate and the combined organic layers were washed with water, dried over magnesium sulfate and evaporated to give 1.282 g of colorless oil. The resulting crude product was dissolved in 25 mL of THF and 60% NaH in mineral oil (332 mg, 1.5 eq) were added portionwise. The resulting brown reaction mixture was stirred during 2 h at room temperature. The reaction was stopped with a saturated aqueous ammonium chloride solution. The mixture was decanted and the aqueous layer was extracted three times with dichloromethane. The combined organic layers were washed with water, dried over solid magnesium sulfate, filtered and evaporated to give 1.211 g of crude product as a yellow semi-solid.

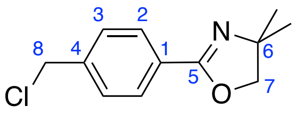

**$^1$H NMR** (400 MHz, $CD_2Cl_2$) δ (ppm): 7.90 (d, *J* = 8.3 Hz, 2H, $H_2$), 7.43 (d, *J* = 8.7 Hz, 2H, $H_3$), 4.63 (s, 2H, $H_8$), 4.10 (s, 2H, $H_7$), 1.34 (s, 6H, $H_{Me2}$). **$^{13}$C {$^1$H} NMR** (101 MHz, $CD_2Cl_2$) δ (ppm): 161.6 ($C_5$), 141.0 ($C_4$), 129.0 (3C, $C_1$, $C_2$, $C_3$), 79.6 ($C_7$), 68.2 ($C_6$), 46.3 ($C_8$), 28.7 ($C_{Me2}$). **HRMS** (ESI+): *m/z* calculated for $C_{12}H_{14}ClNOH$: 224.0837; found: 224.0838 $[M+H]^+$ (+0.6 ppm).

#### L08N$_3$: 4,4-dimethyl-2-(4-(azidomethyl)phenyl)-4,5-dihydrooxazole

The crude product **L08** was dissolved in 10 mL of acetonitrile containing 0.5 mL of dimethylformamide, 100 mg of sodium azide (1.7 eq) were added and the reaction mixture was refluxed during 14 h. 10 mL of $H_2O$ were added to the mixture and the aqueous layer was extracted twice with dichloromethane. The organic layers were washed twice with water, dried over magnesium sulfate, and evaporated *in vacuo* to give 173 mg of pure product as a colorless oil (yield: 82%)

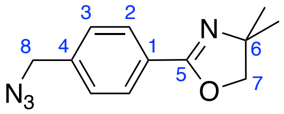

**$^1$H NMR** (400 MHz, $CD_2Cl_2$) δ (ppm): 7.92 (d, *J* = 8.3 Hz, 2H, $H_2$), 7.37 (d, *J* = 8.6 Hz, 2H, $H_3$), 4.40 (s, 2H, $H_8$), 4.10 (s, 2H, $H_7$), 1.34 (s, 6H, $H_{Me2}$). **$^{13}$C {$^1$H} NMR** (101 MHz, $CD_2Cl_2$) δ (ppm): 161.7 ($C_5$), 139.1 ($C_4$), 129.0 ($C_3$), 128.8 ($C_1$), 128.5 ($C_2$), 79.7 ($C_7$), 68.2 ($C_6$), 54.9 ($C_8$), 28.7 ($C_{Me2}$). **HRMS** (ESI+): *m/z* calculated for $C_{12}H_{14}N_4OH$: 231.1240; found: 231.1242 $[M+H]^+$ (+0.7 ppm).

#### Ir2N$_3$: Chlorido(η5-pentamethylcyclopentadienyl)(2-((4-azidomethyl)phenyl-κC2)-4,5-dihydro-2,2-dimethyloxazole-κN)Iridium (III)

$[Cp^*IrCl_2]_2$ (726 mg) was dissolved in dichloromethane (50 mL) and sodium acetate (448 mg, 6.0 eq, mortar, anhydrous) was added to the red solution. The mixture was stirred at room temperature in the dark during 2 h, by which it turned to orange. **L08N$_3$** (2.2 eq, 456 mg) dissolved in dichloromethane (3 mL) was added and the mixture quickly turned brown. The resulting brown mixture was stirred at room temperature during 45 min and filtered over a pad of Celite® 545. The filtrate was evaporated to give a residue which was triturated with cyclohexane. The resulting solid was redissolved in dichloromethane (3 mL) and cyclohexane (60 mL) was added to the homogeneous solution. The resulting red precipitate was discarded and the filtrate was evaporated to give ca. 750 mg of crude product. This crude product containing 45:55 ligand:complex according to $^1$H NMR to a conversion of 52%, was purified over an alumina column (neutral) using Cy/AcOEt : 9/1 as eluant. 150 mg of pure product were obtained and crystallized by slow evaporation (253 µmol, 14% yield)

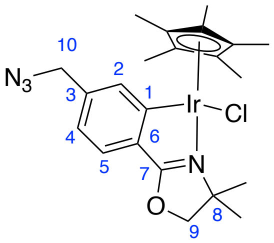

**$^1$H NMR** (400 MHz, $CD_2Cl_2$) δ (ppm) : 7.70 (d, *J* = 1.6 Hz, 1H, $H_2$), 7.42 (d, *J* = 7.7 Hz, 1H, $H_5$), 6.96 (dd, *J* = 7.8, 1.6 Hz, 1H, $H_4$), 4.56 – 4.34 (m, 4H, $H_9$,$H_{10}$), 1.77 (s, 15H, $H^{Cp*}$), 1.49 (d, *J* = 3.0 Hz, 6H, $H_{Me2}$). **$^{13}$C {$^1$H} NMR** (101 MHz, $CD_2Cl_2$) δ (ppm): 177.8 ($C_1$), 164.2 ($C_7$), 139.1 ($C_3$), 135.6 ($C_2$), 133.1 ($C_6$), 126.6 ($C_4$), 122.1 ($C_5$), 88.6 ($C^{Cp*}$), 83.5 ($C_9$), 68.1 ($C_8$), 55.8 ($C_{10}$), 28.8 ($C_{Me2}$), 26.6 ($C_{Me2}$), 10.2 ($CH_3^{Cp*}$). **HRMS** (ESI+): *m/z* calculated for $C_{22}H_{28}IrN_4O$: 557.1887; found 557.1887 $[M-Cl]^+$ (0.0 ppm).

### 2. Supplementary figures-associated protocols and instrumentation

**X-Ray crystal structure determinations (Figure S1)**. For the three compounds **Ir2**-DMSO, **IrN3** and **Ir2N3**, a single crystal was selected, mounted onto a cryoloop and transferred into a cold nitrogen gas stream. Intensity data were collected with a Bruker Kappa APEX-II CCD diffractometer using a graphite-monochromated Mo-Kα radiation (λ = 0.71073 Å). Data collection were performed at 200K, with the Bruker APEXIII suite. Unit-cell parameters determinations, integrations and data reductions were carried out with SAINT program. SADABS was used for scalings and absorption corrections. The structures were solved with SHELXT[15] and refined by full-matrix least-squares methods with SHELXL[16] using Olex2 software package[17]. All non-hydrogen atoms were refined anisotropically. These structures were deposited at the Cambridge Crystallographic Data Centre with numbers CCDC from 2205966 to 2205968 respectively and can be obtained free of charge via http://www.ccdc.cam.ac.uk.

**(Chemical) Mass spectrometry analysis (Figure S2 and Supplementary Tables 2-3).** ESI-MS experiments were carried out using LTQ-Orbitrap XL from Thermo Scientific (Thermo Fisher Scientific, Courtaboeuf, France) and operated in positive ionization mode, with a spray voltage at 3.6 kV. Sheath and auxiliary gas were set at a flow rate of 45 and 15 arbitrary units (a.u.), respectively. Applied voltages were 20 and 70 V for the ion transfer capillary and the tube lens, respectively. The ion transfer capillary was held at 275°C. Detection was achieved in the Orbitrap with a resolution set to 100,000 (at m/z 400) and a m/z range between 200-1000 in profile mode. Spectrum was analyzed using the acquisition software XCalibur 2.1 (Thermo Fisher Scientific, Courtaboeuf, France). The automatic gain control (AGC) allowed accumulation of up to $2.10^5$ ions for FTMS scans, Maximum injection time was set to 500 ms and 1 µscan was acquired. 5 or 20 µL were injected using a Thermo Finnigan Surveyor HPLC system (Thermo Fisher Scientific, Courtaboeuf, France) with a continuous infusion of methanol or methanol/water mixture at 100 µL/min.

***In vitro* measurement of NADH oxidation and $H_2O_2$ production (Figure S3).** The iridium complexes (10 µM) and NADH (100 µM) were incubated for 150 min in 5 mM Phosphate Buffer pH 7.4/MeOH 99:1. Mixtures were analyzed over time by UV-Vis spectroscopy at 20°C. The concentration of NADH was calculated from triplicates using the extinction coefficient $\varepsilon_{340}$ = 6200 $M^{-1}cm^{-1}$. *Amplex Red® HRP-linked enzymatic assay:* the iridium complexes (10 µM) or $H_2O_2$ standards (150 nM - 5 µM), the Amplex substrate (5 µM), HRP (1 U/mL) and NADH (100 µM) containing SOD (40 U/mL) were dispensed in a black 96-well plate. Fluorescence (exc. 520 nm; em. 595 nm) was recorded every 5 min during 150 min from triplicate samples to calculate the concentration of $H_2O_2$ over time.

**Clonogenicity assay (Figure S4).** Three 6-well plates were seeded with 400 HeLa cells per well in 2 mL of complete medium. The next day (D2), cells were treated with **Ir2** at 1.25, 2.5 or 5 µM or the equivalent amounts of DMSO vehicle, in triplicate. Final serum concentration was 5 %. At D4, control colonies were of about 8 to 32 cells, cells treated with 5 µM **Ir2** did not form colonies. At 2.5 µM, colonies were of about 2 to 8 cells and at 1.25 µM, colonies were of about 4 to 16 cells. At D5, cells were fixed in 1 mL of PFA 4%/PBS and stained for 30 min at room temperature with 1 mL of 0.05 % Crystal Violet in PBS. Wells were washed three times with PBS and dried. In a second experiment, a 35-mm dish was first seeded with 80,000 HeLa cells in 2 mL of complete HeLa medium. The next day, cells were exposed to 5 µM **Ir2** for 24 h in medium containing 5% FBS. The dish was washed three times with PBS and adherent cells were trypsinized and pelleted. Afterwards, the clonogenicity experiment was reproduced using this cell sample or untreated cells.

**Phospho-H2AX immunofluorescence assay (Figure S5).** HeLa and hTERT-RPE1 cells were seeded on 14 mm coverslips (25,000 cells per $cm^2$) and incubated in complete medium at 37°C 5% $CO_2$ in a humidified atmosphere. After 24 h, cells were treated with the indicated drug or the DMSO vehicle at the indicated time. Cells were fixed with 4% PFA in PBS and washed 3 times with PBS. Samples were blocked and permeabilized using 3% BSA and 0.5 % Triton X-100 in PBS for 30 min at room temperature. Coverslips were incubated overnight with a primary antibody (1:100 Rabbit anti-phospho-H2AX (Ser 139) (Upstate Technology #05-636 clone JBW301)) in PBS 3 % BSA at 4°C, then for 1 h at room temperature with a fluorescent secondary antibody (Invitrogen® goat anti-rabbit Ig-G (H+L) Alexa Fluor 488 1:500 in PBS) and finally with 5 µg/mL Hoechst 33342 in PBS for 10 min. Coverslips were mounted using Invitrogen® Prolong Diamond mounting medium. **ICP-OES analysis.** Briefly, 10-mm culture dishes were seeded with 250,000 HeLa or hTERT-RPE1 cells in complete medium. After two days, each dish was washed with warm PBS and cells were incubated with 10 mL of serum-free medium containing 5 or 10 µM **Ir2** for 30 min. Supernatants were collected, cells were washed twice with PBS and harvested. Cell samples were digested *ca.* 1 h in ultrasonic bath with concentrated $HNO_3$ and 1% Triton X-100, and the extract was diluted with water to a final volume of 5 mL and 2% $HNO_3$. Samples were filtered through 0.2 µm nylon membrane filters and analysed by ICP-OES for content in iridium (212.681 nm) using an ICP-OES 5100 SVDV Agilent analyser. Iridium calibration standards (8, 16, 31, 63, 125, 250, 500 and 1000 ppb in 2% $HNO_3$) were prepared using a 10 ppm PlasmaCAL SCP Science® multi-element stock solution. Measurements were performed in triplicate.

**CuAAC of ethynyl-$PEG_4$-biotin on a cell suspension (Figure S6).** $2.10^6$ hTERT-RPE1 cells were seeded in 4 dishes (100 mm) and cultured overnight in 10 mL of complete medium. The next day, cells were washed with PBS and treated for 1 h with either the DMSO vehicle or **$IrN_3$** (0.1, 1 or 10 µM) in serum free medium. After treatment, cells were washed with PBS and detached with 0.05% Trypsin-EDTA, the dissociation reagent was diluted with PBS containing 1 % BSA. Cells were pelleted and washed 3 times with PBS at 4°C, then permeabilized with 1 % Saponin in PBS containing protease inhibitors for 15 min at 4°C. Next, pellets were incubated with the click cocktail (10 µM ethynyl-$PEG_4$-biotin, Lumiprobe) for 30 min at room temperature, washed three times with the permeabilization buffer and lysed in 100 µL of RIPA buffer per sample for 30 min at 4°C. Lysates were cleared by centrifugation at 14,000 rpm and fractionated by SDS-PAGE in 10 % polyacrylamide. Proteins were transferred onto a 0.2 µm nitrocellulose membrane stained with Ponceau S, blocked with 5 % BSA in TBS 1 % Tween-20 for 1 h. After 1 h incubation with Streptavidin-HRP (1:1000) in TBS-t, three washes with TBS-t and one wash with distilled water, the membrane revelation was carried out using the BioRad ECL kit.

**CuAAC of ethynyl-5-FAM on fixed cells (Figure S8).** $20.10^3$ hTERT-RPE1 cells were seeded on 14 mm coverslips (#1.5) in a 12-well plate and cultured in complete medium. The next day, cells were washed and treated for 1 h with **$IrN_3$** at 1, 5 or 10 µM in serum-free media. Cells were fixed with 4 % PFA in PBS and washed three times with PBS. Specimens were blocked and permeabilized using PBS containing 3 % BSA and 0.5 % Triton X-100 during 30 min at room temperature. Coverslips were washed with PBS and incubated with the click cocktail containing 4.5 µM ethynyl-5-FAM (Lumiprobe) during 30 min. Finally, samples were washed 3 times with PBS and stained with 1 µg/mL Hoechst 33342 and Alexa Fluor® 633-Phalloidin in PBS during 30 min. Coverslips were washed with PBS and mounted in Prolong Diamond mounting medium. Acquisitions were performed on an inverted SP5 Leica confocal system, using the 63x oil immersion NA 1.4 objective and the following excitation laser lines : 405 nm, 488 nm, 633 nm.

**Heat Shock Proteins 70 activity assays of *in vitro* (Figure S10).** ATPase assay: Steady-state ATPase activity of HSP70-1 (HSPA1B) was reported by PK-LDH catalyzed oxidation of NADH as described earlier (see main text) in 100 µL on 96 well plates. The reaction buffer was changed to 50 mM Hepes pH=7.5, 100mM KCl, 5mM ATP; 5 mM $MgCl_2$. Luciferase refolding assay: QuantiLum® Recombinant Luciferase (Promega) was diluted (55 µM) and chemically denatured into denaturation buffer (6M Guanidium/HCl; 1 mM DTT) during 30 minutes at room temperature. The luciferase is then diluted 125 times (0.4 µM) into renaturation buffer (20 mM Tris pH=7.5; 50 mM KCl; 5 mM $MgCl_2$; 1 mM DTT) and incubated with HSP70 (10 µM), DNAJB1 (2 µM) and eventually with STIP1 (0.5 µM) and HSP90β (2 µM). Luciferase activity was measured after mixing of 5 µL sample with 120 µL of luciferin buffer (20 mM Tris pH=7.5; 200 µM luciferin; 0.5 mM ATP; 10 mM $MgCl_2$). The chemiluminescence was measured with a ClarioStar (BMG Labtech). The percentage of refolding was evaluated relatively to the activity of native luciferase.

**Polymerization of actin microfilaments *in vitro* (Figure S11).** GAB: General Actin Buffer (5 mM Tris-HCl pH 8.0, 0.2 mM $CaCl_2$ supplemented with 0.2 mM ATP and 0.5 mM DTT). Purified actin and polymerization buffer purchased from Cytoskeleton® were resuspended, aliquoted and snap-frozen according to the supplier's protocol. Actin monomer stock solution (32 µL, 320 µg) was diluted in 800 µL of GAB. This solution was incubated for 1 h at 4°C, and centrifuged (15 min at 14'000 rpm) to eliminate already polymerized actin and aggregates. The classic procedure is: adding the ATP-containing polymerization buffer (100X) and incubating for 1 h at room temperature, then centrifugate for 1 h at 100,000 g. Experiments: **A**- Polymerization buffer is replaced with 100 mM Tris-HCl pH 7.5 (no ATP added, negative control); **B**- Classic procedure with 15 min pre-incubation of the actin monomer with 1 or 10 µL DMSO; **C**- 15 min of pre-incubation with 10 or 100 µM **$Ir2N_3$**; **D**- 15 min post-incubation in the same conditions after polymerization. Supernatants (Sn) were separated from pellets (Pl). Pellets of microfilaments were resuspended in 180 µL GAB and left for 1 h at 4°C to depolymerize the formed microfilaments. Samples were incubated for 5 min at 95°C with 5X denaturation buffer, then migrated on a 10-well 4-20% polyacrylamide gel by SDS-Page. The gel was stained using Instant-Blue (abcam ab119211) and 30 µL (equivalent to 10 µg of actin) were deposed on each lane.

### 3. Heat Shock Proteins cloning and purification

**Plasmids**. The pRSF-DUET1-Smt3m plasmid was created by modification of the pRSF-DUET-1 (Novagen) in order to express residues 1-98 of *S. cerevisiae* protein Smt3 fused to the 6-HIS tag[18]. The initial *Sfo1* restriction site was then removed and a new *SfoI* site was created by mutation of the Smt3 Gly97 codon GGT to GGC and the introduction of the sequence 5'-GCC-3' in 3' of this modified codon using a cassette amplified by PCR (primers: RSF_SFO1_fwd & RSF_SFO1_rev) and cloned using *Sfo1* and *BamH1* restriction enzymes. The human gene of stress-inducible HSP70-1, (HSPA1A, UniprotKB: P0DMV8) was amplified by PCR (primers: HSP701_fwd & HSP701_rev) from HSPA1A cDNA (DNASU: HsCD00076277) and cloned using Gibson assembly and the restriction enzymes *SfoI* and *NotI* into the pRSF-DUET-1-Smt3m. The human gene of STIP1 (UniprotKB: P31948) was amplified by PCR (primers: STIP1_fwd & STIP1_rev) from STIP1 cDNA (DNASU: HsCD00519528) and cloned into the pRSFDuet-1-Stm3m plasmid in fusion with an amino-terminal His6-Smt3 tag, as previously described using Gibson assembly and restriction enzymes *SfoI* and *HindIII*. The human gene of DNAJB1 (UniprotKB : P25685) from a pDONR plasmid (DNASU: HsCD00005559) was amplified by PCR (primers: DNAJB1_fwd & DNAJB1_rev) and cloned into a pCDF-DUET1 plasmid (Novagen) using restriction enzymes *BamHI* and *NotI*. The human gene of HSP90β (UniprotKB: P08238) cloned into the plasmid pRSETa was kindly provided by Prof. L. H. Pearl (University of Sussex). All plasmids were validated by full sequencing of insert and flanking regions (Eurofins).

**Bacterial culture**. All plasmids were transformed into chemicompetent *E. coli* Rosetta™(DE3)pLysS cells (Novagen). The cells were grown in 2xYT medium at 37°C until the $OD_{600nm}$ reached 0.8, except for HSP90β where TB medium was used. The expression was then induced by 0.4 mM of IPTG at 18°C during 24 hours (for HSP90β and STIP1) or at 30°C during 4 hours (for HSP70 and STIP1). Cell pellets were collected by 4,000 g centrifugation for 15min.

**Purification of recombinant proteins**. The cell pellets were resuspended in 50 mL lysis buffer (25 mM Tris-HCl pH=8.0; 500 mM NaCl; 20 mM imidazole pH=8.0) supplemented with SIGMAFAST™ antiprotease (SIGMA) and DNAase . Cells were then disrupted with a French Press at 16,000 PSI (1.1 kbar). Crude lysates were clarified by centrifugation at 15,000 g for 20 min and loaded on Ni-NTA resin on Äkta FPLC (Cytiva). Bound proteins were eluted over a 8 CV linear gradient from 20 mM to 500 mM imidazole. Peaks containing the protein of interest, as assessed by SDS-PAGE, were pooled and submitted to subsequent purification steps, according to the protein, as described bellow.

HSP90-β protein was diluted into 25 mM Tris-HCl pH=8.0; 50 mM NaCl and injected on a Q-Sepharose Fast Flow resin (Cytiva). Bound proteins were eluted over a 13 CV linear gradient from 50 mM to 500 mM NaCl. Peaks containing HSP90-β, as assessed by SDS-PAGE, were further subjected to size-exclusion chromatography on a HiLoad 26/60 Superdex 200 PG column (Cytiva) and eluted with buffer containing 25 mM Tris-HCl pH=7.5; 150 mM NaCl; 0.5 mM EDTA pH=8.0. Peaks containing HSP90-β, as assessed by SDS-PAGE, were pooled and concentrated before snap freezing in liquid nitrogen and storage at -80°C.

His6-Smt3-HSP70-1 protein was cleaved by Ulp1[5] at enzyme:substrate mass ratio of 1:100 at 4°C overnight. Cleaved HSP70 was separated from Smt3 and Ulp1 and was collected into the flow through of NiNTA. Proteins were further subjected to size-exclusion chromatography on a HiLoad 26/60 Superdex 200 PG column (Cytiva), and eluted with buffer containing 25 mM HEPES/KOH pH=7.5; 150 mM KCl; 5 mM $MgCl_2$. Peaks containing proteins of interest, as assessed by SDS-PAGE, were pooled and concentrated before being snap-frozen and stored at -80°C.

His6-Smt3-STIP1 was cleaved by Ulp1 at enzyme:substrate mass ratio of 1:100 at 4°C overnight. Proteins were further subjected to size-exclusion chromatography on HiLoad 26/60 Superdex 200 PG (Cytiva), and eluted with buffer containing 25 mM Tris-HCl pH=7.5; 150 mM NaCl; 0.5 mM EDTA pH=8.0. The remaining non-cleaved protein was separated on a Ni-NTA resin and the flow through was collected. Peaks containing STIP1, as assessed by SDS-PAGE, were pooled and concentrated before snap freezing and storage at -80°C.

This His6-DNAJB1 protein was cleaved by TEV protease at enzyme:substrate mass ratio of 1:100 at 4°C overnight. Proteins were further subjected to size-exclusion chromatography on a HiLoad 26/60 Superdex 75 PG column (Cytiva), and eluted with a buffer containing 25 mM Tris-HCl pH=7.5; 150 mM NaCl; 1 mM DTT; 0.5 mM EDTA pH=8.0. The remaining non-cleaved protein was separated by Ni-NTA resin and the flow through was collected. Peaks containing DNAJB1, as assessed by SDS-PAGE, were pooled and concentrated before snap freezing and storage at -80°C.

| **Oligonucleotide** | **Source** | **Identifier** |
|---|---|---|
| RSF_SFO1_fwd :<br>5’ TCCAGGGTGGTTTTTCTTTTCACCAGTG 3’ | Eurofins | RSFmsfo_f |
| RSF_SFO1_rev :<br>5’ TTCGGATCCggcgCCAATCTGTTCTCTGT 3’ | Eurofins | RSFmsfo_r |
| HSP701_fwd:<br>5’ AGGCTCACAGAGAACAGATTGGCGGCGCCAAAGCCGCGGCGATCGGC 3’ | Eurofins | HSP70-1_f2 |
| HSP701_rev:<br>5’ TGTTCGACTTAAGCATTATGCGGCCGCCTACCCCATCAGGATGGCCGCCT 3’ | Eurofins | HSP70-1_r4 |
| STIP1_fwd:<br>5’ GGGGAGCAGGTCAATGAGCTGAAG 3’ | Eurofins | STIP1_f1 |
| STIP1_rev:<br>5’ AAGCTTTCACCGAATTGCAATCAGACC 3’ | Eurofins | STIP1_r1 |
| DNAJB1_fwd:<br>5’ CACGGGATCCGGAAAACCTGTACTTCCAGGGCGGTAAAGACTACTACCAGACG 3’ | Eurofins | DNAJB1_f1 |
| DNAJB1_rev:<br>5’ CGATGCGGCCGCCTACTATATTGGAAGAACCTGCTCAAGTA 3’ | Eurofins | DNAJB1_f2 |

### II. Supplementary figures, videos and tables

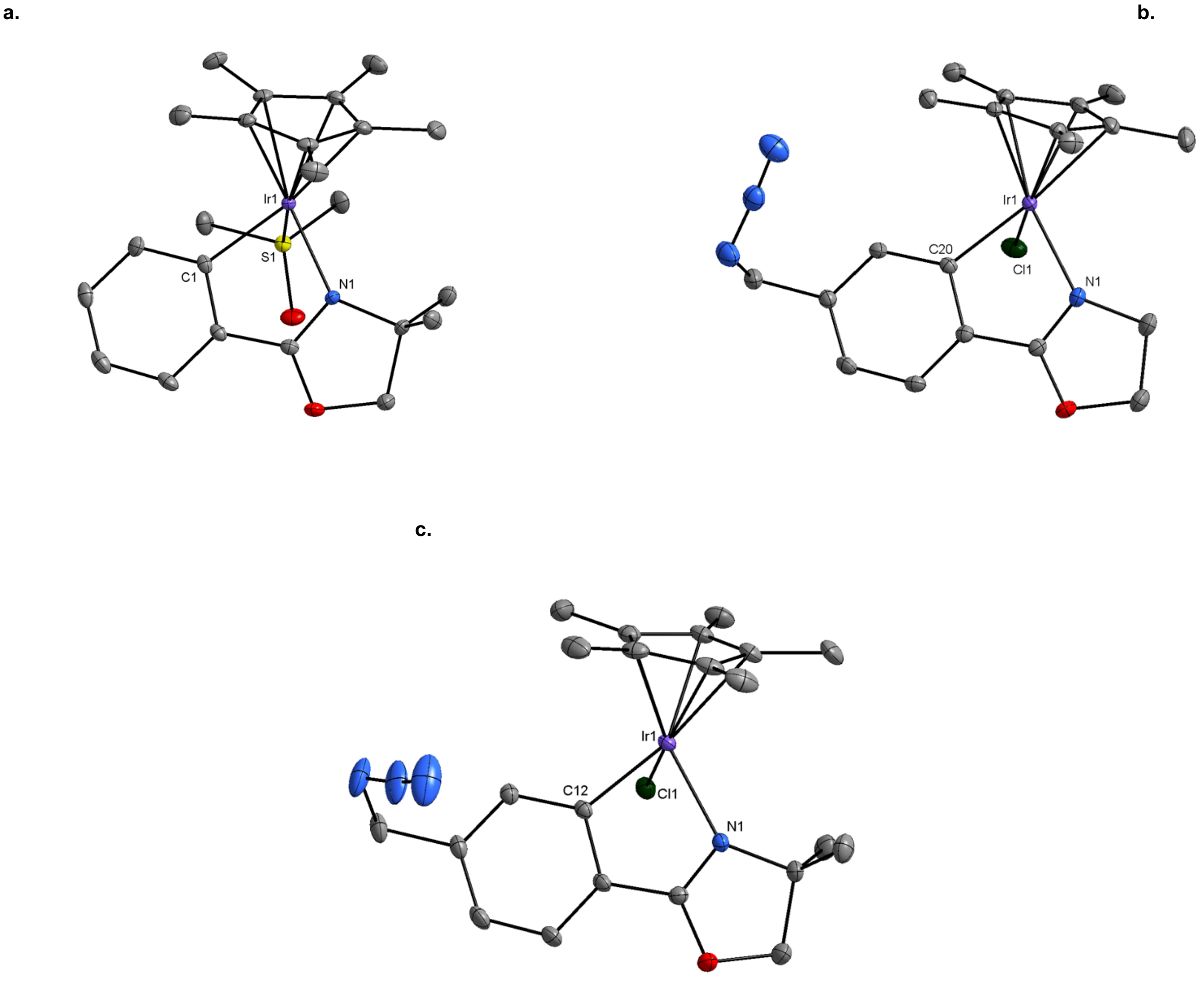

**Figure S1.** Crystal structure representations of the three half-sandwich complexes

All non-hydrogen atoms were refined anisotropically and ellipsoids are drawn with 30% probability. Hydrogen atoms are omitted for clarity. **a.** Cationic adduct [**Ir2**-DMSO]$^+$. The hexafluorophosphate counter-anion is omitted and only one enantiomer is shown. **b.** Neutral **IrN$_3$** complex. **c.** Neutral **Ir2N$_3$** complex. Only one enantiomer is represented, one disordered co-crystallized molecule of solvent (dichloromethane) is omitted, only the major position for the azido function is represented (occupancy=0.8).

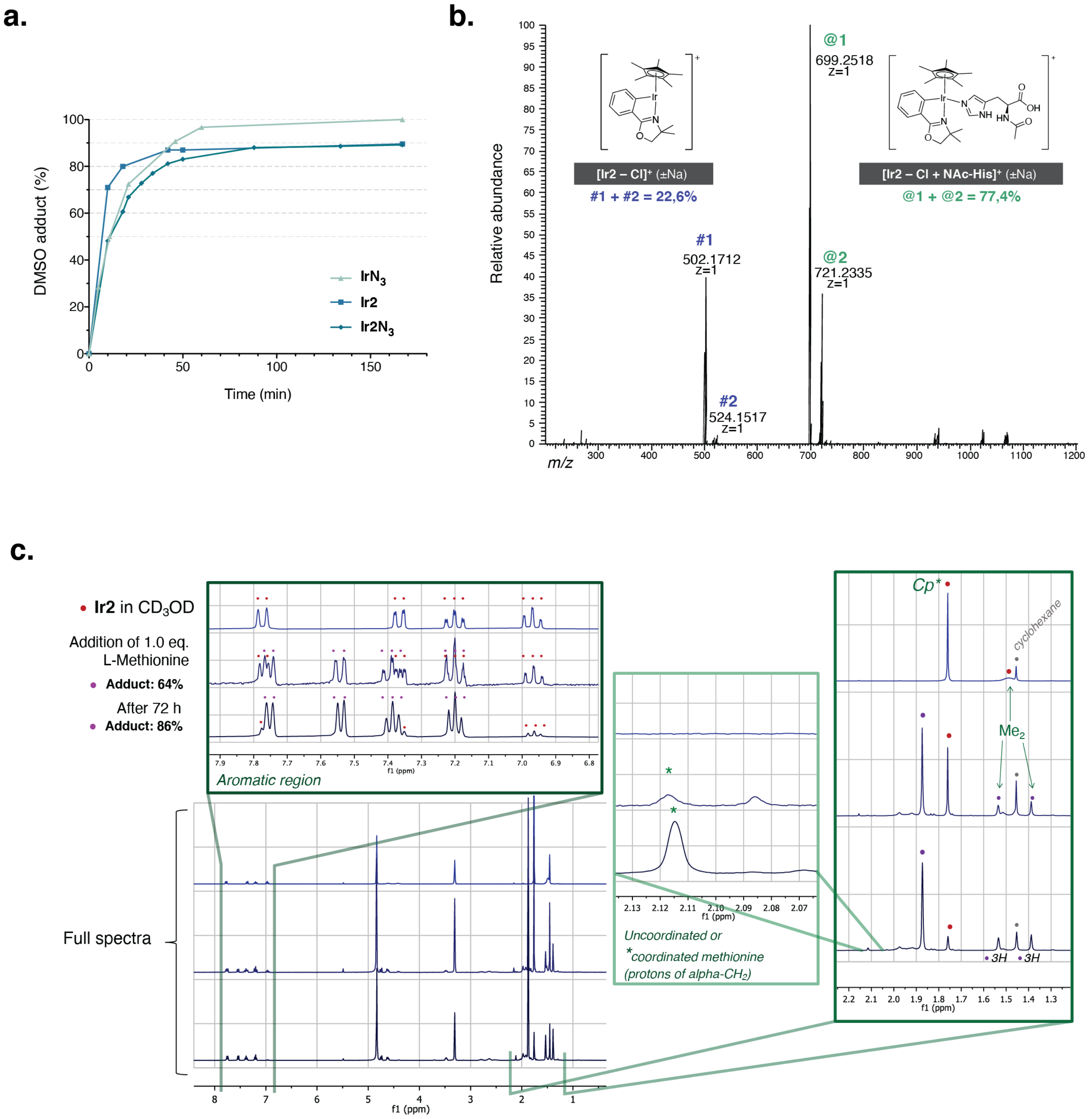

**Figure S2. Ir2** forms adducts with DMSO, Histidine and Methionine

**a.** Comparison of the rates of solvolysis of the three complexes in DMSO-d6 by integration of the characteristic $^1$H NMR peaks attributed to the 15 H of the Cp* ligand in the neutral and cationic forms. All three complexes undergo a fast ligand exchange in DMSO.
**b.** ESI-HRMS spectrum obtained after incubation of a solution of **Ir2** with an excess N-Acetyl-Histidine, relative abundance of the peaks representative of the attributed structures.
**c.** $^1$H NMR spectra (MeOD) of **Ir2** before and after incubation with a stoichiometric amount of L-Methionine

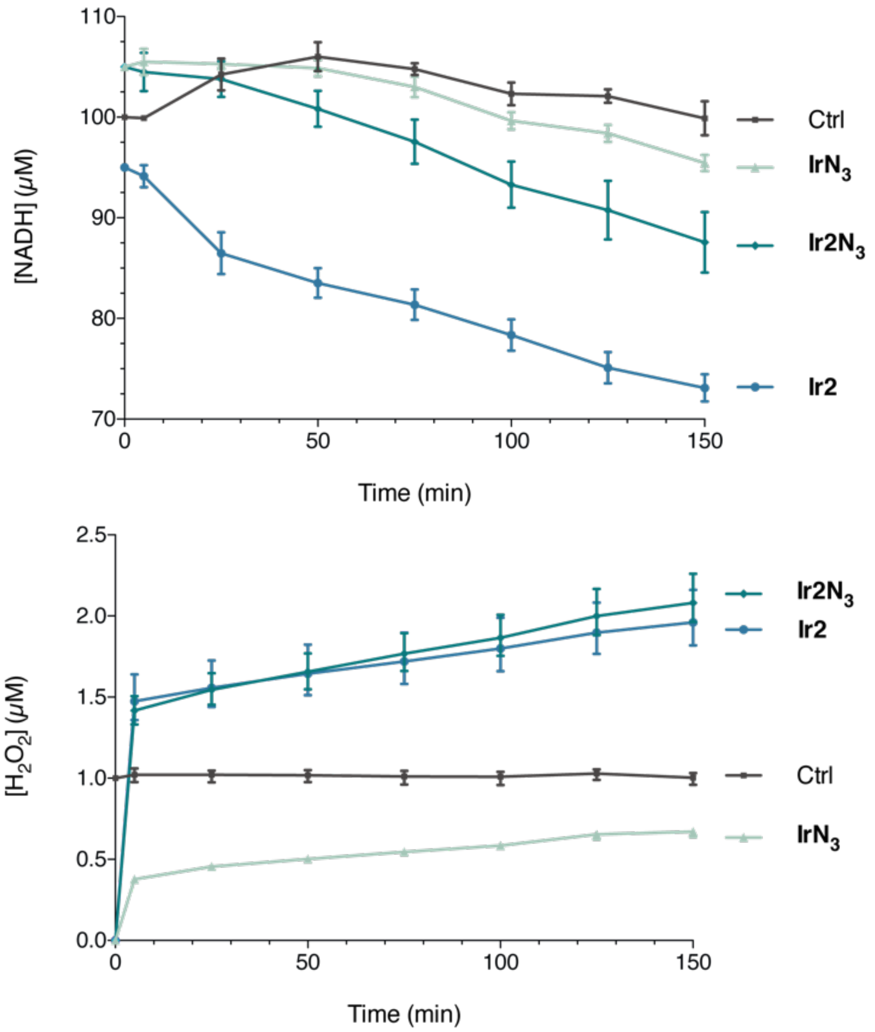

**Figure S3. Ir2** and **Ir2N₃** catalyze the oxidation of NADH and the production of $H_2O_2$ *in vitro*

Comparison of the catalytic activity of each complex. Consumption of NADH and production of $H_2O_2$ in model physiological environment, plotted as mean ± SEM of three independent experiments performed at 20 °C in 5 mM phosphate buffer pH 7.4 containing 1% MeOH in the presence of 10 mol% iridium complex. Solutions of known concentrations of substrates were used as control experiments.

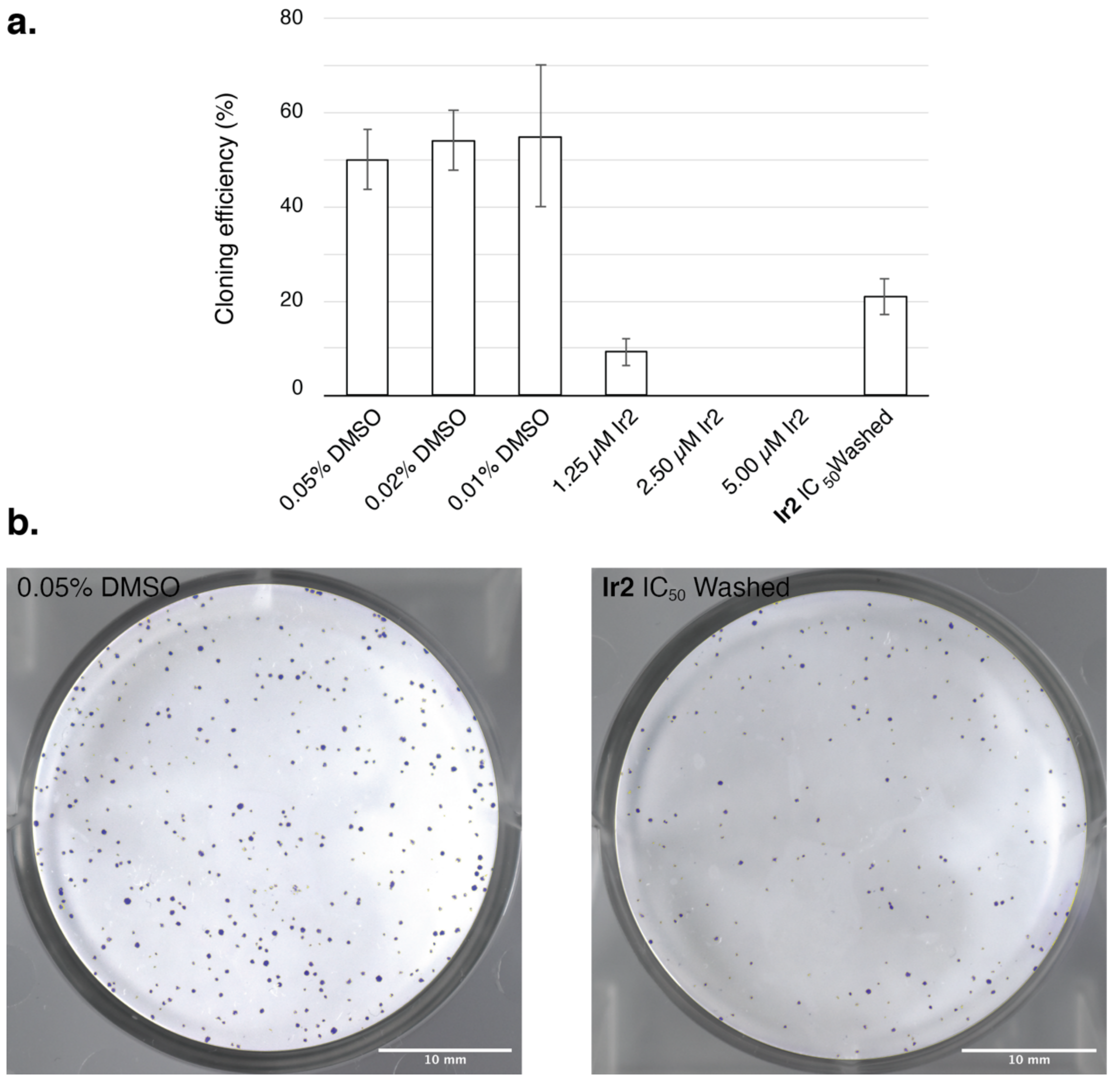

**Figure S4.** Clonogenicity assay on HeLa cells

**a.** Cloning efficiency, *i.e.* percentage of cells forming colonies containing more than 50 cells after 6 days of culture with the indicated concentration of **Ir2** or DMSO vehicle. **b.** Photographs of a representative replicate after fixation and crystal violet staining. For the condition “**Ir2** $IC_{50}$ Washed”, cells were treated with 4 µM **Ir2** for 24 h, trypsinized and used for the clonogenicity assay in **Ir2**-free medium to probe cell recovery.

**a.**

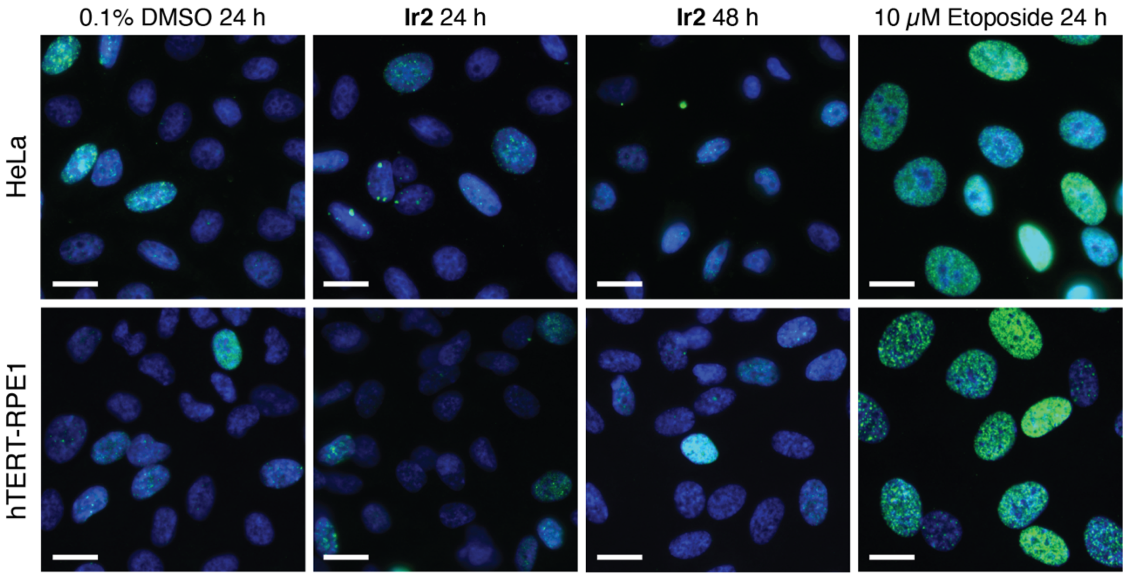

**b.**

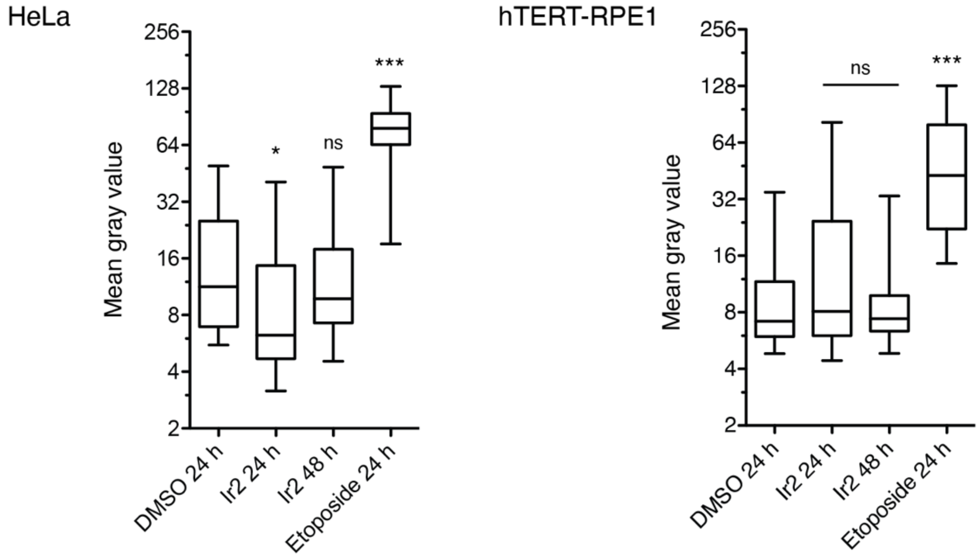

**Figure S5. Ir2** does not induce severe DNA damage in HeLa and hTERT-RPE1 cells.

**a.** Representative images of cells immuno-labelled for phosphorylated histone H2AX (green) to reveal DNA damage foci, DNA stained with Hoechst 33342 (blue). HeLa and hTERT-RPE1 cells were cultivated for the indicated time in the presence of 4 and 8 μM **Ir2** respectively. The topoisomerase inhibitor etoposide was used as a positive control, inducing a lethal replication stress. Scale bar, 10 μm.
**b.** Plot showing mean gamma-H2AX intensity per nucleus (±SD, error bars: 95% percentile) calculated over N=66-139 segmented nuclei on three or more distinct acquisition fields. Statistical significance was calculated using one-way analysis of variance with a Bonferroni post-test (*p < 0.05; ***p < 0.001).

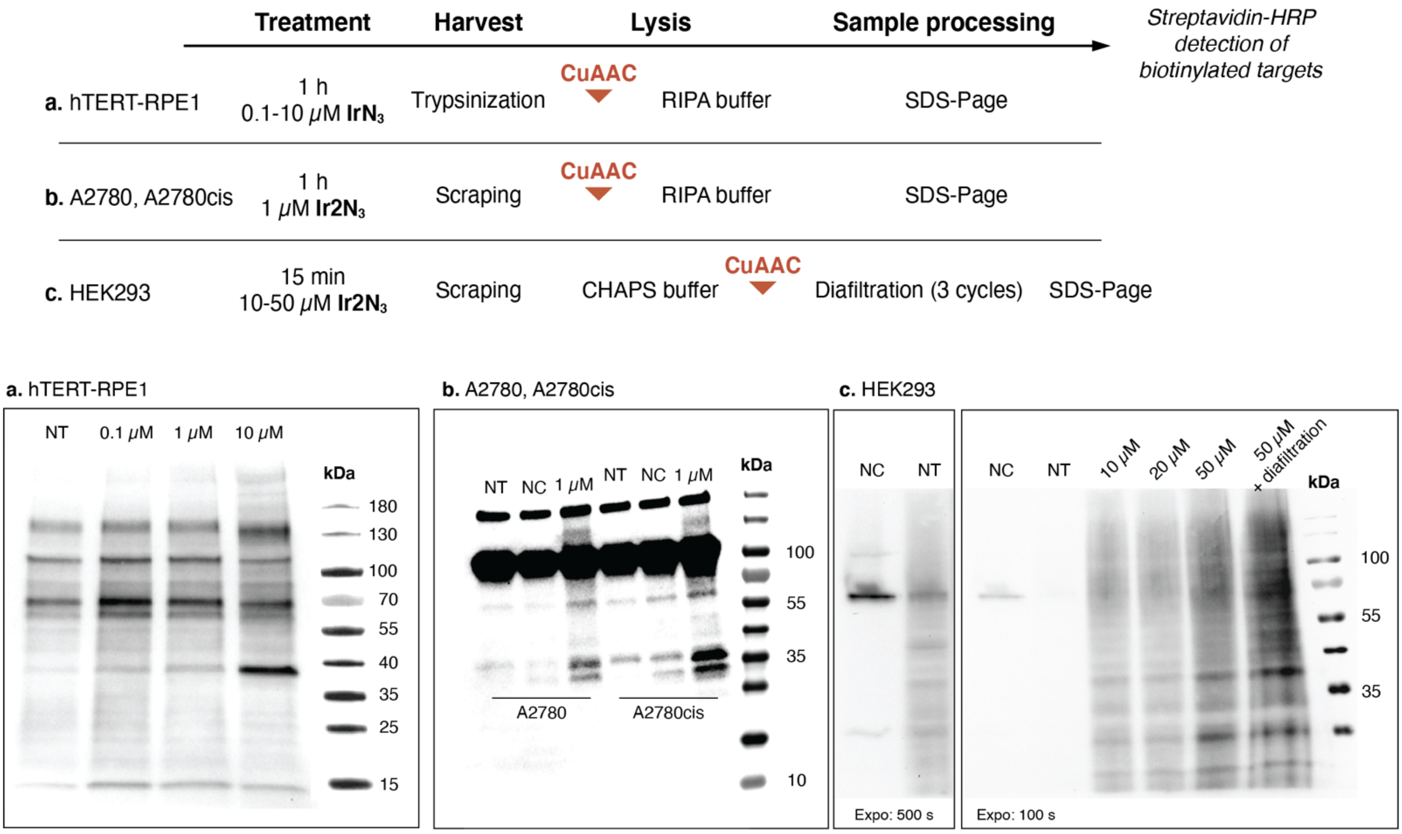

**Figure S6.** On-blot detection of **$Ir2N_3$** protein adducts using CuAAC with an ethynyl-substituted biotin and streptavidin peroxidase

**a.** hTERT-RPE1, **b.** A2780 and A2780cis, and **c.** HEK293 cells were treated with vehicle (NT: non treated) or the indicated concentrations of **$IrN_3$** or **$Ir2N_3$**, then harvested and submitted to CuAAC with a clickable ethynyl-biotin derivative before or after cell lysis. After protein separation by SDS-PAGE and blotting, biotinylated proteins were visualized with streptavidin-HRP. This procedure was found to give a low signal-to-noise ratio owing to the presence of endogenous biotinylated proteins when the click reaction was performed before lysis (**a** and **b**). These preliminary experiments allowed the detection of selective targets, with a more prominent band at *ca.* 39 kDa in all three cell lines. Sample diafiltration after CuAAC improved the detection of iridium protein adducts over endogenous biotinylated proteins (**c**). NC (non clicked): untreated cells with the click reaction omitted.

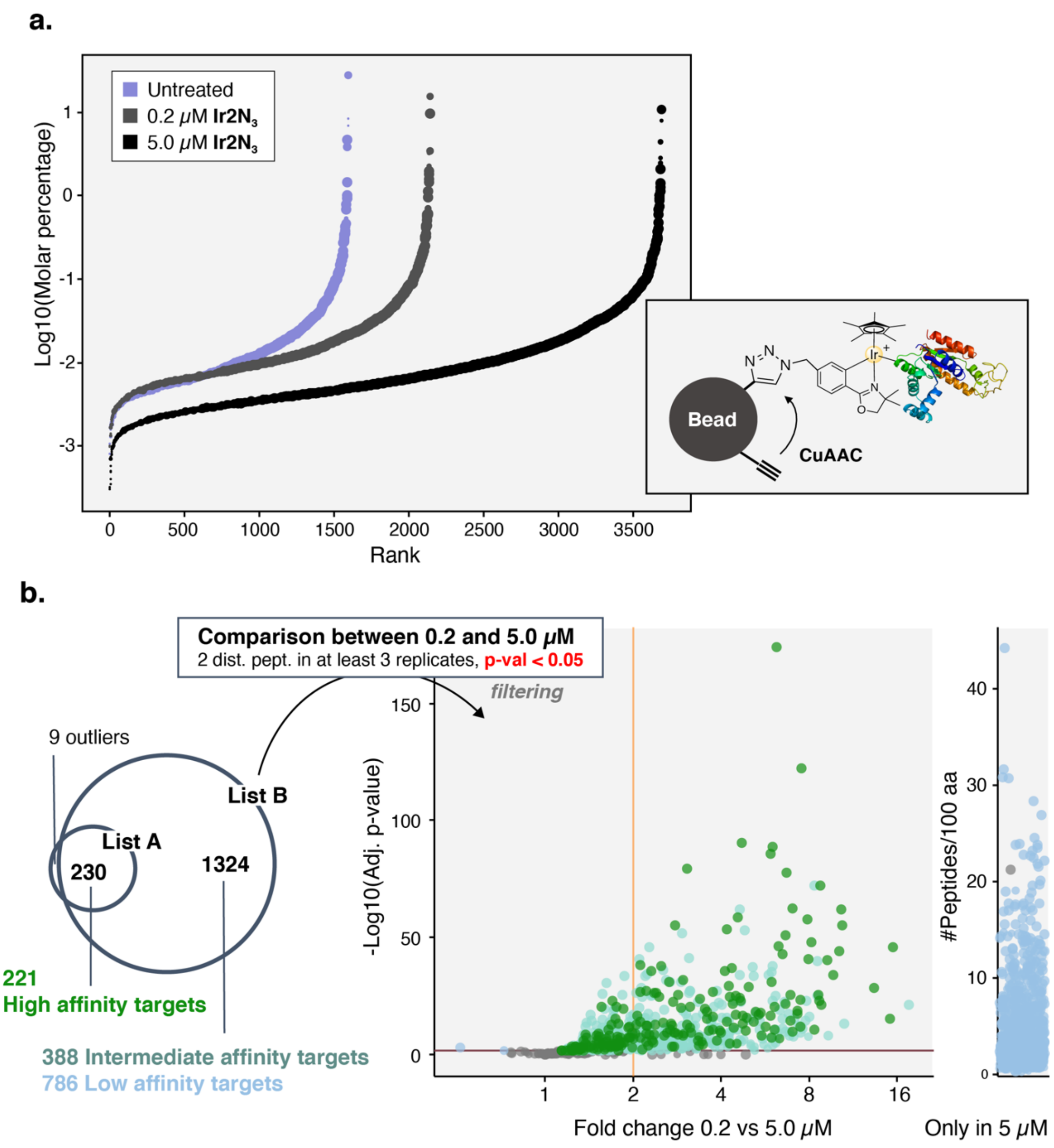

**Figure S7.** Absolute quantification and scheme of the chemoproteomic analysis of targets

**a.** Graphical representation of the results of absolute quantification performed over the three conditions. Scheme of the capture of **Ir2N3**-bound targets on ethynyl-beads by CuAAC.
**b.** Venn diagram of lists A (proteins enriched in the 0.2 μM treated *versus* non treated sample) and B (proteins enriched in the 5 μM treated *versus* non treated sample). Volcano plot showing statistical significance (p-value) versus magnitude of change (fold change between the two concentrations of **Ir2N3**) for the 1,554 targets from list B. High affinity targets intersects with List A (green), low affinity targets are only found in the 5.0 μM sample (blue) and the remaining proteins constitute the intermediate affinity targets list.

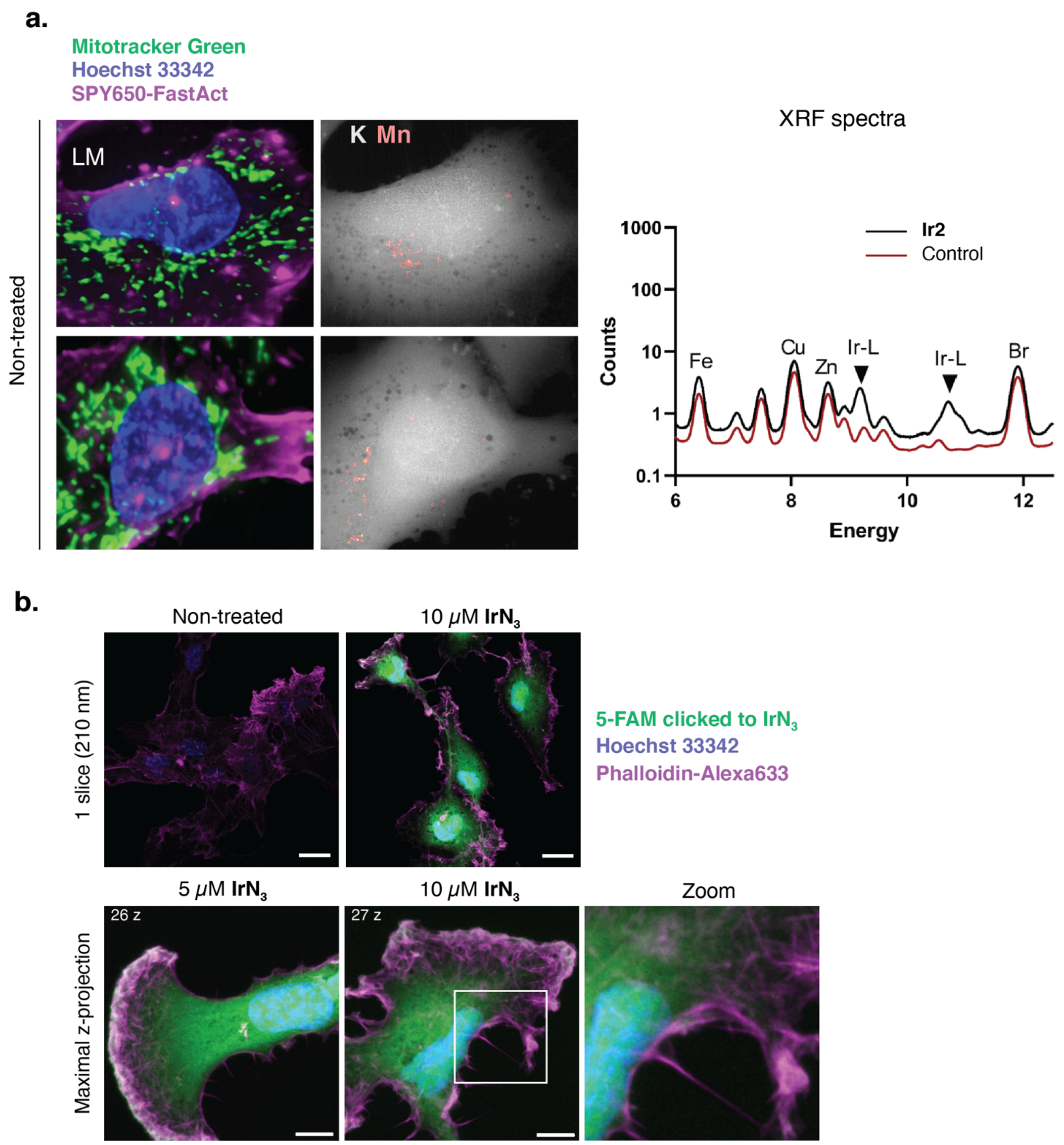

**Figure S8.** Bioorthogonal labelling of **$IrN_3$** by ethynyl-FAM and correlative LM-XRF imaging of **Ir2** in hTERT-RPE1 cells (in support to **Figure 4**)

**a.** Correlative analysis of cells treated with 5 µM **Ir2** for 15 min. Light cryomicroscopy (LM) of the actin cytoskeleton (SPY650-Fast Act, magenta), mitochondria (Mitotracker Green) and the nucleus (Hoechst 33342, blue). Cryo-X-Ray Fluorescence quantitative imaging (XRF) of elemental Mn (accumulated in Golgi structures) and K (present in the whole cell volume) acquired at 50 nm/px/50 ms resolution. Scale bars: 10 and 1 µm respectively. Right: Sum X-ray fluorescence spectra normalized to the number of pixels in XRF cell maps of control and **Ir2**-treated cells showing the Ir-L X-ray emission lines absent in the control cases.

**b.** Confocal images of cells exposed to **$IrN_3$** (less toxic than **$Ir2N_3$**) for 1 h, fixed and clicked *in situ* with ethynyl-FAM (green). Cells were stained for actin cytoskeleton (phalloidin, magenta) and DNA (Hoechst 33342, blue). Colocalization of **$IrN_3$** and microfilaments appear white on the merged images. Scale bars: 20 and 5 µm, zoom on a selected region of 225 µm$^2$.

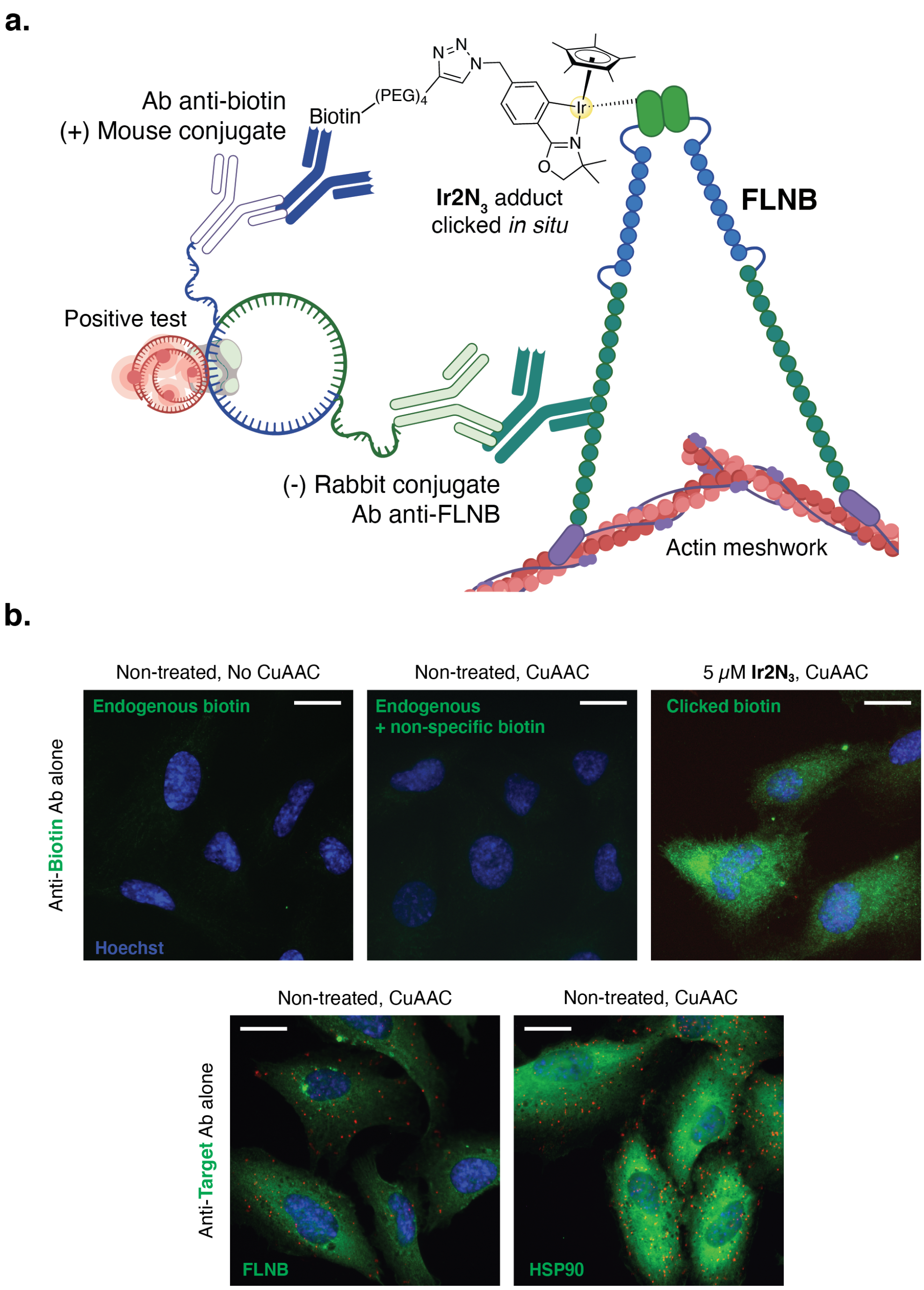

**Figure S9.** Principle of PLA and negative controls (in support to **Figure 5a**)

**a.** Scheme of the proximity ligation assay (PLA) carried out between a biotin-clicked to the Iridium complex and FLNB, probing interaction of the considered moieties at a distance inferior to 40 nm (created with BioRender.com).
**b.** Negative controls of the proximity ligation assay (red) performed on hTERT-RPE1 cells with only the anti-biotin or anti-target antibody, post-PLA detection by immunofluorescence of the considered target (green) and staining of nuclei (Hoechst 33342, blue). Both endogenous biotin and free ethynyl-biotin from CuAAC give little to no immunofluorescence signal and no PLA signal. Anti-FLNB and anti-HSP90 antibodies used alone give a background PLA signal used for normalization (**Fig 5b**). Scale bars 20 μm.

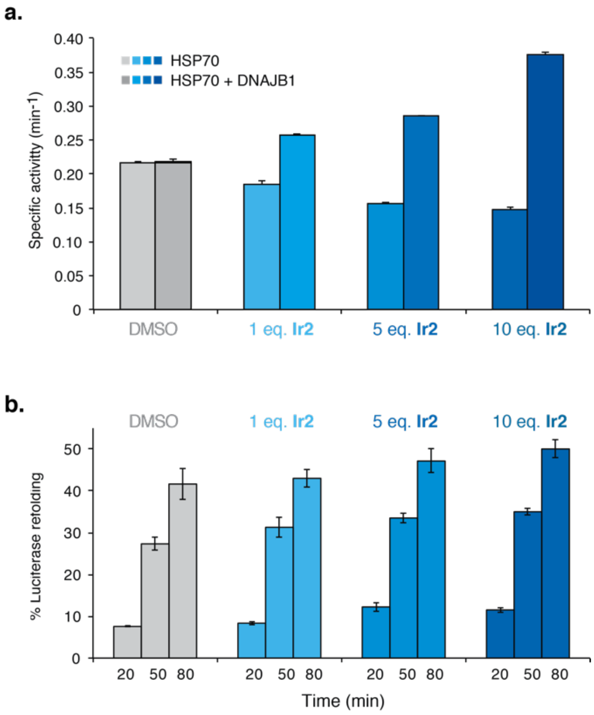

**Figure S10. Ir2** labeled HSP70 functional assays

**a.** HSP70 ATPase activity in absence (light shades) and presence (dark shades) of DNAJB1 cochaperone after incubation of the chaperone with DMSO vehicle (grey), 1 molar eq. **Ir2** (light blue), 5 molar eq. **Ir2** (medium blue) and 10 molar eq. **Ir2** (dark blue), given as mean of three replicates ± SD.
**b.** Luciferase refolding activity, reported as the percentage of luciferase renaturation, of HSP70 incubated with DMSO vehicle (grey), 1 molar eq. **Ir2** (light blue), 5 molar eq. **Ir2** (medium blue) and 10 molar eq. **Ir2** (dark blue), given as mean of three replicates ± SD.

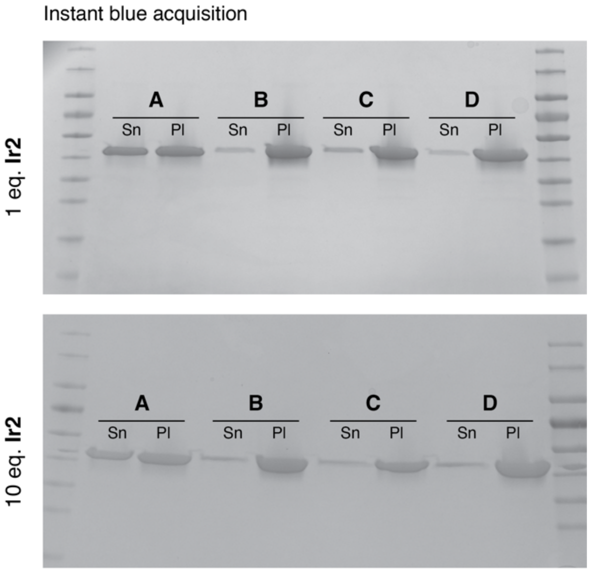

**Figure S11.** *In vitro* polymerization of actin monomer in presence of **Ir2**

**A**: No ATP added, negative control. **B**: Polymerisation procedure with 15 min pre-incubation of the actin monomer with 1 or 10 µL DMSO. **C** Polymerisation procedure with 15 min of pre-incubation with 10 or 100 µM **Ir2N**$_3$. **D**: Polymerisation procedure with 15 min post-incubation in the same conditions. Supernatants (Sn) were separated from pellets (Pl) and samples were migrated on polyacrylamide gels by SDS-Page. The gels were stained using Instant-Blue to reveal the protein content.

**Video S1.** Cell motility over 9 h

Differential phase contrast videomicroscopy of hTERT-RPE1 cells treated with **Ir2**, **IrN$_3$** or **Ir2N$_3$** at concentrations of 5 and 10 µM or left untreated in medium containing 5% FBS. The cell behaviour was monitored in real time using an Olympus IX83 inverted microscope with 20x objective and time-lapse movies (1 frame per 15 min) were processed using Fiji.

**Video S2.** Actin cytoskeleton dynamics over 3 h
Fluorescence time-lapse videomicroscopy of live hTERT-RPE1 cells stained for actin. Prior to acquisition, cells were incubated for 90 min with SPY650-FastAct (Spirochrome®) and left untreated (control cells, left) or treated with 5 µM **Ir2** (right) just before recording. The signal of the fluorescent probe (far red) was acquired during 180 min using an Olympus IX83 inverted videomicroscope with a 63x objective and time-lapse movies (1 frame per min) were processed using Fiji.

**Table S1.** Crystal data and structure refinement for [**Ir2**-DMSO], **IrN$_3$** and **Ir2N$_3$**

| | **Ir2**-DMSO | **IrN$_3$** | **Ir2N$_3$** |
|---|---|---|---|
| ***CCDC deposit number*** | *2205966* | *2205967* | *2205968* |
| **Empirical formula[a]** | $C_{23}H_{33}F_6IrNO_2PS$ | $C_{20}H_{24}ClIrN_4O$ | $C_{23}H_{30}Cl_3IrN_4O$ |
| **Moiety Formula** | $C_{23}H_{33}IrNO_2S^+$, $PF_6^-$ | $C_{20}H_{24}ClIrN_4O$ | $C_{22}H_{28}ClIrN_4O$, $CH_2Cl_2$ |
| **Formula weight (g/mol)** | 724.73 | 564.08 | 677.06 |
| **Temperature (K)** | 200 | 200 | 200 |
| **Crystal system** | Monoclinic | Monoclinic | Monoclinic |
| **Space group** | $P2_1$ | $P2_1/n$ | $P2_1/n$ |
| **a (Å)** | 9.1589(15) | 9.2821(3) | 9.8126(6) |
| **b (Å)** | 15.727(3) | 11.3807(4) | 21.2326(13) |
| **c (Å)** | 9.1986(15) | 19.0861(6) | 12.2881(7) |
| **α (°)** | 90 | 90 | 90 |
| **β (°)** | 104.516(3) | 101.900(2) | 99.938(3) |
| **γ (°)** | 90 | 90 | 90 |
| **Volume (Å$^3$)** | 1282.7(4) | 1972.86(11) | 2521.8(3) |
| **Z** | 2 | 4 | 4 |
| **$\rho_{calc}$ (g/cm$^3$)** | 1.876 | 1.899 | 1.783 |
| **Absorption coefficient μ (mm$^{-1}$)** | 5.414 (Mo Kα) | 6.921 (Mo Kα) | 5.635 (Mo Kα) |
| **F(000)** | 712 | 1096.0 | 1328.0 |
| **Crystal size (mm²)** | 0.27 × 0.15 × 0.11 | 0.22 × 0.22 × 0.06 | 0.30 × 0.20 × 0.20 |
| **Wavelength λ (Å)** | 0.71073 | 0.71073 | 0.71073 |
| **2Θ range (°)** | 4.574 - 61.152 | 4.192 - 61.138 | 3.836 - 61.246 |
| **Miller indexes ranges** | -13 ≤ h ≤ 13,<br>-22 ≤ k ≤ 22,<br>-13 ≤ l ≤ 13 | -12 ≤ h ≤ 13,<br>-16 ≤ k ≤ 16,<br>-27 ≤ l ≤ 22 | -14 ≤ h ≤ 14,<br>-29 ≤ k ≤ 30,<br>-17 ≤ l ≤ 17 |
| **Measured reflections** | 27734 | 37492 | 41576 |
| **Unique reflections** | 7848 | 6042 | 7732 |
| **$R_{int}$ / $R_{sigma}$** | 0.0287 / 0.0290 | 0.0477 / 0.0316 | 0.0312 / 0.0226 |
| **Reflections [I ≥ 2σ(I)]** | 7427 | 4988 | 6669 |
| **Restraints** | 1 | 0 | 170 |
| **Parameters** | 322 | 249 | 361 |
| **Goodness-of-fit F²** | 1.012 | 1.019 | 1.176 |
| **Final R indexes[b c] [all data]** | R1 = 0.0202,<br>wR2 = 0.0373 | R1 = 0.0318,<br>wR2 = 0.0469 | R1 = 0.0441,<br>wR2 = 0.0823 |
| **Final R indexes[b c] [I ≥ 2σ(I)]** | R1 = 0.0181,<br>wR2 = 0.0368 | R1 = 0.0218,<br>wR2 = 0.0441 | R1 = 0.0359,<br>wR2 = 0.0793 |
| **Largest diff. peak/hole (e/Å$^3$)** | 0.56/-0.57 | 1.16/-0.75 | 2.38/-2.46 |
| **Flack parameter** | -0.025(3) | Not applicable | Not applicable |

[a]Including solvent molecules (if presence)
[b] $R1 = \sum||F_o| - |F_c|| / \sum|F_o|$ [c] $wR2 = \sqrt{\sum(w(F_o^2 - F_c^2))/\sum(w(F_o^2)^2)}$

**Table S2.** Selected distances (Å ±98% CI) and angles (° ±98% C) for [**Ir2**-DMSO], **IrN$_3$** and **Ir2N$_3$**

| | **Ir2-DMSO** | **IrN$_3$** | **Ir2N$_3$** |
|---|---|---|---|
| **Ir—C** | 2.068 (0.012) | 2.045 (0.006) | 2.050 (0.012) |
| **Ir—N** | 2.106 (0.009) | 2.089 (0.006) | 2.112 (0.012) |
| **Ir—Z** | 2.302 (0.003) | 2.394 (0.002) | 2.409 (0.003) |
| **Ir-centroid#** | 1.872 (0.001) | 1.821 (0.001) | 1.827 (0.001) |
| **C-Ir-N** | 77.82 (0.42) | 77.17 (0.27) | 77.79 (0.38) |
| **N-Ir-Z** | 84.64 (0.33) | 85.74 (0.21) | 85.15 (0.33) |
| **Z-Ir-C** | 82.57 (0.30) | 85.07 (0.21) | 85.28 (0.38) |
| **C-Ir-centroid#** | 127.36 (0.32) | 133.02 (0.21) | 131.56 (0.35) |
| **N-Ir-centroid#** | 131.33 (0.24) | 134.35 (0.19) | 136.06 (0.32) |
| **Z-Ir-centroid#** | 133.42 (0.08) | 123.78 (0.07) | 123.28 (0.10) |

Z : Cl for **IrN$_3$** and **Ir2N$_3$**, DMSO for **Ir2**-DMSO
**#** : the centroid of the 5-membered ring composing Cp*

**Table S3.** Observed adducts in MeOH with the lateral chain of model substrates (ESI-HRMS); relative intensity of peaks

| **Substrate** | **Calculated *m/z*** | **Observed *m/z*** | **Relative adduct intensity* (%)** |
|---|---|---|---|
| N-Acetyl-Histidine | 699.2522<br>721.2342 (M+Na) | 699.2518<br>721.2336 | 71 |
| N-Acetyl-Cysteine Methyl ester | 679.2182<br>701.2002 | 679.2178<br>701.1994 | 70 |
| Phenylbutylamine | 651.2926 | 651.2921 | 48 |
| Butyramide | 589.2406 | none | - |

* Relative intensities of total form of compounds are expressed in %. Please note that peak intensities depend on the ionizing efficiency and do not necessarily represent the amount of each species initially present in solution.

**Table S4.** Protein lists and GO analyses (spreadsheet)

### IV. Author Contributions

RR* Investigation, formal analysis, visualization, writing of original draft. AK, CB, Conceptualization, writing review and editing. MT FL JF SB PM TC CBe DL FD Data acquisition, formal analysis. AC CC MS Data acquisition. MS*, JST* Funding acquisition, data curation, project administration, manuscript editing, joint supervision. All authors analyzed data, discussed results. The manuscript was written and edited through contributions from all the authors.

### V. Annexes

1. Cryo-XRF (relevant elements) and photonic raw images acquired in a treated cell
2. Cryo-XRF (relevant elements) and photonic raw images acquired in untreated cells

3. $^{1}$H and $^{13}$C NMR (DEPT-135) Spectra of **Ir2-DMSO**
4. $^{1}$H and $^{13}$C NMR Spectra of **L06**
5. $^{1}$H and $^{13}$C NMR Spectra of **L06N$_3$**
6. $^{1}$H and $^{13}$C NMR Spectra of **IrN$_3$**
7. $^{1}$H and $^{13}$C NMR Spectra of **IrN$_3$-DMSO**
8. $^{1}$H and $^{13}$C NMR Spectra of **phenyl-©-IrN$_3$**
9. $^{1}$H and $^{13}$C NMR Spectra of **L08**
10. $^{1}$H and $^{13}$C NMR Spectra of **L08N$_3$**
11. $^{1}$H and $^{13}$C NMR (DEPT-135) Spectra of **Ir2N$_3$**

1. Cryo-XRF (relevant elements) and photonic raw images acquired in a treated cell

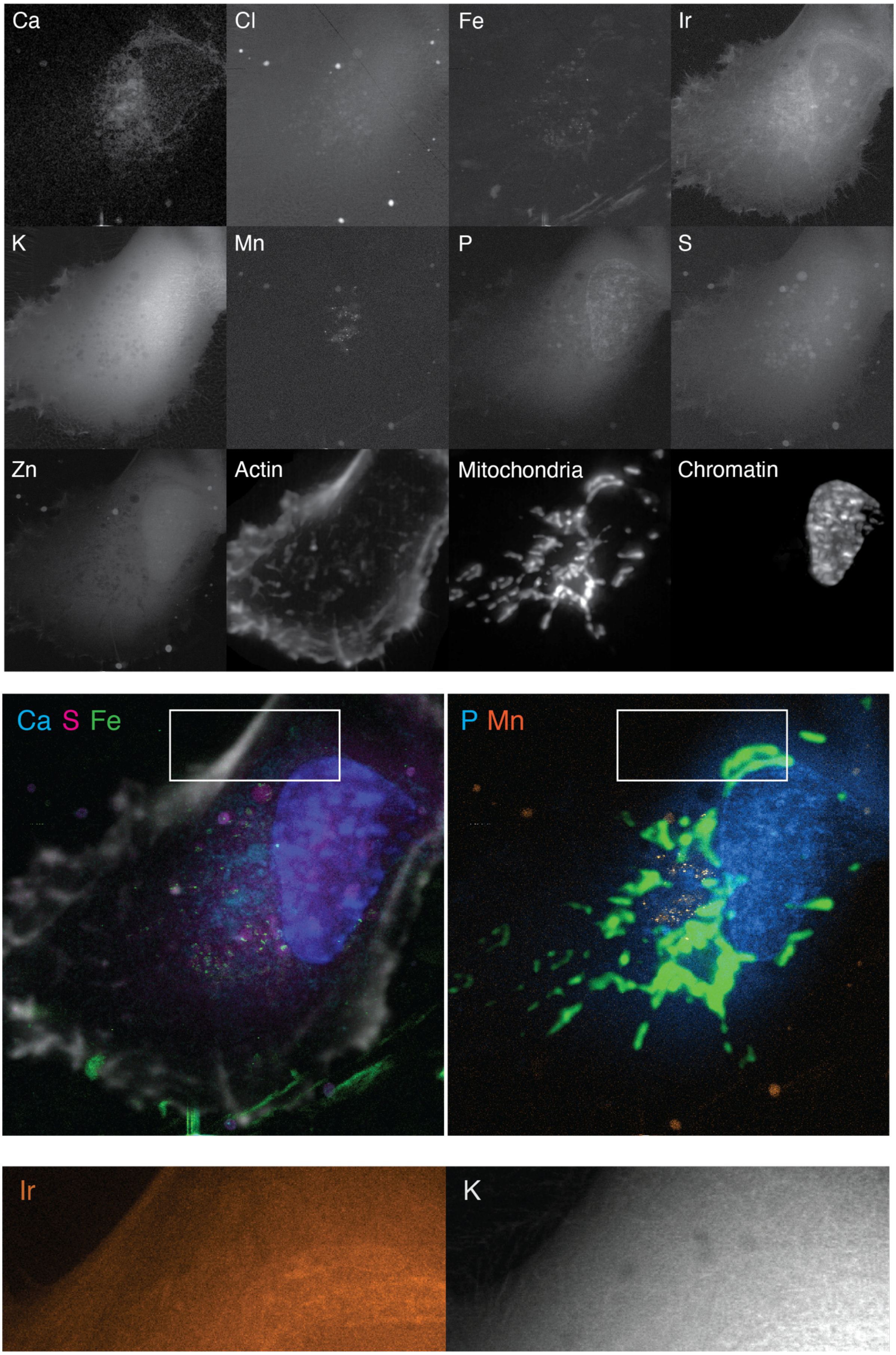

2. Cryo-XRF (relevant elements) and photonic raw images acquired in untreated cells

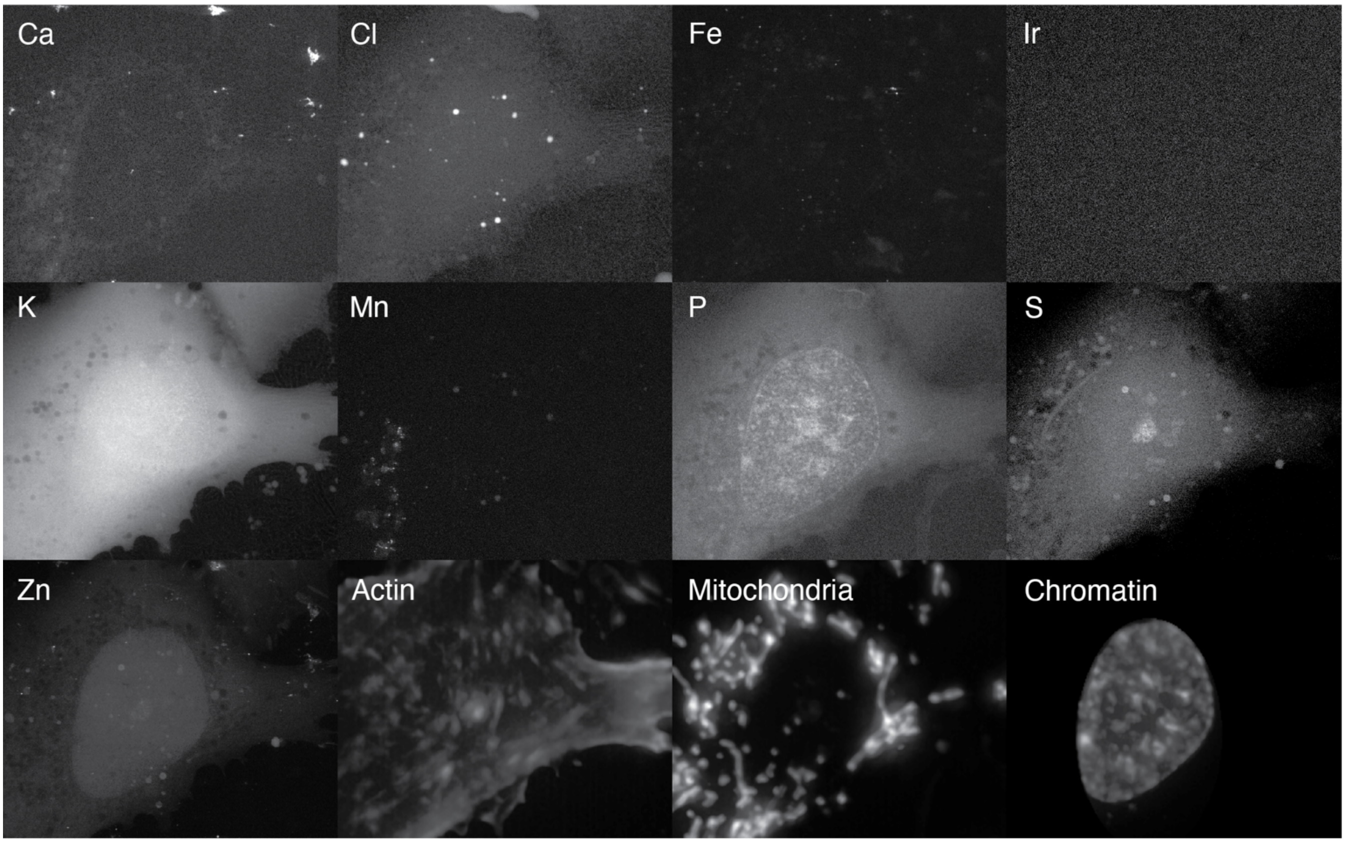

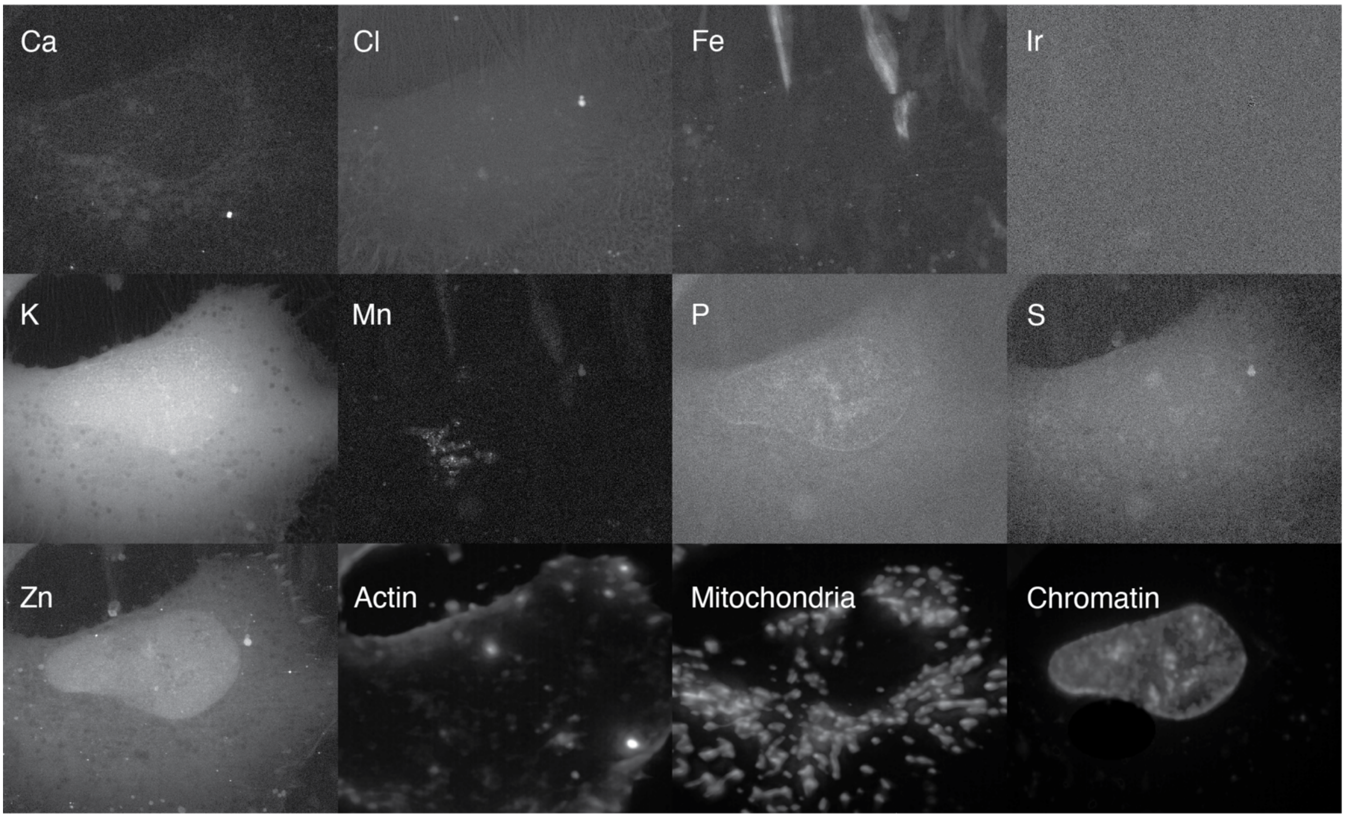

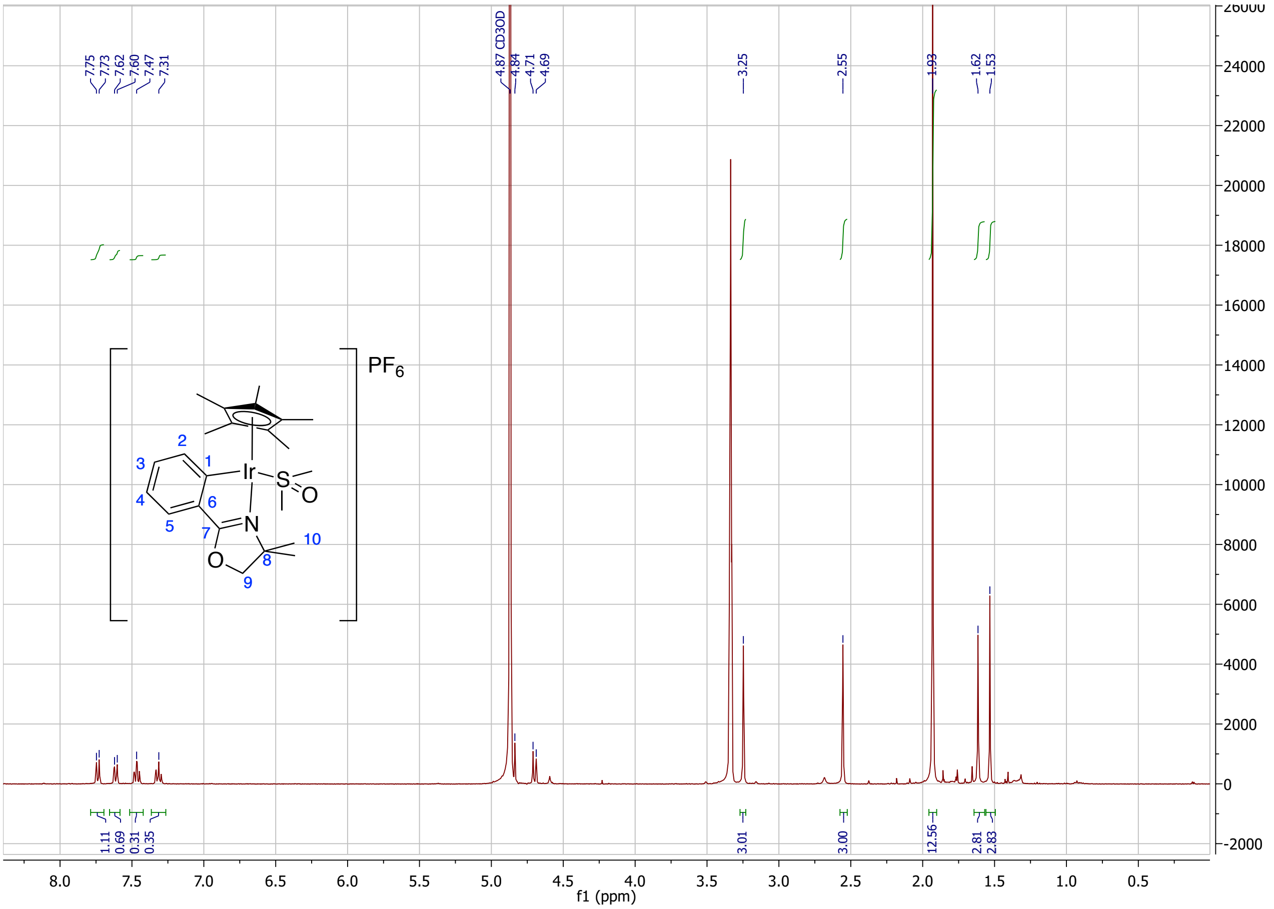
7.75
7.73
7.62
7.60
7.47
7.31
4.87 CD3OD
4.84
4.71
4.69
3.25
2.55
1.93
1.62
1.53
1.11
0.69
0.31
0.35
3.01
3.00
12.56
2.81
2.83
PF6
Ir
S
O
N
O
1
2
3
4
5
6
7
8
9
10
f1 (ppm)

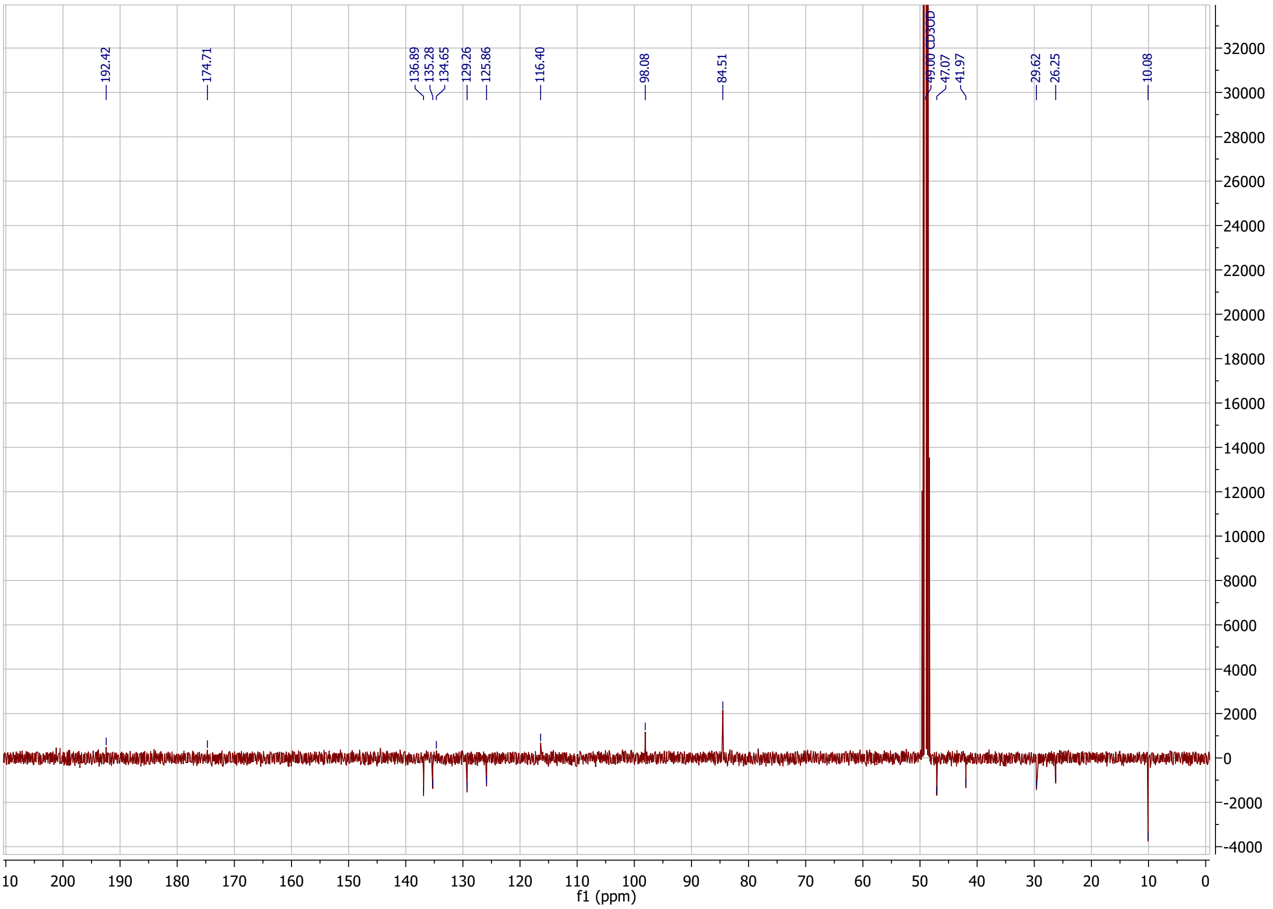
192.42
174.71
136.89
135.28
134.65
129.26
125.86
116.40
98.08
84.51
49.00 CD3OD
47.07
41.97
29.62
26.25
10.08
f1 (ppm)

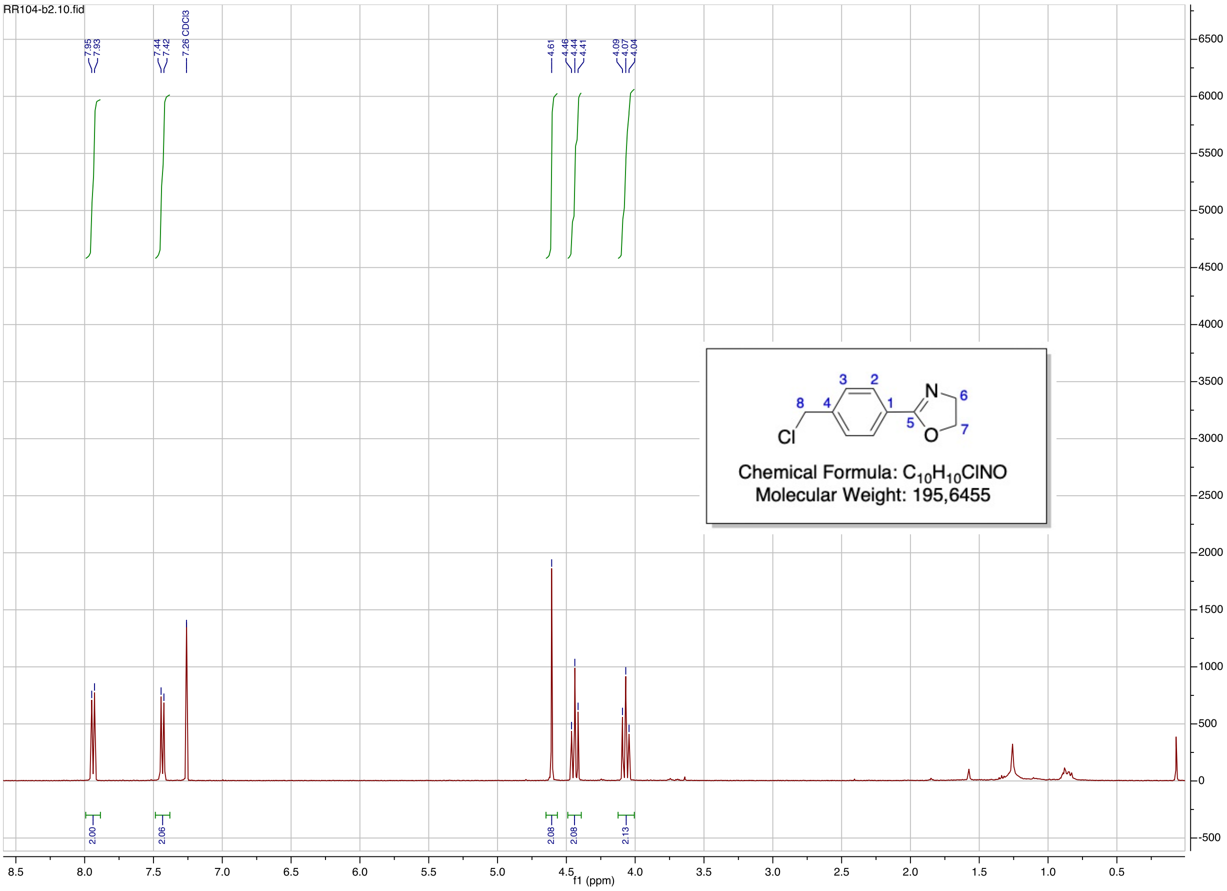
RR104-b2.10.fid
7.95
7.93
7.44
7.42
7.26 CDCl3
4.61
4.46
4.44
4.41
4.09
4.07
4.04
Cl
N
O
Chemical Formula: C10H10ClNO
Molecular Weight: 195,6455
2.00
2.06
2.08
2.08
2.13
f1 (ppm)

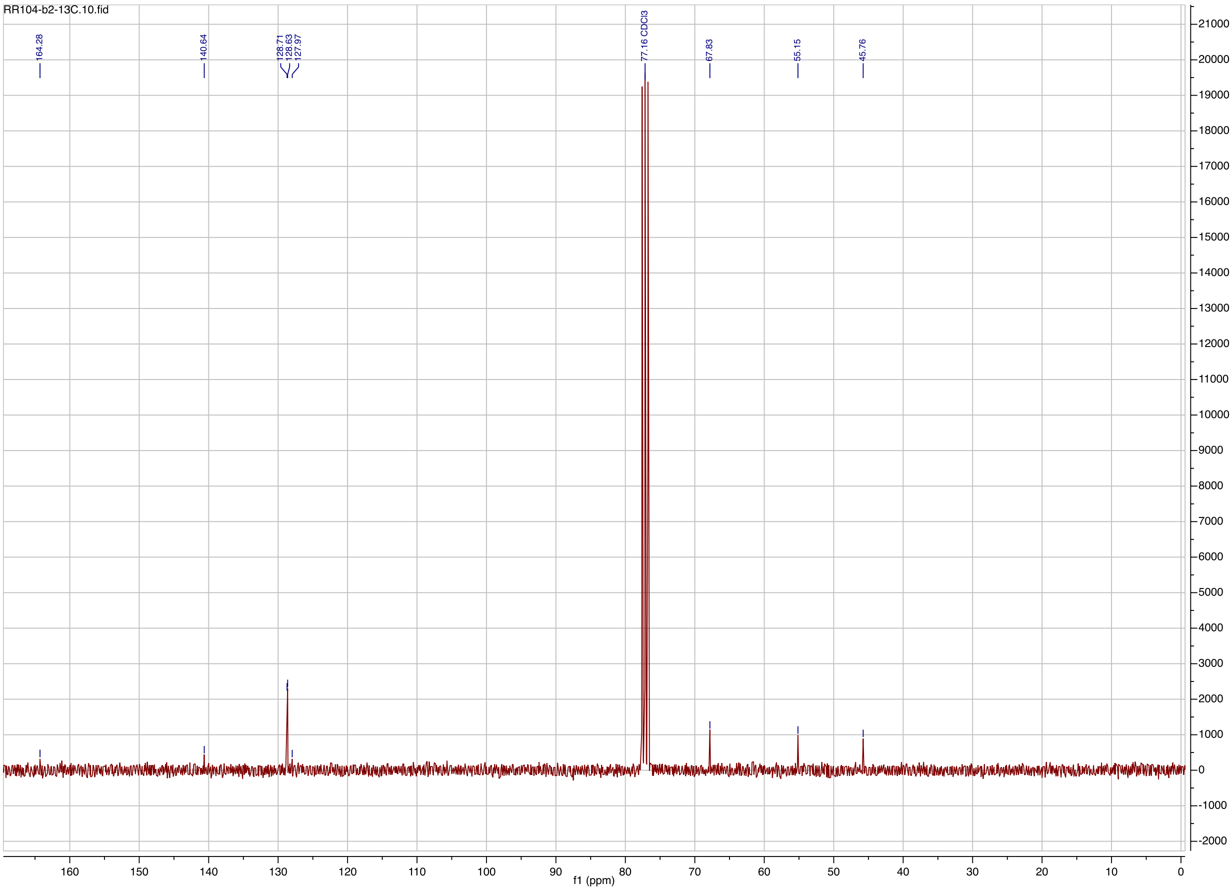
RR104-b2-13C.10.fid
164.28
140.64
128.71
128.63
127.97
77.16 CDCl3
67.83
55.15
45.76
f1 (ppm)

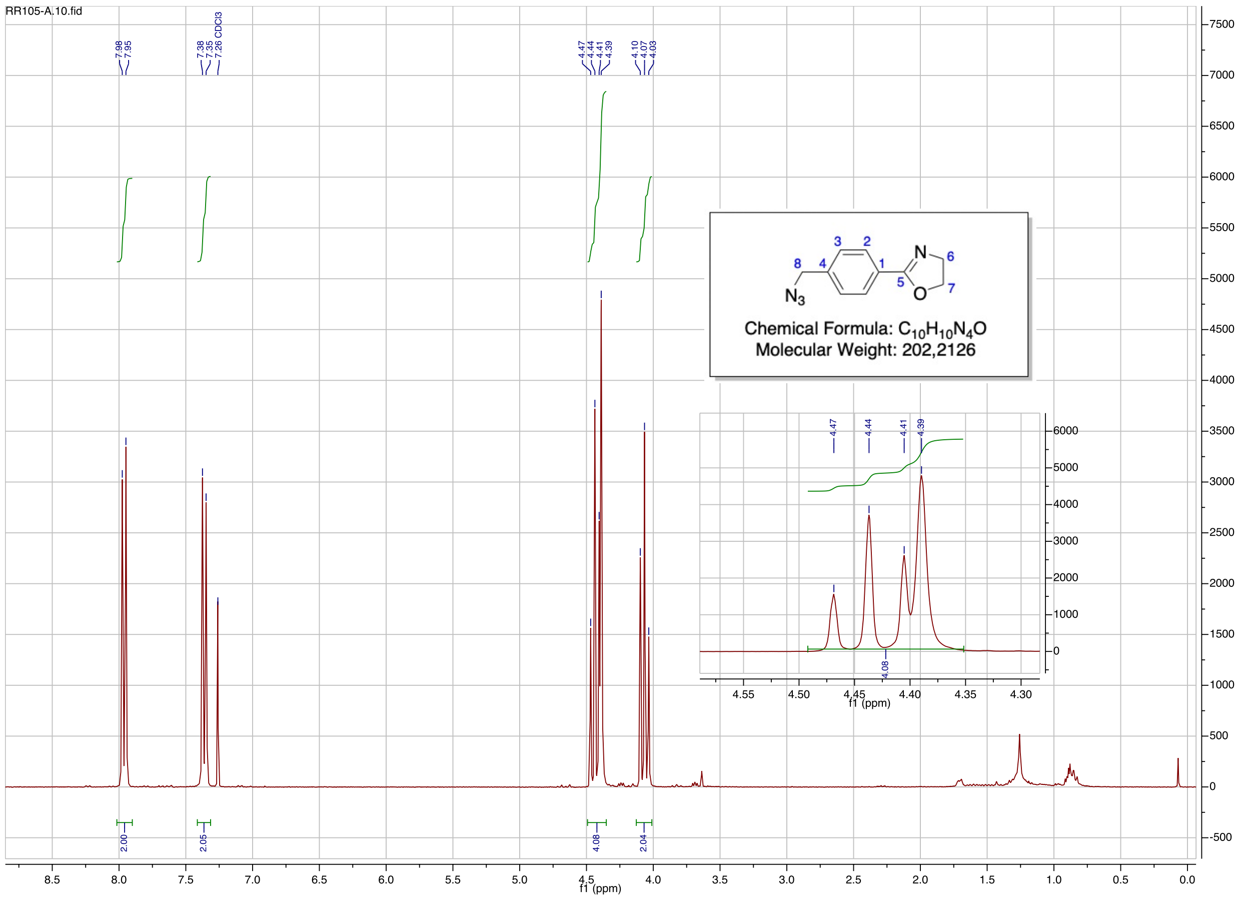
RR105-A.10.fid
7.98
7.95
7.38
7.35
7.26 CDCl3
4.47
4.44
4.41
4.39
4.10
4.07
4.03
Chemical Formula: C10H10N4O
Molecular Weight: 202,2126
2.00
2.05
4.08
2.04
f1 (ppm)

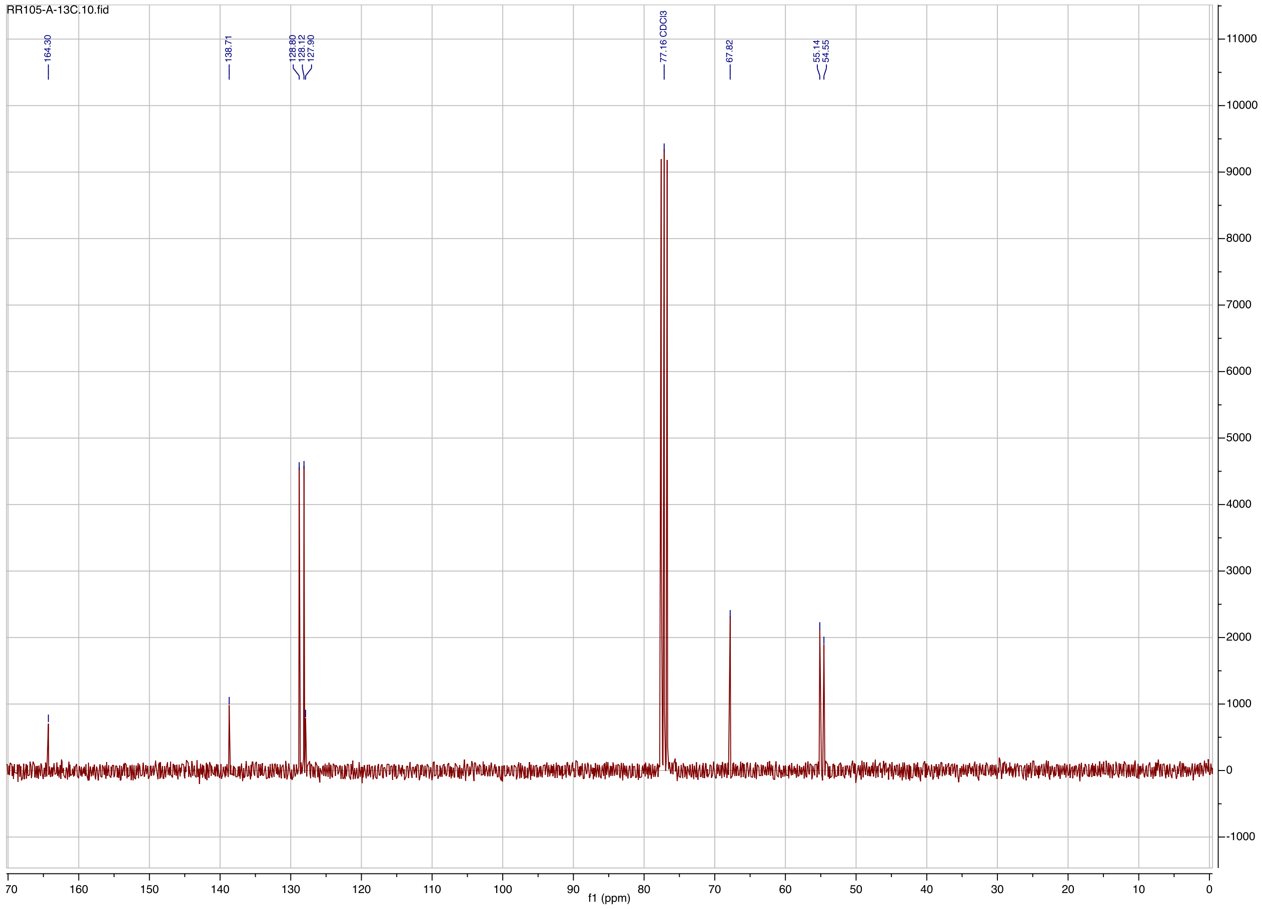
RR105-A-13C.10.fid
164.30
138.71
128.80
128.12
127.90
77.16 CDCl3
67.82
55.14
54.55
f1 (ppm)

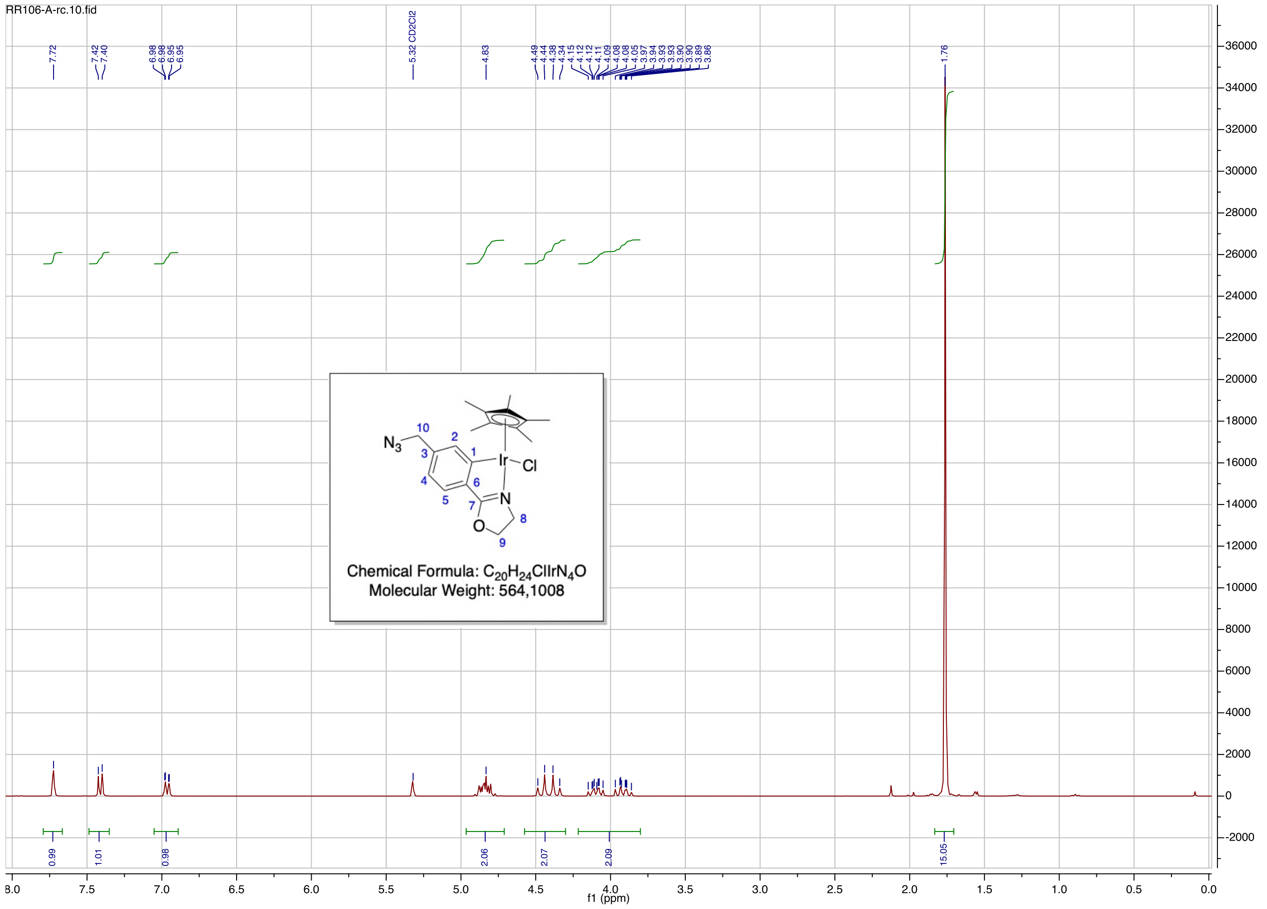
RR106-A-rc.10.fid
7.72
7.42
7.40
6.98
6.98
6.95
6.95
5.32 CD2Cl2
4.83
4.49
4.44
4.38
4.34
4.15
4.12
4.11
4.09
4.08
4.08
4.05
3.97
3.94
3.93
3.93
3.90
3.89
3.86
1.76
0.99
1.01
0.98
2.06
2.07
2.09
15.05
f1 (ppm)
N3
Ir
Cl
N
O
Chemical Formula: C20H24ClIrN4O
Molecular Weight: 564,1008

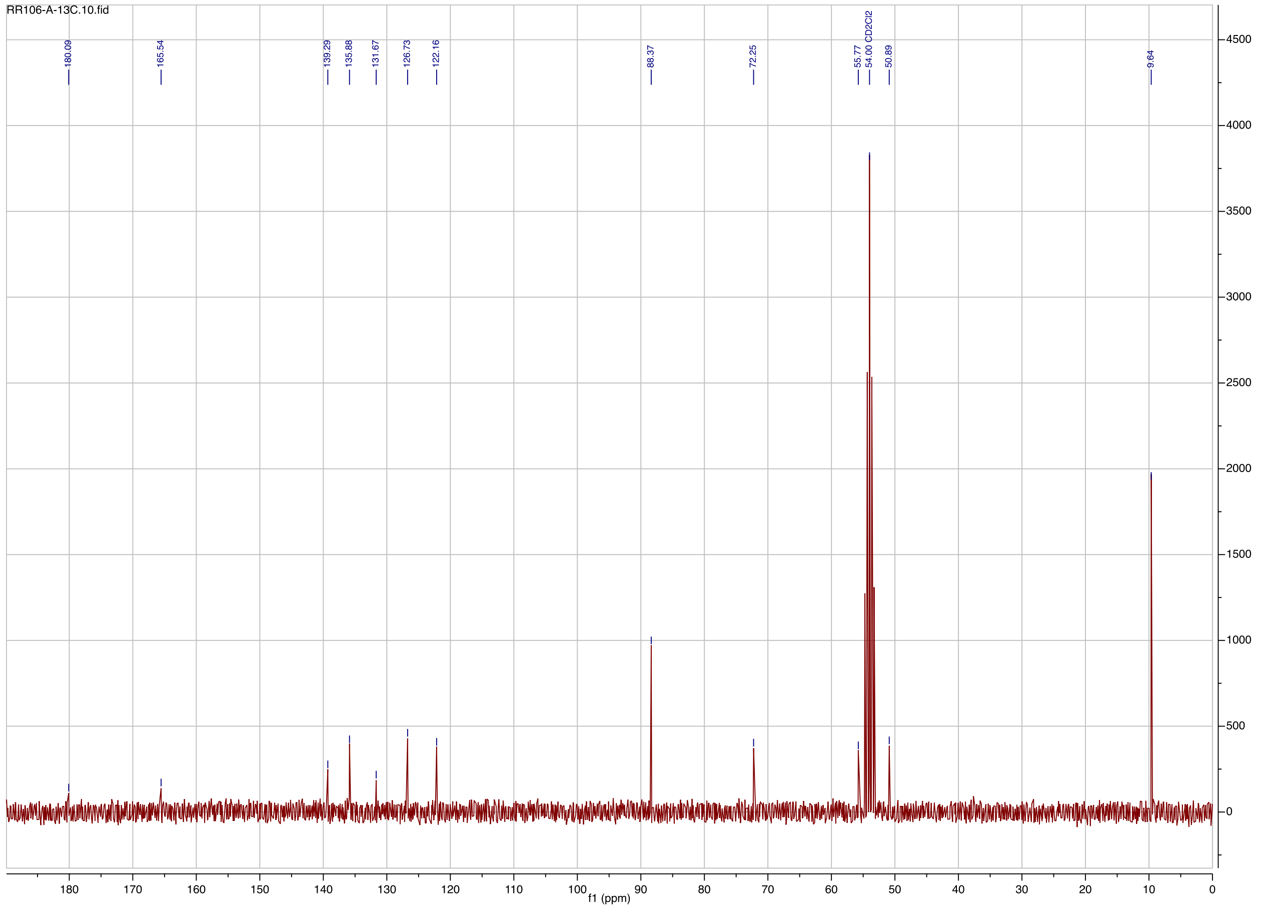
RR106-A-13C.10.fid
180.09
165.54
139.29
135.88
131.67
126.73
122.16
88.37
72.25
55.77
54.00 CD2Cl2
50.89
9.64
f1 (ppm)

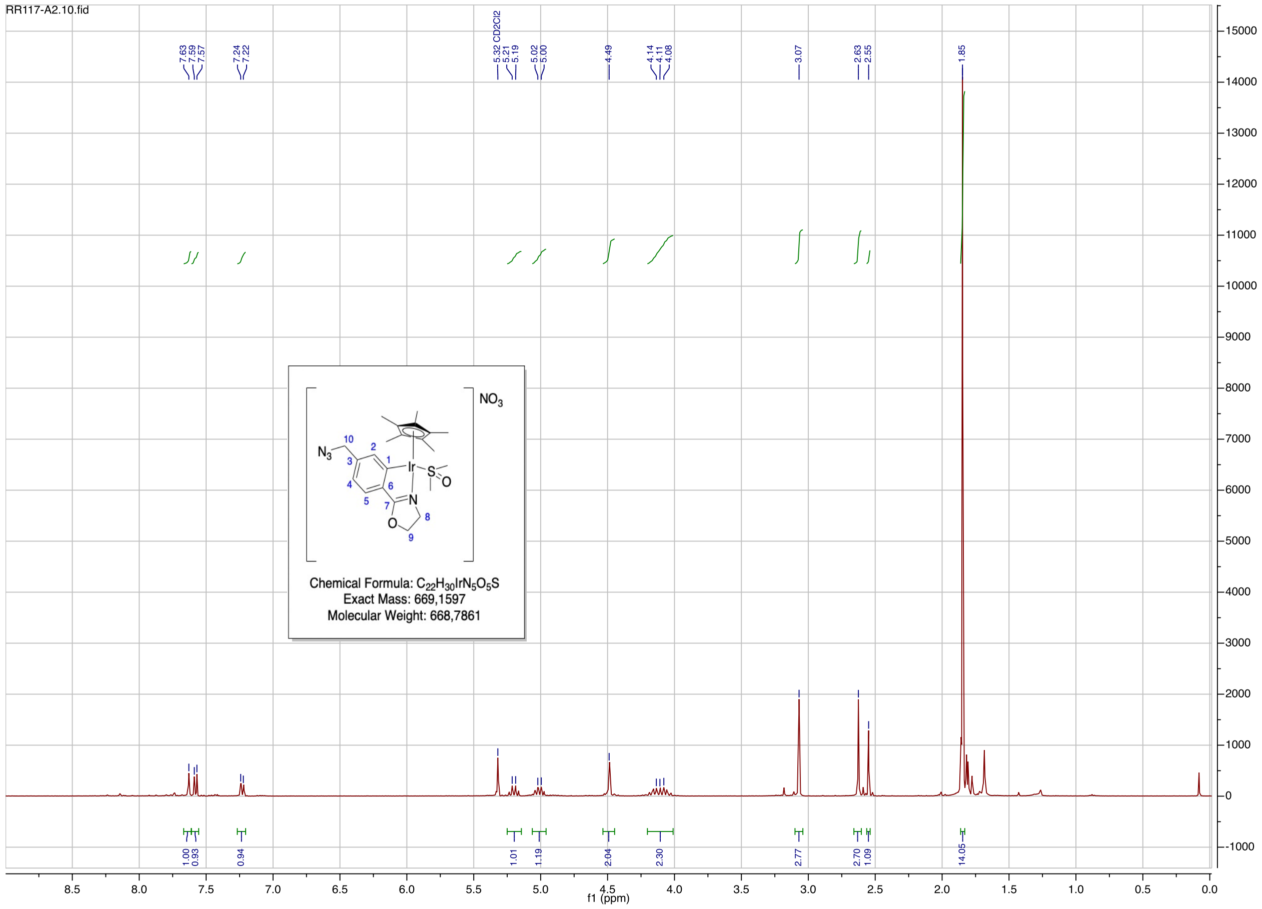
RR117-A2.10.fid
f1 (ppm)
NO3
Chemical Formula: C22H30IrN5O5S
Exact Mass: 669,1597
Molecular Weight: 668,7861

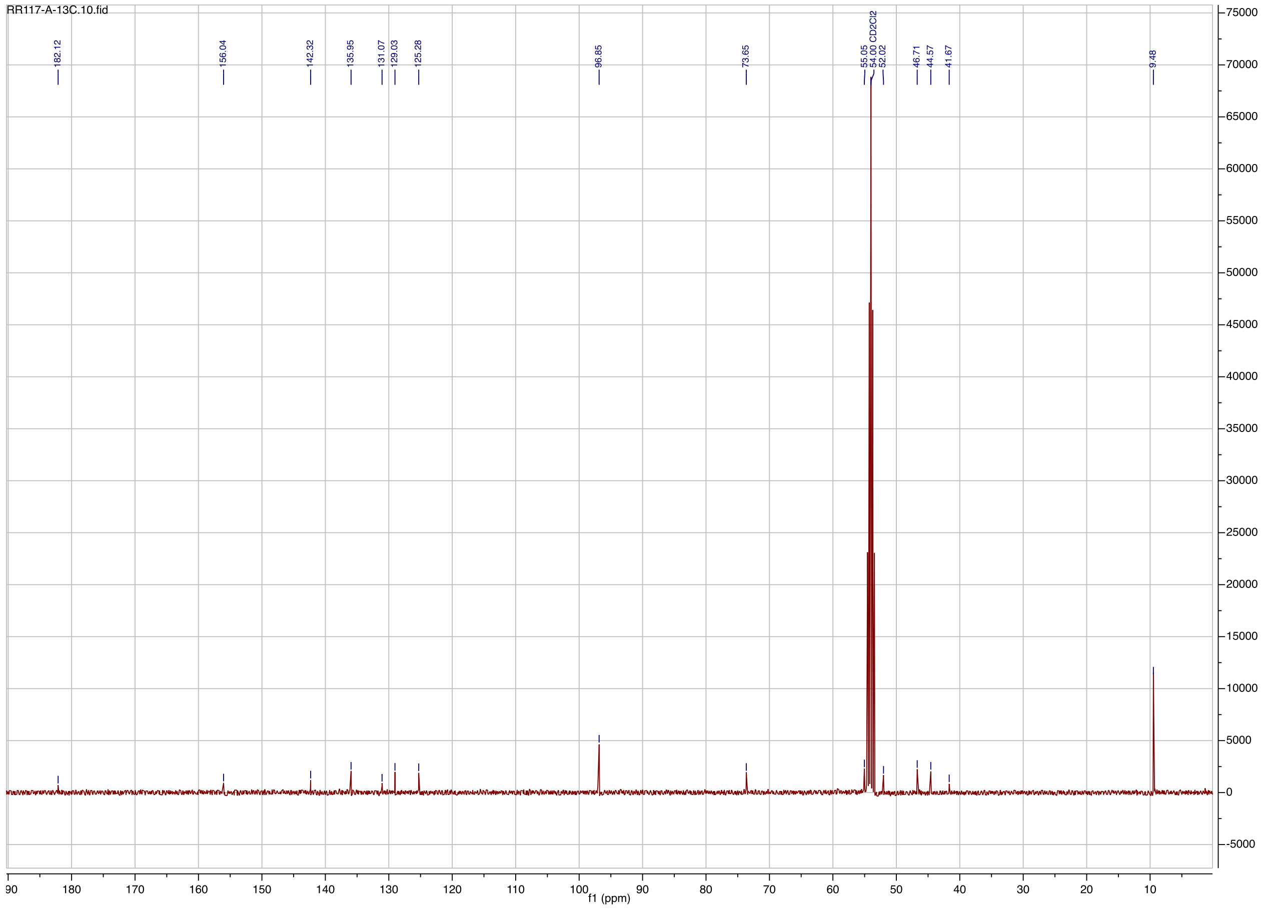
RR117-A-13C.10.fid
f1 (ppm)

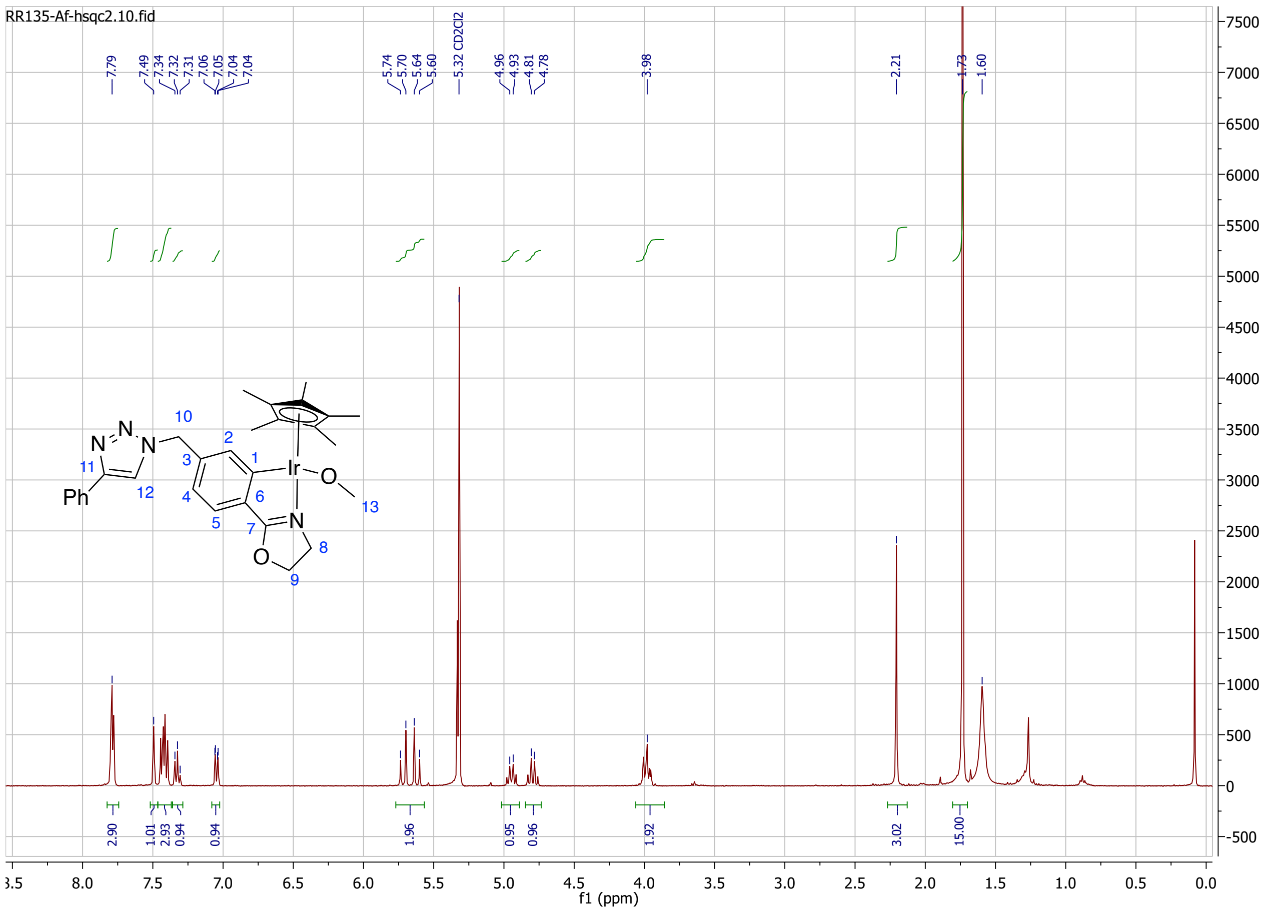
RR135-Af-hsqc2.10.fid
7.79
7.49
7.34
7.32
7.31
7.06
7.05
7.04
7.04
5.74
5.70
5.64
5.60
5.32 CD2Cl2
4.96
4.93
4.81
4.78
3.98
2.21
1.73
1.60
2.90
1.01
2.93
0.94
0.94
1.96
0.95
0.96
1.92
3.02
15.00
f1 (ppm)

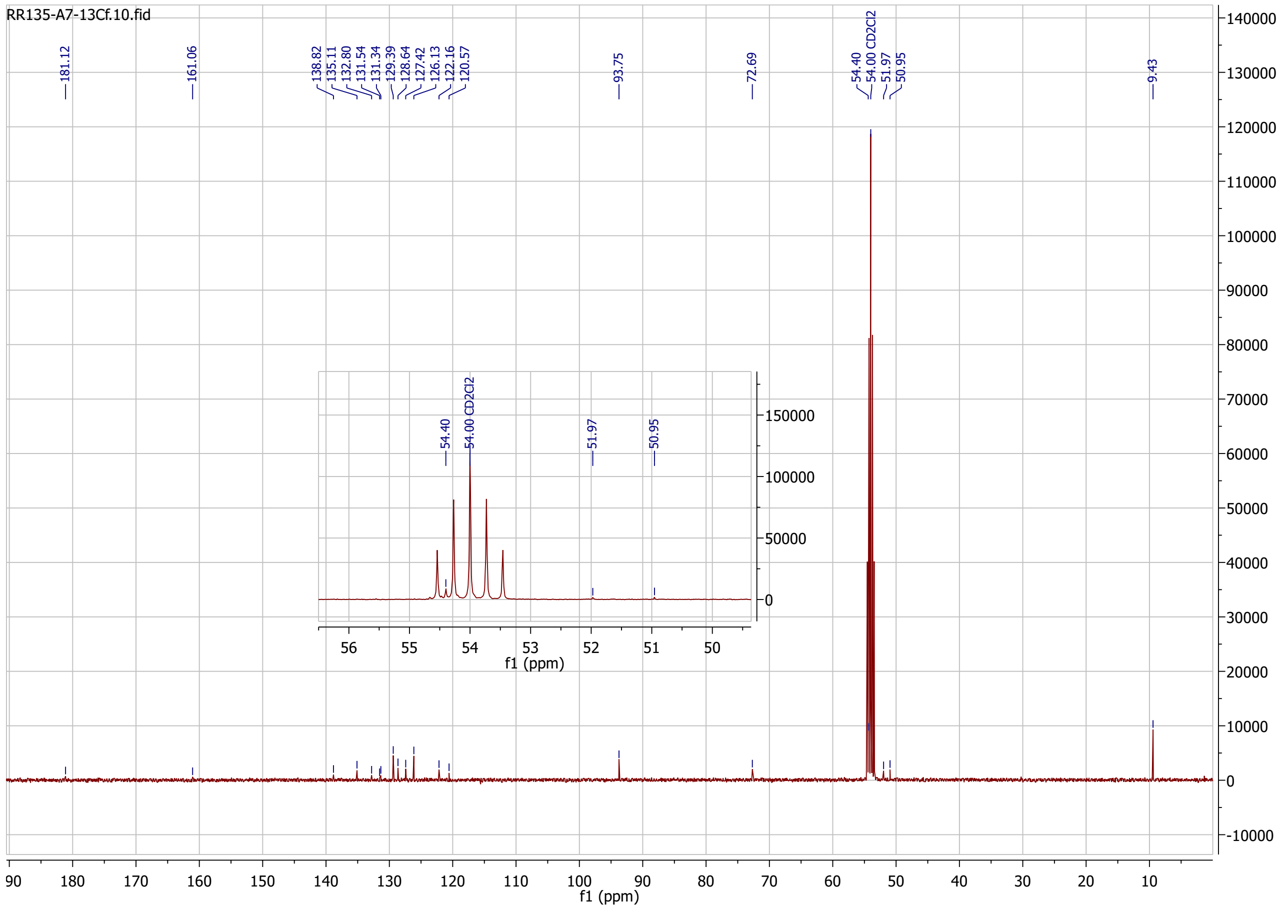
RR135-A7-13Cf.10.fid
181.12
161.06
138.82
135.11
132.80
131.54
131.34
129.39
128.64
127.42
126.13
122.16
120.57
93.75
72.69
54.40
54.00 CD2Cl2
51.97
50.95
9.43
f1 (ppm)

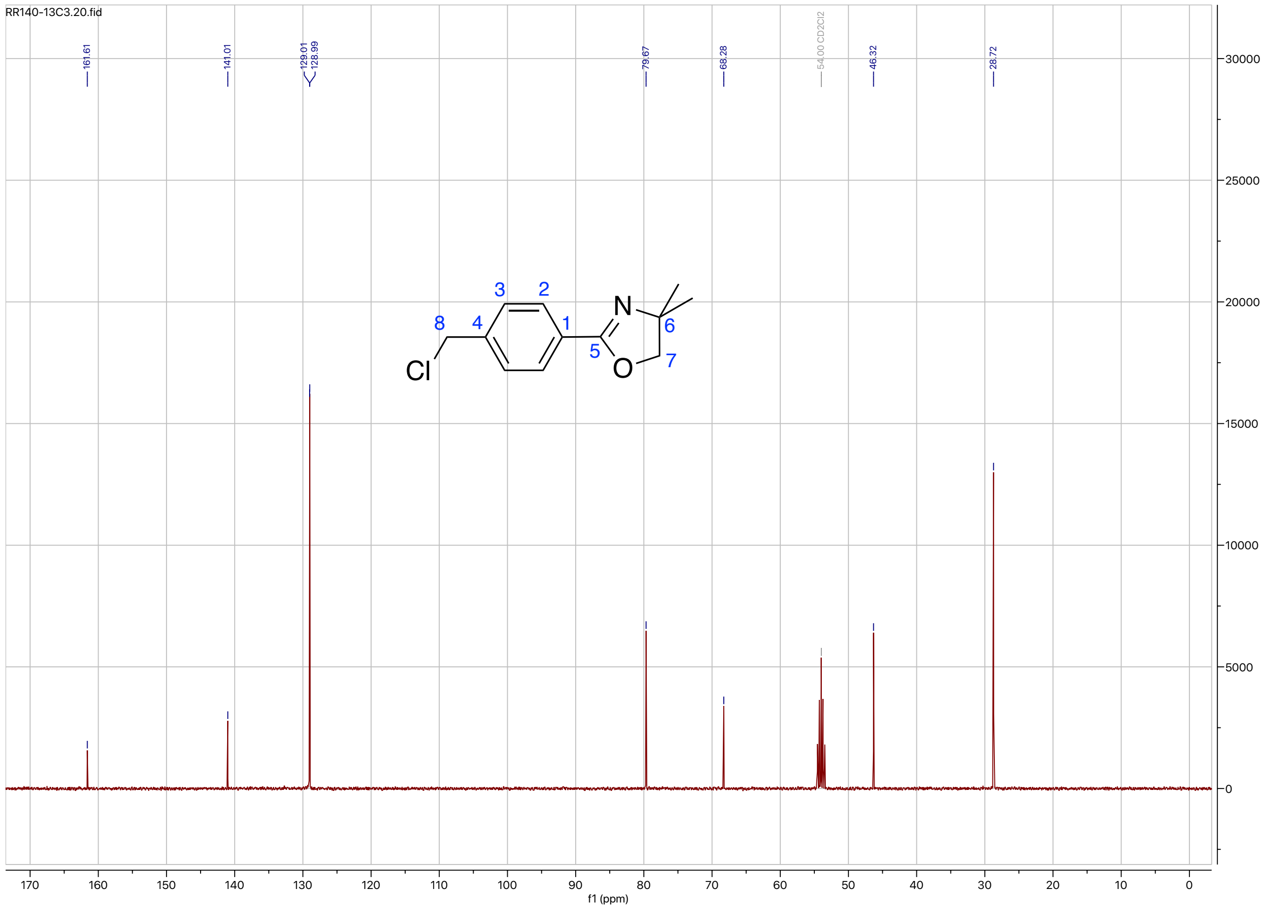
RR140-13C3.20.fid
161.61
141.01
129.01
128.99
79.67
68.28
54.00 CD2Cl2
46.32
28.72
Cl
N
O
f1 (ppm)

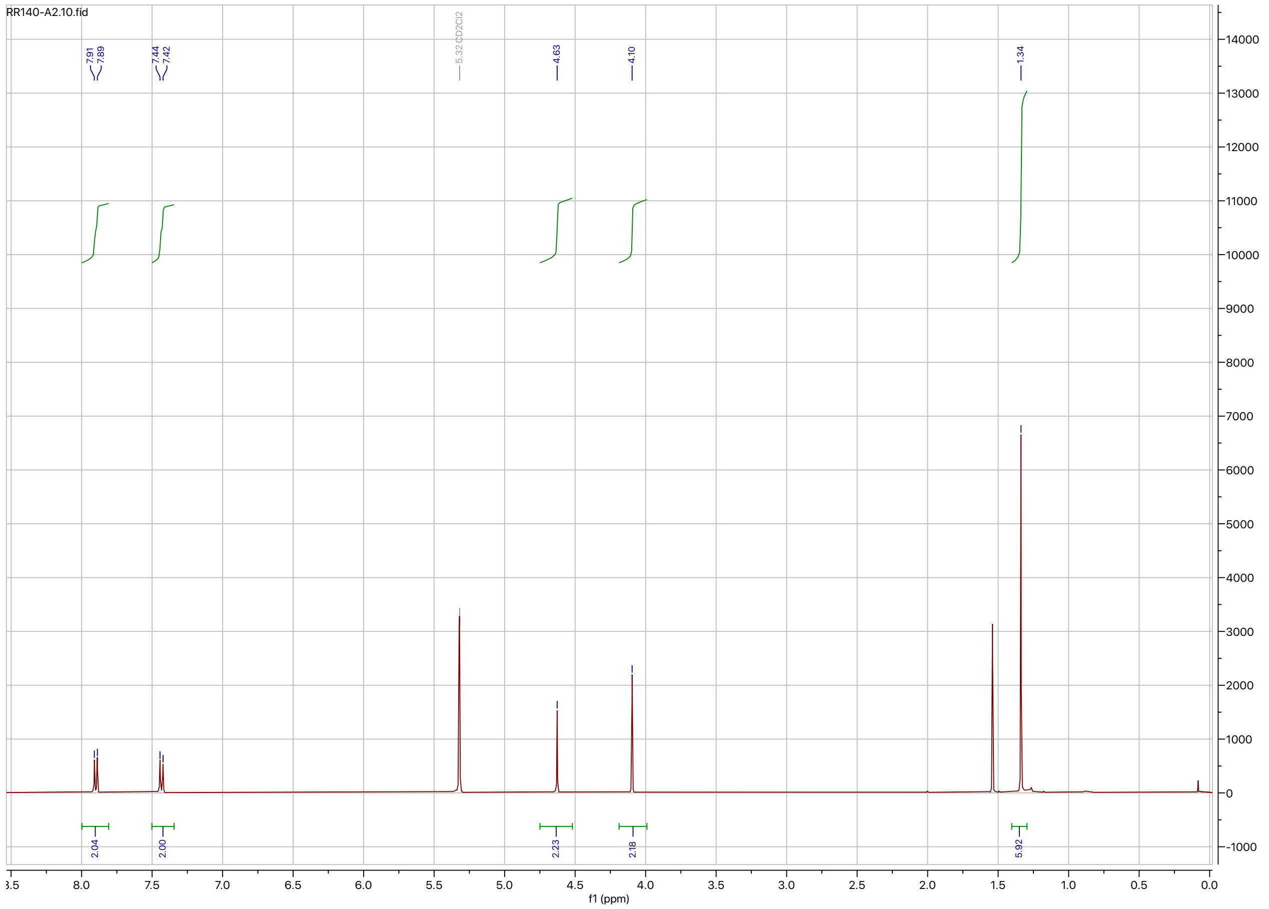
RR140-A2.10.fid
7.91
7.89
7.44
7.42
5.32 CD2Cl2
4.63
4.10
1.34
2.04
2.00
2.23
2.18
5.92
f1 (ppm)

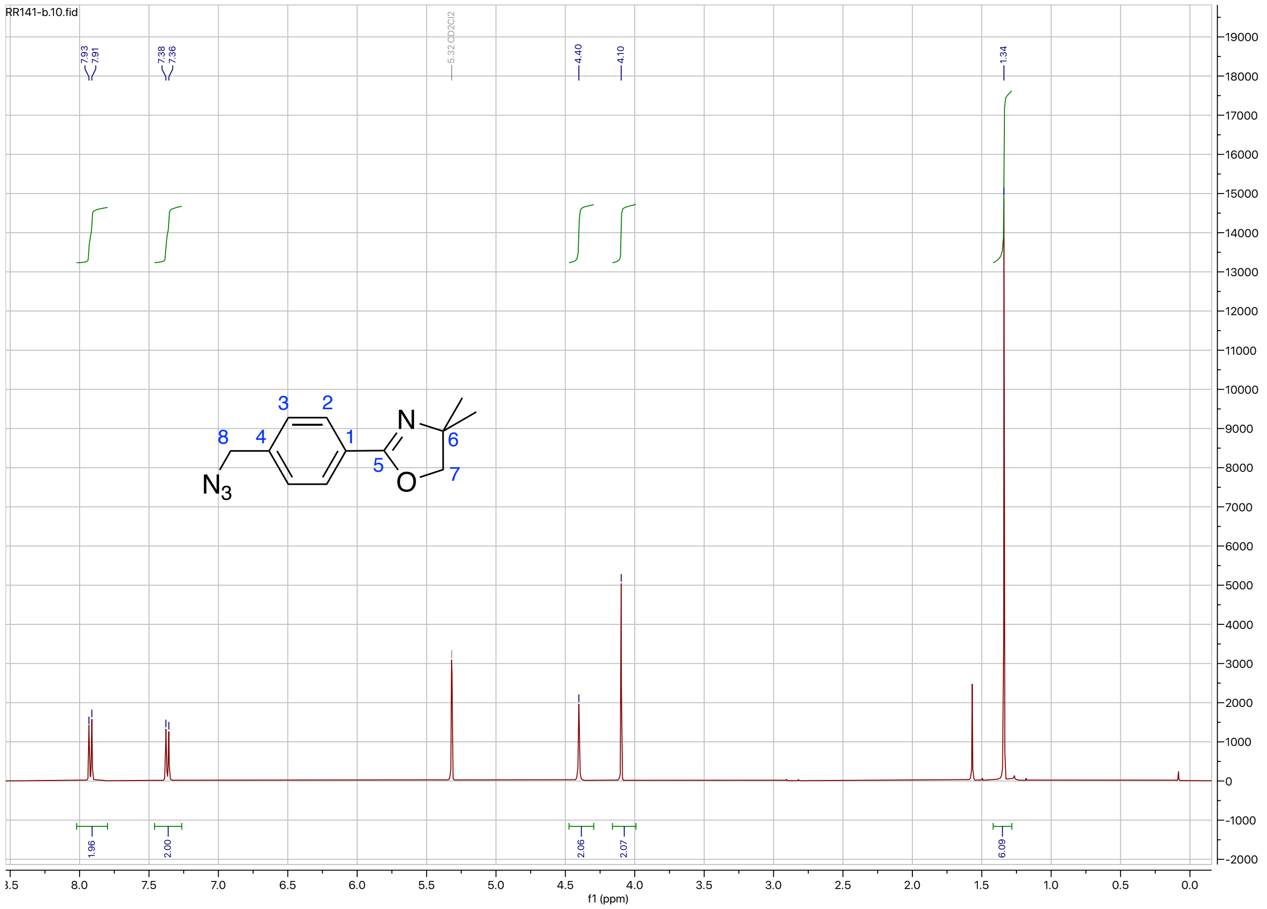
RR141-b.10.fid
7.93
7.91
7.38
7.36
5.32 CD2Cl2
4.40
4.10
1.34
1.96
2.00
2.06
2.07
6.09
N
O
N3
f1 (ppm)

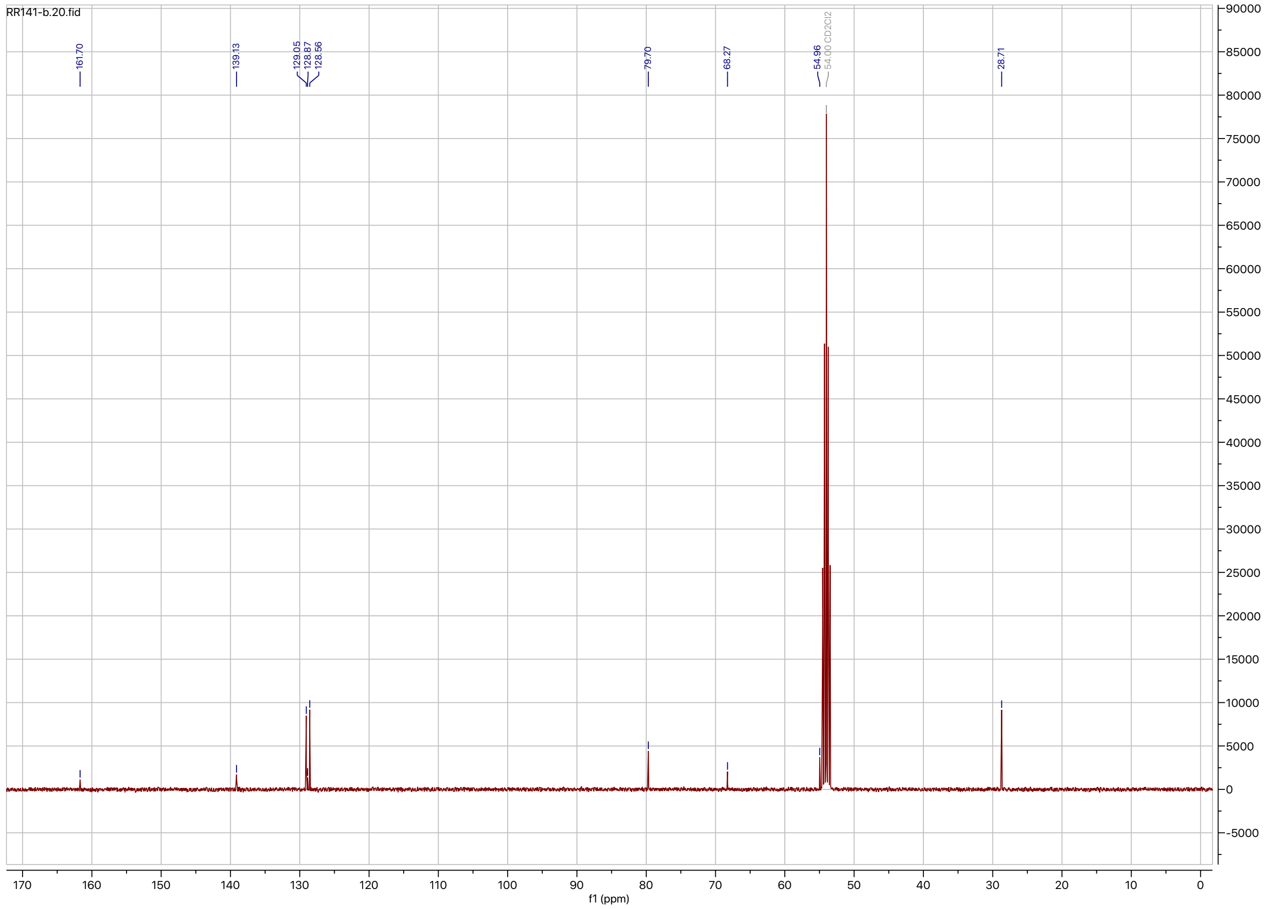
RR141-b.20.fid
161.70
139.13
129.05
128.87
128.56
79.70
68.27
54.96
54.00 CD2Cl2
28.71
f1 (ppm)

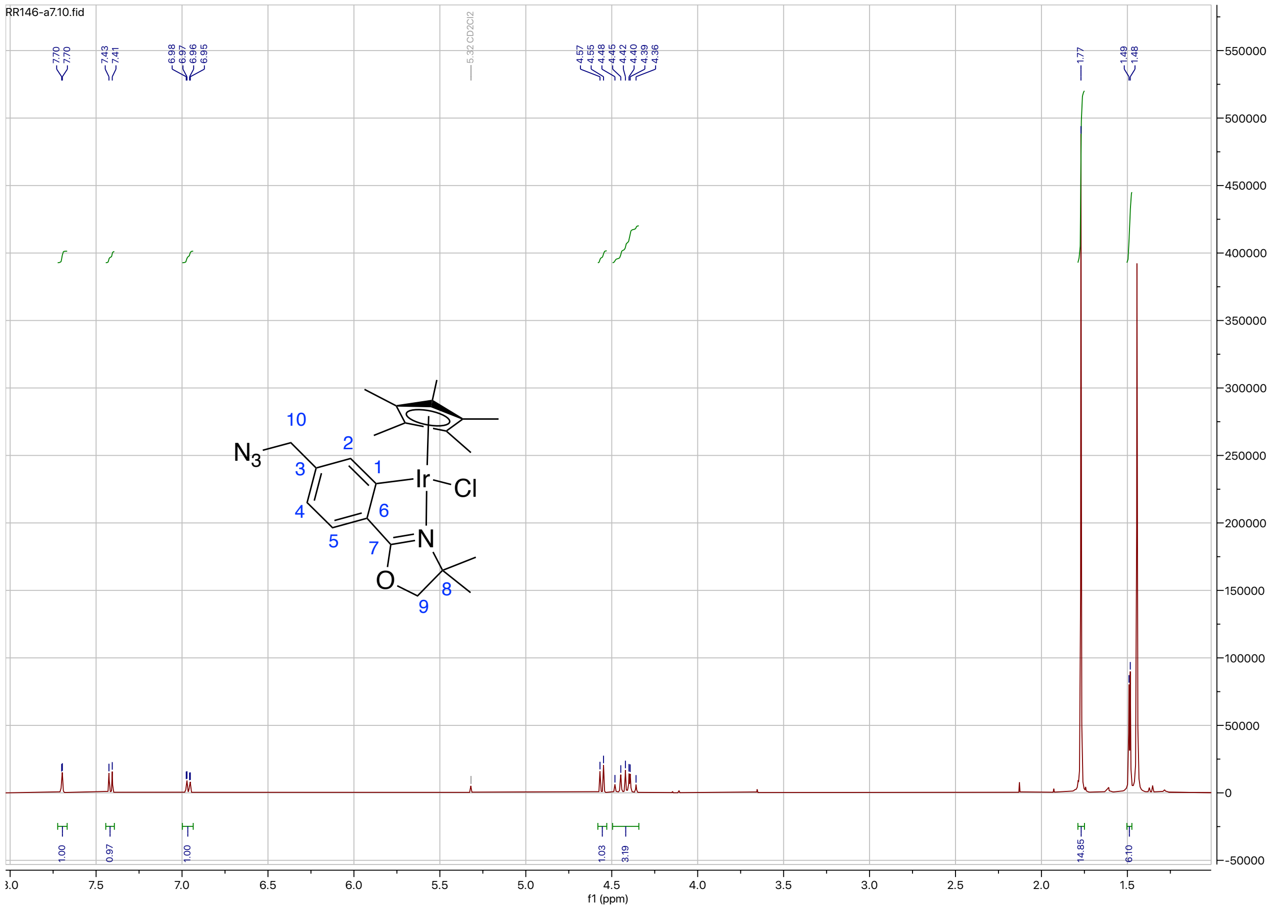
RR146-a7.10.fid
f1 (ppm)
N3
Ir
Cl
N
O

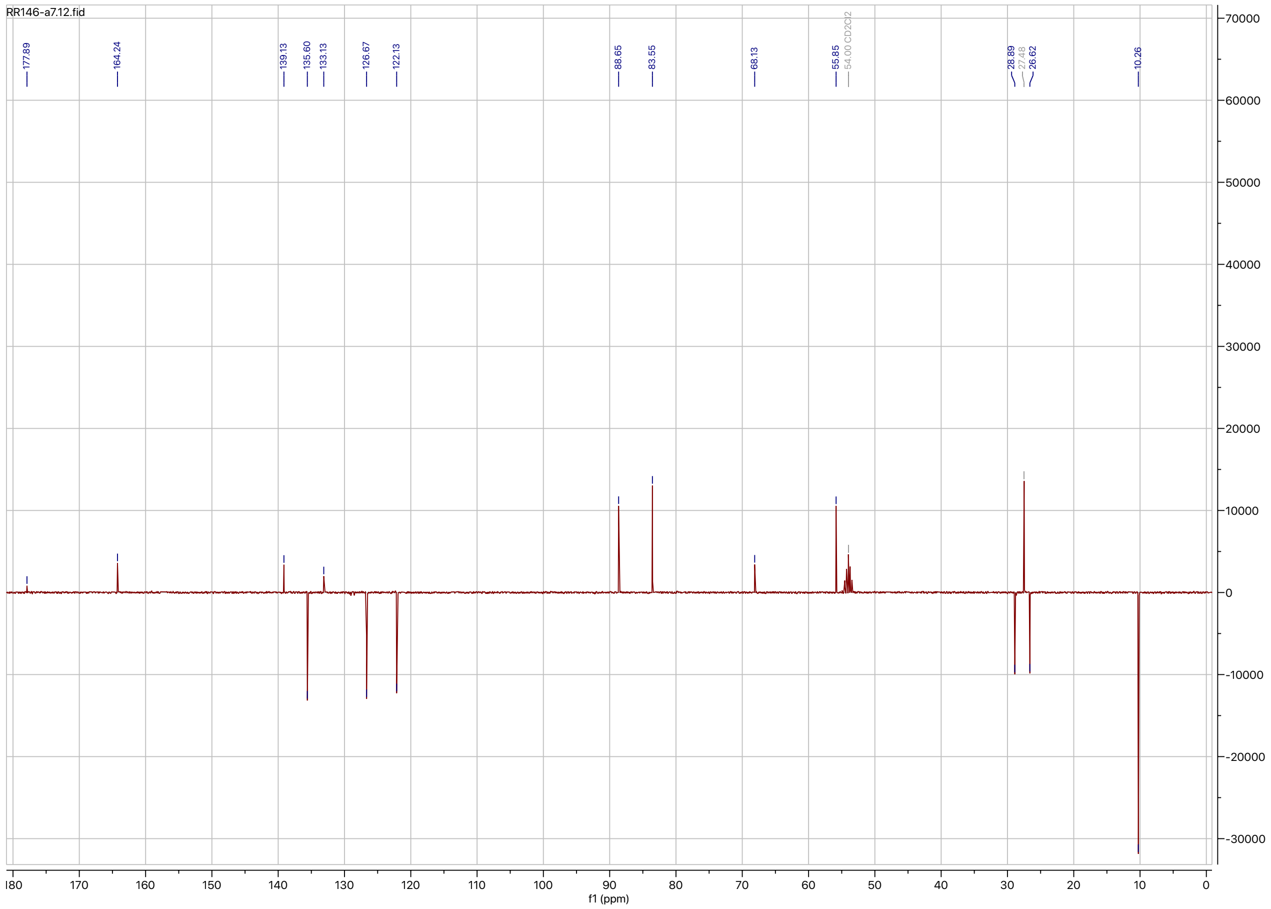
RR146-a7.12.fid
f1 (ppm)
